# seq2ribo: Structure-aware integration of machine learning and simulation to predict ribosome location profiles from RNA sequences

**DOI:** 10.64898/2026.02.08.700508

**Authors:** Gün Kaynar, Carl Kingsford

## Abstract

**Motivation:** Ribosome dynamics are vital in the process of protein expression. Current methods rely on ribosome profiling (Ribo-seq), RNA-seq profiles, and full genomic context. This restricts their use in de novo sequence design, like messenger RNA (mRNA) vaccines. Simulation-only approaches like the Totally Asymmetric Simple Exclusion Process (TASEP) oversimplify translation by focusing solely on codon elongation times.

**Results:** We present `seq2ribo`, a hybrid simulation and machine learning framework that predicts ribosome A-site locations using only an mRNA sequence as input. Our method first employs a novel structure-aware TASEP (sTASEP), which models translation using a comprehensive set of fitted parameters that include codon wait times and structural features, such as local angles, base-pairing, and discrete positional buckets. The ribosome locations generated by sTASEP are then processed by a polisher model, which learns to refine the simulated ribosome distributions. `seq2ribo` provides high-fidelity predictions of ribosome locations across diverse cell types (iPSC, HEK293, LCL, and RPE-1), significantly outperforming baselines. `seq2ribo` is the first method to achieve meaningful positional correlation with observed ribosome profiles from sequence alone, reaching transcript-level Pearson correlations up to 0.920 and within-transcript shape correlations up to 0.186, where all baselines yield near-zero values on these metrics. `seq2ribo` also reduces elementwise error by up to 37.7% relative to the sequence-only Translatomer baseline. By adding a task-specific head, `seq2ribo` achieves Pearson correlations up to 0.732 with experimental translation efficiency (TE) across several cell lines, and up to 0.903 with measured protein expression. By operating from sequence alone, `seq2ribo` provides a new tool for synthetic biology, enabling the rational design and optimization of mRNA sequences without the need for expression-level data or genomic context.

**Availability:** `seq2ribo` is available at https://github.com/Kingsford-Group/seq2ribo.

**Supplementary information:** Supplementary data are available.

## 1 Introduction

The translation of messenger RNA (mRNA) into protein is a fundamental biological process and a primary determinant of protein expression levels in the cell. Translation does not proceed as a uniform, steady-state production line. It is a dynamic system governed by initiation, elongation, and termination (Hershey et al., 2012). Ribosomes move at variable speeds, pause at specific sequences (Yu et al., 2015), and form traffic jams that influence protein output, co-translational folding, and mRNA stability (Jia et al., 2024; Ingolia et al., 2009, 2012). Quantitative models of these ribosome dynamics are therefore central to molecular biology and synthetic biology.

Translation rate and translation efficiency (TE) are key quantitative outcomes of this process (Ingolia et al., 2009; Schwanhäusser et al., 2011; Zarai et al., 2017). They partially emerge from the combined effects of codon usage and mRNA structure. The mRNA folds into structured elements such as hairpins, loops, and pseudoknots (Rouskin et al., 2014). These structures act as physical barriers that slow or stall ribosomes (Caliskan et al., 2014; Bao et al., 2020; Lin et al., 2022). At the same time, the degeneracy of the genetic code means that many different coding sequences can encode the same protein. These synonymous sequences fold into distinct structures and produce very different ribosome traffic patterns and expression levels (Mauro and Chappell, 2014). This makes sequence-level prediction of translation dynamics both challenging and attractive, especially for the rational design of mRNA therapeutics such as vaccines and protein replacement therapies (Pardi et al., 2018; Verbeke et al., 2022; Leppek et al., 2022; Jin et al., 2025). Sequence-only deep learning models have also been developed for related tasks such as translation initiation site prediction (Zhang et al., 2017) and RNA subcellular localization (Yan et al., 2019; Cao et al., 2018).

Ribosome profiling (Ribo-seq) experiments produce codon-level ribosome occupancy profiles by sequencing ribosome-protected mRNA fragments (Ingolia et al., 2012; Spealman et al., 2016; Ingolia et al., 2009; Ahmed et al., 2019; Ingolia et al., 2019; Wang et al., 2016). These data have enabled analysis tools that detect translated open reading frames, normalize ribosome profiles, and quantify translation from existing experiments (Calviello et al., 2016; Xiao et al., 2018; O’Connor et al., 2016). Most computational models use these experimental profiles as input rather than predicting them de novo. Translatomer (He et al., 2024) predicts Ribo-seq signal by integrating genomic sequence with cell-type-specific RNA-seq coverage. It also includes a sequence-only variant, but this simplified model underperforms the full sequence-plus-RNA-seq approach.

Sequence-based TE models such as RiboNN learn to predict TE directly from RNA sequence, using experimental TE labels only during training (Zheng et al., 2025). These approaches demonstrate that sequence carries a strong signal about translational output, but they still require experimental labels for each transcript during model training or rely on additional expression measurements, and they do not produce codon-level ribosome profiles for novel sequences. At the other end of the spectrum, simulation-based approaches such as the totally asymmetric simple exclusion process (TASEP) model ribosome traffic on a one-dimensional lattice (Spitzer, 1970; MacDonald et al., 1968; Zia et al., 2011). Classical implementations treat resistance along the mRNA as a function of codon-specific elongation times and tRNA availability (Dong et al., 2007; von der Haar, 2012) and do not account for mRNA structure, even though structural elements and sequence motifs are known to induce strong ribosome pausing and stalling (Qu et al., 2011; Bao et al., 2020).

We develop `seq2ribo` (Figure 1), a hybrid framework that combines mechanistic simulation with deep learning to predict high-fidelity ribosome dynamics from nucleotide sequence alone. `seq2ribo` consists of two components. First, we introduce sTASEP, a structure-aware totally asymmetric simple exclusion process that models translation with a fitted parameter set that includes codon wait times and additional wait times for local base pairing, structural angle changes, and positional buckets along the coding sequence (CDS). Second, we pass the ribosome distributions that sTASEP generates through a polisher based on the Mamba (Gu and Dao, 2024) structured state space model, which refines the simulated A-site profiles into final ribosome count predictions.

**Figure 1.**
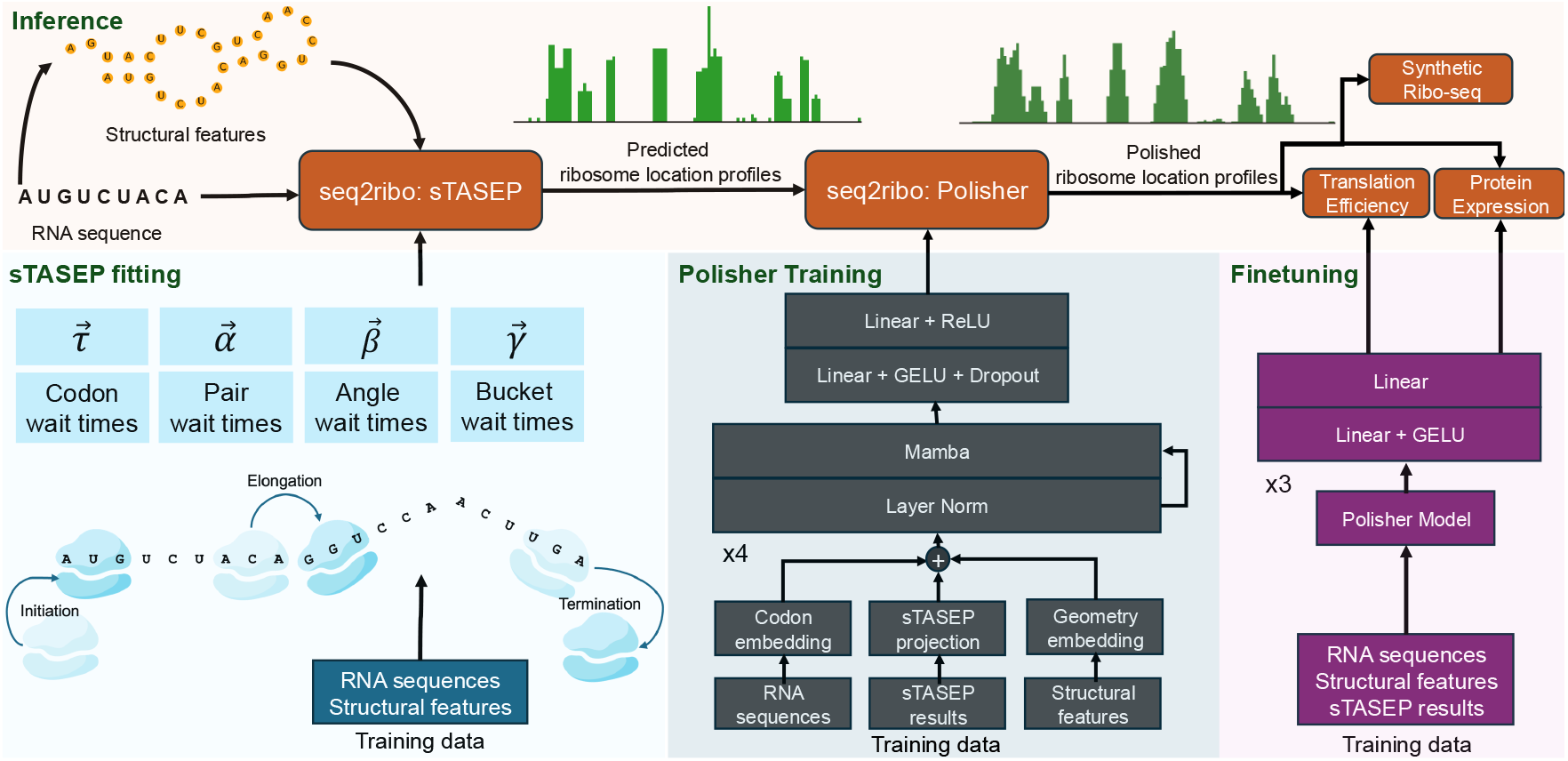
Overview of `seq2ribo` during inference. Given an input RNA sequence, we first derive codon-level structural features and pass the sequence and features into the sTASEP module of `seq2ribo`, which simulates a raw ribosome location profile. The polisher module of `seq2ribo` then combines the codon sequence, structural features, and simulated profile to produce a polished ribosome location profile. We reuse this shared polished profile for two sequence-only downstream tasks. We predict translation efficiency (TE) for human transcripts and predict protein expression, and we can also use the predicted ribosome profiles to generate synthetic ribosome profiling data. The bottom panels summarize how we fit the sTASEP parameters, train the polisher model, and finetune `seq2ribo` for the downstream regression tasks.

We demonstrate the generalizability of our framework by training and evaluating `seq2ribo` on four distinct cell lines: induced pluripotent stem cells (iPSC), human embryonic kidney 293 (HEK293), lymphoblastoid cell lines (LCL), and hTERT-immortalized retinal pigment epithelial cells (RPE-1). We show that sTASEP consistently improves on a classical TASEP baseline, and that the full `seq2ribo` pipeline substantially outperforms a sequence-only Translatomer baseline. No existing method achieves meaningful positional agreement with observed ribosome profiles from sequence alone. `seq2ribo` changes this. It attains transcript-level Pearson correlations between 0.657 and 0.920 and within-transcript shape correlations between 0.054 and 0.186 across all four cell lines, while every baseline remains at or below zero on these metrics. `seq2ribo` additionally reduces elementwise mean absolute error (MAE) by 30.3% to 37.7% in three of four cell lines relative to Translatomer. We further evaluate `seq2ribo` as a sequence-only predictor for downstream tasks. For translation efficiency (TE), we outperform the sequence-based RiboNN baseline in the CDS-only setting across all four cell lines, increasing Pearson correlation from 0.529 to 0.688 on the mean label set and up to 0.697 on cell-type-specific sets. When UTR information is included, `seq2ribo` achieves the highest correlation in three of four tasks with Pearson *r* up to 0.732. For protein expression, all four cell-line polishers achieve Pearson correlations between 0.830 and 0.903 on an external expression dataset after finetuning. Together, these results show that predicted ribosome count profiles from `seq2ribo` serve as strong features for TE and expression. Moreover, `seq2ribo` functions as a synthetic Ribo-seq data generator, enabling the in silico profiling of arbitrary sequences and providing a practical tool for de novo mRNA design.

## 2 Methods

### 2.1 Data

We obtain global aggregate A-site ribosome counts for iPSC (Chen et al., 2020), LCL (Cenik et al., 2015), HEK293 (Wagner et al., 2020), and RPE-1 (Haas et al., 2020) cell lines from the GWIPS-viz Ribo-seq resource (Michel et al., 2014). We restrict all analyses to coding sequences and apply the preprocessing and filtering pipeline described in detail in Section 1.1 of the Supplementary Text. For each retained transcript, we have a codon sequence and a matched A-site ribosome count sequence of the same length. For each CDS, we also construct three codon-level structural feature sequences that have the same length as the codon sequence. We generate these structural features using ViennaRNA (Lorenz et al., 2011) and describe the folding and feature construction procedure in Section 1.2 of the Supplementary Text. To verify that the test splits assess generalization to novel sequences rather than memorization of near-neighbors, we perform a train-test similarity analysis in Section 1.16 of the Supplementary Text.

For translation efficiency (TE), we define four distinct prediction tasks based on experimental values from Zheng et al. (2025). We use three cell-line-specific TE measurements (HEK293, RPE-1, and LCL) and a mean TE measurement. We match these labels to the coding sequences in our dataset to create matched datasets of transcripts that possess codon sequence, A-site ribosome counts, and the corresponding experimental TE label. For each task, we split the transcripts into training (80%), validation (10%), and test (10%) sets using stratification by the total A-site ribosome count per transcript. The exact transcript counts for each task are provided in Section 1.1 of the Supplementary Text.

For experimental protein expression, we use a protein (mRFP) expression dataset of Nieuwkoop et al. (2023). This dataset contains 1,459 coding sequences with measured protein production levels and a predefined split into 1,021 training sequences, 219 validation sequences, and 219 test sequences (Li et al., 2024). For each mRFP codon sequence, we compute the same codon-level pair, angle, and bucket features and retain the codon sequence, structural feature sequences, and ground truth protein expression label.

### 2.2 Problem formulation

The inputs and outputs share the same structure across cell lines (iPSC, HEK293, LCL, RPE-1). We define the notation for a generic dataset *D*. Let 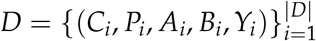 denote the full dataset for the given cell line, where |*D*| is the sample size.

We partition *D* into three disjoint subsets: the training set *D*_train_, the validation set *D*_val_, and the test set *D*_test_. Each sample *i* corresponds to a coding sequence (CDS) with variable length *N*_*i*_ codons. We denote the codon sequence by

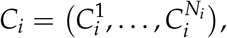

where 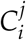 is the *j*-th codon in the CDS of sample *i*. For sample *i*, we define three codon-level structural feature sequences. We denote the pair count sequence by

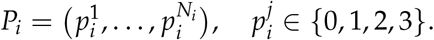

The value 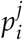 records how many nucleotides in codon 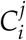 participate in base pairing in the predicted RNA structure. We denote the angle sequence by

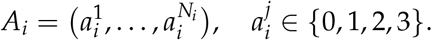

The value 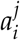 quantifies the change in local backbone angle at codon 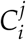 relative to the previous codon in the predicted two-dimensional structure. We discretize continuous angle changes into four bins that represent no change, small change, medium change, and large change. We derive these bins from the distribution of angle changes in the training set and describe this procedure in Section 1.3 of the Supplementary Text. We denote the bucket sequence by

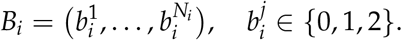

The value 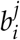 encodes the relative position of codon 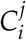 along the CDS. Codons in the first third of the sequence receive bucket value 0. Codons in the middle third receive bucket value 1. Codons in the final third receive bucket value 2. We provide the rationale for this three-bucket design in Section 1.2 of the Supplementary Text. For each sample *i*, we observe A-site ribosome counts at each codon position, where each count is an integer in {0, 1, 2, …}. We denote the target count sequence by 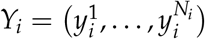, where 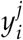 is the observed A-site ribosome count at codon 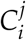. We use an integer *K* ≥ 1 to denote the number of independent simulation runs that we generate for each sample. We implement a stochastic simulator

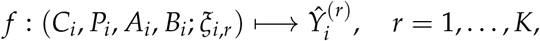

that maps the codon sequence and all structural features to a sequence of simulated A-site counts, where *ξ*_*i,r*_ denotes simulator randomness for run *r*. For each sample *i*, we obtain *K* simulator outputs

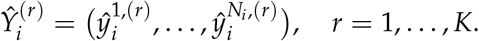

Each 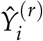 is the output of one independent simulation run, where 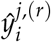 represents the simulated A-site count at the *j*-th codon of sample *i* in run *r*. We report the value of *K* used in each experiment in the corresponding sections. We collect the *K* simulated sequences for sample *i* into the set

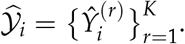

We train a polisher that refines a single simulator output at a time. We learn a function

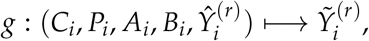

that refines the simulator outputs. The polisher receives the same codon and structural features, and produces polished predictions

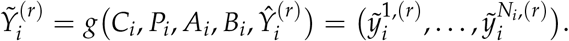

For polisher training, we treat each pair (*i, r*) as an independent example.

We evaluate models with nine metrics that summarize agreement between predicted and observed A-site counts at different levels of aggregation. We treat two Pearson correlation metrics as primary indicators of prediction quality. Tx-level *r* computes the total observed and predicted counts for each transcript and takes the Pearson correlation of these transcript-level sums. Shape *r* computes the Pearson correlation between predicted and observed counts within each transcript and averages across transcripts; it isolates whether the model captures the positional distribution of ribosomes along each coding sequence. We also report Elemwise *r*, the Pearson correlation across all codon positions in all transcripts, and six MAE metrics: elemwiseMAE (average absolute error across all positions), pertranscriptMAE (per-transcript absolute error averaged across transcripts), codonMAE (error aggregated by codon identity), pairMAE (by base pairing category), angleMAE (by discretized backbone angle change), and bucketMAE (across the first, middle, and final thirds of each CDS). Because we compute both pair and angle errors over four discrete groups, pairMAE and angleMAE can occasionally take identical values. We describe the calculation of each metric in Section 1.4 of the Supplementary Text. Higher values are better for the three correlation metrics, and lower values are better for the six MAE metrics.

### 2.3 The Structure-Aware TASEP (sTASEP)

We implement the simulator *f* as a structure-aware totally asymmetric simple exclusion process (sTASEP). For each transcript *i*, we simulate ribosomes that move along a one-dimensional lattice of codon positions *j* = 1, …, *N*_*i*_. Each ribosome occupies a single codon index and excludes other ribosomes within three codons upstream and three codons downstream along the mRNA (Brar and Weissman, 2015). At each simulation step, we sample three types of stochastic events. **Initiation** introduces a new ribosome at the first codon if codon positions 1 through 4 are free. **Elongation** moves a ribosome from position *j* to *j* + 1 only if the downstream exclusion zone is empty and proceeds with an elongation rate

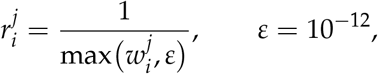

where 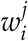 is a codon-specific wait time that combines sequence and structural effects at that position,

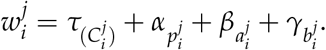

Here 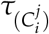 is the wait time for the non-stop codon at position *j*, 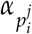 is the wait time for the pair category 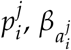 is the wait time for the angle category 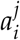 and 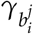 is the wait time for the bucket category 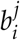. We maintain one parameter for each of the 61 non-stop codons, one parameter for each of the four pair categories, one parameter for each of the four angle categories, and one parameter for each of the three bucket categories. **Termination** removes a ribosome when it reaches the stop codon and records a termination event.

We fit a distinct set of wait time parameters for each cell line. Because the sTASEP uses a single initiation rate shared across all transcripts, the simulator cannot capture transcript-level differences in ribosome load; it can only shape the positional distribution of ribosomes along each coding sequence. We therefore fit the wait time parameters using the scaled MAE metrics, which normalize predicted counts to match the observed ribosome load on each transcript before computing errors. This ensures that the parameter updates are driven by discrepancies in the positional profile rather than by differences in overall transcript abundance. We describe the full simulation schedule, parameter initialization, and update algorithm in Section 1.5 of the Supplementary Text. As an alternative initialization strategy, we also derive analytical first estimates of the wait time parameters directly from observed ribosome occupancy, as described in Section 1.9 of the Supplementary Text.

### 2.4 Structured state space polisher

We implement the polisher *g* as a structured state space model based on Mamba (Gu and Dao, 2024). The model refines the simulated A-site counts produced by sTASEP and outputs polished codon-level predictions. For a transcript *i* with length *N*_*i*_, we construct five input channels. We embed codon identity, the pair sequence, the angle sequence, the bucket sequence, and the simulated A-site counts into a shared latent space and sum these embeddings position-wise to obtain an initial representation 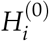 of length *N*_*i*_. The core of the polisher consists of stacked Mamba blocks. Each block applies layer normalization (LN), a Mamba state space layer, and a residual connection,

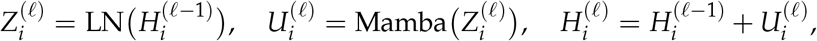

for block index *ℓ*. The Mamba layers operate along the sequence dimension and share parameters across positions. After the final Mamba block, we apply a position-wise feed-forward head and a Softplus nonlinearity to map each position to a single non-negative scalar. We train a separate polisher *g* for each cell line using the corresponding training set *D*_train_. Each training example consists of (*C*_*i*_, *P*_*i*_, *A*_*i*_, *B*_*i*_, 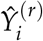) as inputs and the observed A-site counts *Y*_*i*_ as targets. We minimize a Poisson negative log-likelihood loss and select the best checkpoint by validation loss. We provide embedding dimensions, the exact number of layers, and all optimization hyperparameters in Section 1.6 of the Supplementary Text. We use the resulting polisher together with the fitted sTASEP simulator as the `seq2ribo` pipeline in all experiments. We provide a detailed analysis of the computational hardware, theoretical complexity, and runtime estimates for both simulation and training in Section 2 of the Supplementary Text.

### 2.5 Downstream tasks

To demonstrate that the ribosome count profiles produced by `seq2ribo` form a good basis for downstream tasks, we place two regression heads on top of the polisher output. One head predicts TE, and one head predicts protein expression. In both cases, the polisher first produces a codon-level count profile for each transcript and a task-specific head then aggregates this profile to a single scalar prediction.

#### 2.5.1 Translation efficiency prediction

We address the four TE tasks described in Section 2.2 by training a specific prediction head for each task. For each transcript, we treat the polished codon-level profile 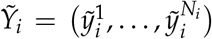 as a sequence of features. We apply an element-wise feed-forward network independently at each codon position, then apply masked mean pooling over positions to obtain a fixed-length transcript representation, and pass this representation through a final linear layer that outputs a single scalar prediction. We train each head with a mean squared error (MSE) loss between predicted and experimental TE values. We average predictions over multiple simulated and polished profiles per transcript to reduce variance. To enable a fair comparison with RiboNN, which ingests the full transcript including 5^′^ and 3^′^ UTRs, we also train a UTR-aware variant of the `seq2ribo` TE head that additionally processes the UTR nucleotide sequences. We describe both head architectures, the number of simulations, and all training details in Section 1.7.1 of the Supplementary Text.

#### 2.5.2 Protein expression prediction

For the mRFP protein expression dataset, we apply the same strategy. We run sTASEP on each sequence, polish the simulated A-site counts to obtain a codon-level profile, apply a per-position feed-forward network followed by masked mean pooling to get a fixed-length transcript representation, and pass this representation through a final linear layer that outputs a single protein expression prediction. We finetune the full model on the mRFP training split, use the validation split for early stopping, and evaluate on the test split. We again average predictions over many simulated and polished profiles per sequence. We provide the exact simulation counts and training setup in Section 1.7.2 of the Supplementary Text.

### 2.6 Baseline models

We benchmark both components of `seq2ribo` against internal and external baselines. We first compare the sTASEP simulator to a classical codon-level TASEP model that fits only codon-specific wait times. We fit separate TASEP baselines for each cell line. We then assess the benefit of the Mamba-based polisher by replacing the structured state space backbone with a transformer-based architecture that uses the same input embeddings and a comparable number of parameters. We also train a variant of the Mamba polisher that optimizes pertranscriptMAE. We describe the classical TASEP formulation, the transformer architecture, and the pertranscriptMAE-only polisher ablation in Sections 1.8, 1.12, and 1.13 of the Supplementary Text.

As an external baseline for ribosome profile prediction, we use Translatomer (He et al., 2024). Translatomer was originally designed for variant interpretation using genomic DNA in a fixed-width window together with RNA-seq coverage to predict ribosome footprint density. To adapt it to our sequence-only, CDS-level setting, we train a variant that uses the same genomic DNA window input but omits RNA-seq coverage from both the architecture and the inputs, and instead predicts A-site ribosome counts. We train four separate sequence-only Translatomer models, one per cell line, and evaluate each on the corresponding test set. We include Translatomer because it is the most closely related prior method that predicts ribosome occupancy from sequence, though the comparison should be interpreted with the caveat that it is being evaluated outside its intended use case. Training details appear in Section 1.10 of the Supplementary Text.

For TE prediction, we compare the `seq2ribo` TE heads with RiboNN (Zheng et al., 2025). We train and evaluate RiboNN under two input settings. In the CDS-only setting, we train RiboNN on coding sequences only, with UTR lengths set to zero, matching the information available to the `seq2ribo` CDS-only head. In the UTR+CDS setting, we train RiboNN on the full transcript including both 5^′^ and 3^′^ UTRs, matching the information available to the `seq2ribo` UTR-aware head. For each input setting, we train RiboNN in two configurations: individually, with a separate model for each of the four TE tasks, and as a multitask model that jointly predicts all four cell-line TE values with shared encoder weights and task-specific output heads. All RiboNN models use the same train, validation, and test splits and the same labels that we use for the `seq2ribo` TE heads so that all models see the same data. We give architecture and training details for RiboNN in Section 1.11 of the Supplementary Text.

## 3 Results

### 3.1 seq2ribo

We fit kinetic parameters for both the classical TASEP model and our sTASEP simulator separately for each cell line dataset. We evaluate both simulators with nine metrics: the two primary correlation metrics (Tx-level *r* and Shape *r*), Elemwise *r*, and six MAE metrics (pertranscriptMAE, elemwiseMAE, codonMAE, pairMAE, angleMAE, and bucketMAE). Figure 2 shows training dynamics for the iPSC dataset. Training dynamics for HEK293, LCL, and RPE-1 appear in Supplementary Figures S24–S37. Table 1 reports test set metrics for TASEP, sTASEP, the sequence-only Translatomer baseline, and `seq2ribo` for each cell line. Note that the six MAE metrics in Table 1 are computed after per-transcript scaling to isolate positional profile errors from differences in overall count magnitude; unscaled metrics appear in Supplementary Table S12.

**Table 1:** Test set metrics across cell lines. The six MAE metrics are computed after per-transcript scaling so that predicted and observed ribosome loads match (lower is better). The three Pearson correlation metrics are computed on unscaled predictions (higher is better). Bold indicates the best performance within each cell line and metric.

| Cell line | Metric | TASEP | sTASEP | Translat. (seq.) | seq2ribo |
| --- | --- | --- | --- | --- | --- |
| iPSC | Tx-level $r$ | 0.200 | 0.210 | 0.051 | **0.920** |
| | Shape $r$ | -0.015 | 0.039 | 0.001 | **0.162** |
| | Elemwise $r$ | 0.056 | 0.078 | -0.002 | **0.598** |
|  | pertranscriptMAE | 1.339 | **1.308** | 1.532 | 1.386 |
|  | elemwiseMAE | 4.895 | 4.561 | 6.638 | **4.135** |
|  | codonMAE | 40 894 | **3 725** | 5 916 | 12 028 |
|  | pairMAE | 40 181 | **15 120** | 94 648 | 44 153 |
|  | angleMAE | 16 182 | **12 608** | 94 648 | 85 032 |
|  | bucketMAE | 43 280 | **27 123** | 311 640 | 287 231 |
| Cell line | Metric | TASEP | sTASEP | Translat. (seq.) | seq2ribo |
| HEK293 | Tx-level $r$ | 0.006 | 0.013 | 0.023 | **0.657** |
| | Shape $r$ | 0.002 | 0.007 | 0.000 | **0.054** |
| | Elemwise $r$ | -0.004 | -0.001 | 0.000 | **0.396** |
|  | pertranscriptMAE | 1.811 | 1.807 | **1.001** | 1.909 |
|  | elemwiseMAE | 0.225 | 0.222 | **0.150** | 0.247 |
|  | codonMAE | 244 | **183** | 4 859 | 3 627 |
|  | pairMAE | **300** | 318 | 77 744 | 15 164 |
|  | angleMAE | 431 | **259** | 77 744 | 23 899 |
|  | bucketMAE | 7 403 | **6 685** | 103 658 | 66 934 |
| Cell line | Metric | TASEP | sTASEP | Translat. (seq.) | seq2ribo |
| LCL | Tx-level $r$ | 0.079 | 0.101 | 0.055 | **0.901** |
| | Shape $r$ | -0.012 | 0.046 | 0.000 | **0.186** |
| | Elemwise $r$ | -0.006 | 0.014 | 0.000 | **0.612** |
|  | pertranscriptMAE | 1.642 | 1.518 | 1.599 | **1.471** |
|  | elemwiseMAE | 32.927 | 28.956 | 32.164 | **22.418** |
|  | codonMAE | 800 628 | 35 579 | **33 724** | 38 902 |
|  | pairMAE | 298 006 | 248 121 | 539 576 | **187 622** |
|  | angleMAE | 119 977 | 155 088 | 562 607 | **84 351** |
|  | bucketMAE | 464 613 | **299 512** | 789 398 | 920 437 |
| Cell line | Metric | TASEP | sTASEP | Translat. (seq.) | seq2ribo |
| RPE-1 | Tx-level $r$ | 0.137 | 0.130 | 0.035 | **0.771** |
| | Shape $r$ | -0.005 | 0.041 | 0.003 | **0.139** |
| | Elemwise $r$ | 0.003 | 0.017 | 0.003 | **0.610** |
|  | pertranscriptMAE | 1.411 | **1.369** | 1.590 | 1.503 |
|  | elemwiseMAE | 3.750 | 3.506 | 5.100 | **3.362** |
|  | codonMAE | 23 034 | **2 383** | 4 740 | 12 404 |
|  | pairMAE | 27 098 | **6 858** | 75 832 | 24 184 |
|  | angleMAE | 17 323 | **4 286** | 76 336 | 15 336 |
|  | bucketMAE | **15 733** | 22 854 | 497 259 | 356 474 |

**Figure 2.**
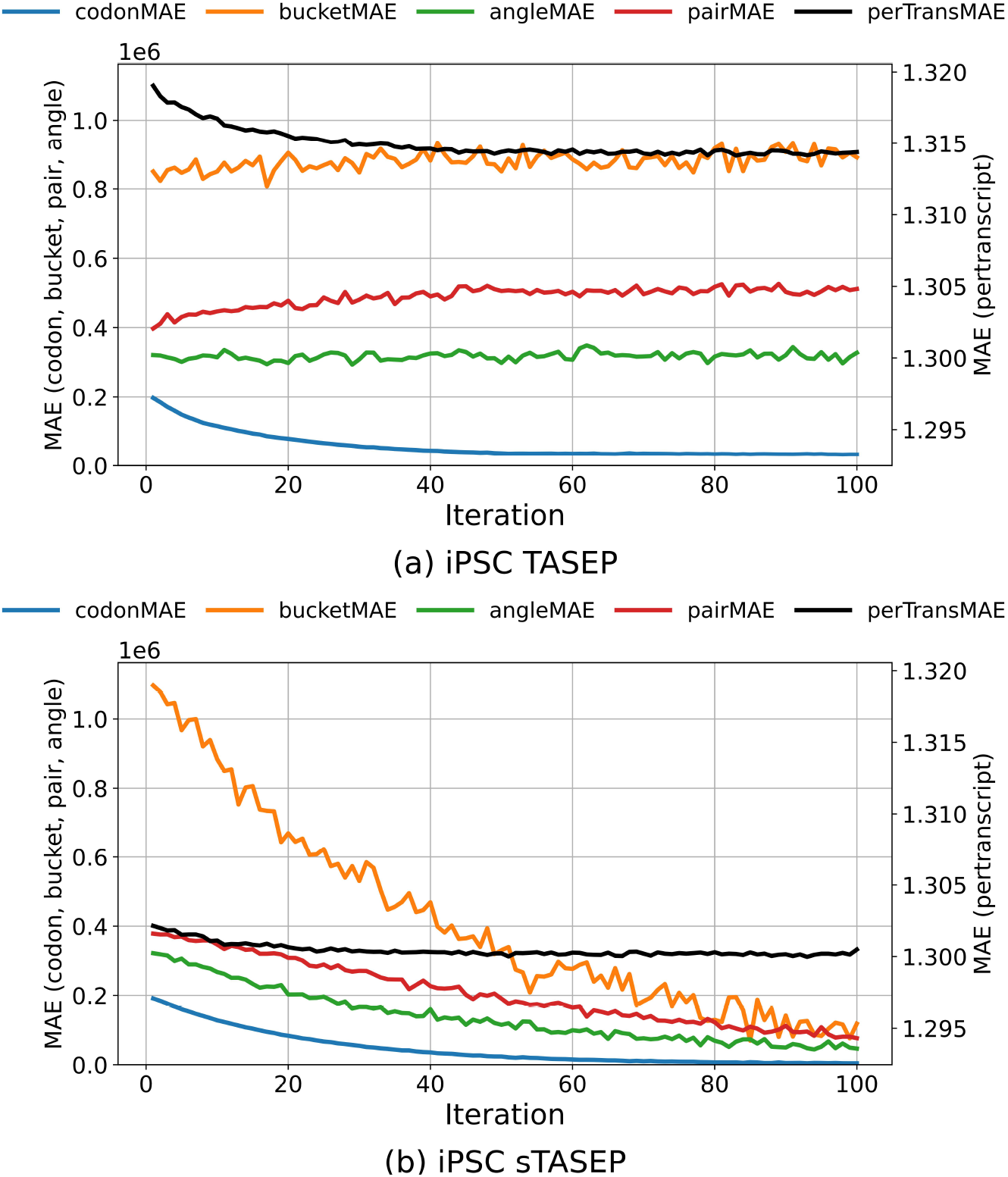
Loss convergence for classical TASEP and sTASEP fits on the iPSC dataset.

Across all cell lines, sTASEP improves simulator accuracy over classical TASEP. sTASEP raises Shape *r* in every cell line from negative or near-zero to small positive values, lowers both pertranscriptMAE and elemwiseMAE in every dataset, and improves Elemwise *r* in all four cell lines and Tx-level *r* in three of four.

The gains are even more pronounced in structural metrics, where sTASEP reduces codonMAE by 89.7%–95.6% in iPSC, LCL, and RPE-1. While classical TASEP retains an advantage on isolated structural metrics in individual cell lines, sTASEP yields improvements across the majority of structural metrics in every dataset. These results demonstrate that adding explicit structural terms to the simulator improves agreement with observed ribosome profiles across diverse experimental settings, and we therefore adopt sTASEP as the simulator *f* for each cell line. The fitted sTASEP parameters are biologically interpretable, with codon wait times positively correlated across iPSC, LCL, and RPE-1 (Spearman *ρ* = 0.268–0.421) and correlated with the Codon Stabilization Coefficient (CSC) (Presnyak et al., 2015) in all four cell lines (*ρ* = 0.308–0.418) (Section 1.20 of the Supplementary Text).

Using the fitted sTASEP simulator, we generate 32 independent simulation runs per transcript and treat each run as a separate training example for the Mamba-based polisher *g* for each cell line. We train the polisher with the Poisson negative log-likelihood loss described in Section 1.6 of the Supplementary Text, and we select checkpoints using the validation split for each cell line. Training and validation loss curves for the polisher across all four cell lines appear in Supplementary Figures S38–S45. We report full pipeline metrics for each cell line in Table 1, and we use the resulting simulator and polisher as the `seq2ribo` pipeline in subsequent experiments.

### 3.2 seq2ribo benchmark against baselines

We benchmark the complete `seq2ribo` pipeline against Translatomer (sequence only) across four cell lines. As shown in Table 1, `seq2ribo` achieves the highest Shape *r* and Tx-level *r* in every cell line. Shape *r* reaches 0.054–0.186 across cell lines, where all baselines produce near-zero or negative values, indicating that `seq2ribo` is the only method that reliably captures the positional profile of ribosome occupancy within individual transcripts. The gains in Tx-level *r* are even more striking: `seq2ribo` achieves 0.657–0.920, far exceeding all baselines and demonstrating strong agreement in transcript-level ribosome load.

`seq2ribo` achieves the lowest elemwiseMAE in three of four cell lines, with reductions of 30.3%– 37.7% relative to Translatomer. In HEK293, Translatomer attains lower MAE by predicting all-zero counts, which minimizes absolute error but captures no positional variation, as reflected in its zero correlations on this dataset.

We also observe a trade-off between transcript-level fidelity and fine-scale structural fit. While `seq2ribo` achieves the best performance on all three correlation metrics, the intermediate sTASEP simulator often achieves lower error on structural metrics such as codonMAE, pairMAE, and angleMAE, suggesting the polisher prioritizes positional correlation over fine-scale structural features. Despite this, `seq2ribo` still improves over Translatomer on structural metrics in the majority of cases. Bootstrap 95% confidence intervals confirm the statistical reliability of these results, with narrow bounds on all key metrics (e.g., iPSC Tx-level *r* = 0.920 [0.883, 0.946]) and negligible run-to-run variance across *K* = 32 simulations (Sections 1.22 and 1.23 of the Supplementary Text). Cross-cell-line evaluation shows that transferred models retain substantial predictive power (e.g., iPSC polisher achieves Tx-level *r* = 0.576 on HEK293), indicating that the learned representations generalize across cellular contexts (Section 1.21 of the Supplementary Text).

### 3.3 Translation efficiency prediction

We evaluate `seq2ribo` on the TE prediction tasks defined in Methods, covering both mean TE and cell-type-specific efficiency for HEK293, LCL, and RPE-1. We compare against Translatomer (sequence-only) and RiboNN trained in both individually per cell line and multitask configurations, using the same train, validation, and test splits for all models.

As shown in Table 2, finetuned `seq2ribo` achieves the best Pearson correlation in the CDS-only setting across all four tasks, outperforming the best RiboNN variant by a wide margin. Translatomer shows the weakest performance across all tasks. When UTR information is added, both RiboNN and `seq2ribo` improve, with `seq2ribo` achieving the highest correlation in three of four tasks. In RPE-1 with CDS+UTR input, RiboNN outperforms `seq2ribo` (*r* = 0.667 vs. 0.624), suggesting that UTR features carry cell-type-specific information that the current `seq2ribo` architecture does not fully exploit. As shown in Figure 3, the finetuned `seq2ribo` model displays a strong monotonic trend and a narrow spread for the mean TE label set. We observe the same trend for LCL, HEK293, and RPE-1 in Supplementary Figures S47–S49. See Supplementary Figures S50–S57 for scatter plots of TE versus Translatomer and RiboNN predictions.

**Table 2:**
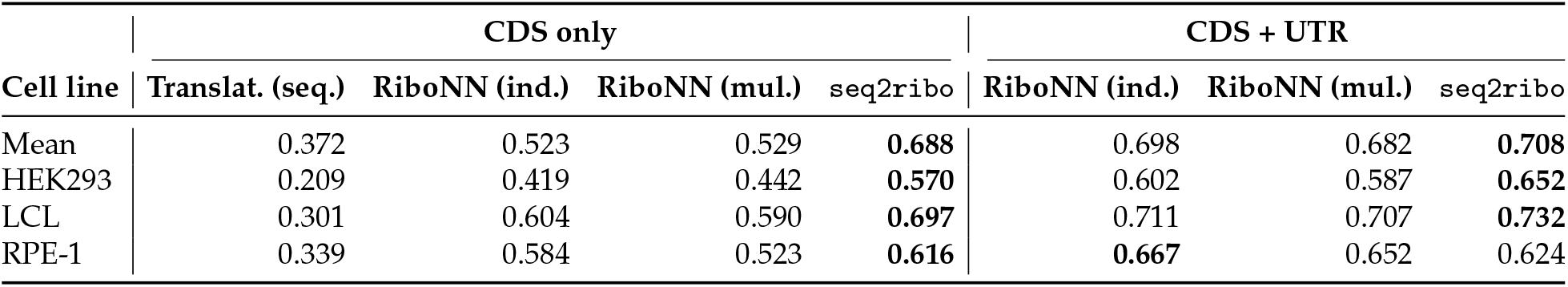
Translation efficiency prediction results on the test set. We report Pearson correlation *r*. Bold indicates the best performance within each input setting and cell line. Indiv. = independently trained per cell line; Multi. = multitask learning across cell lines.

**Figure 3.**
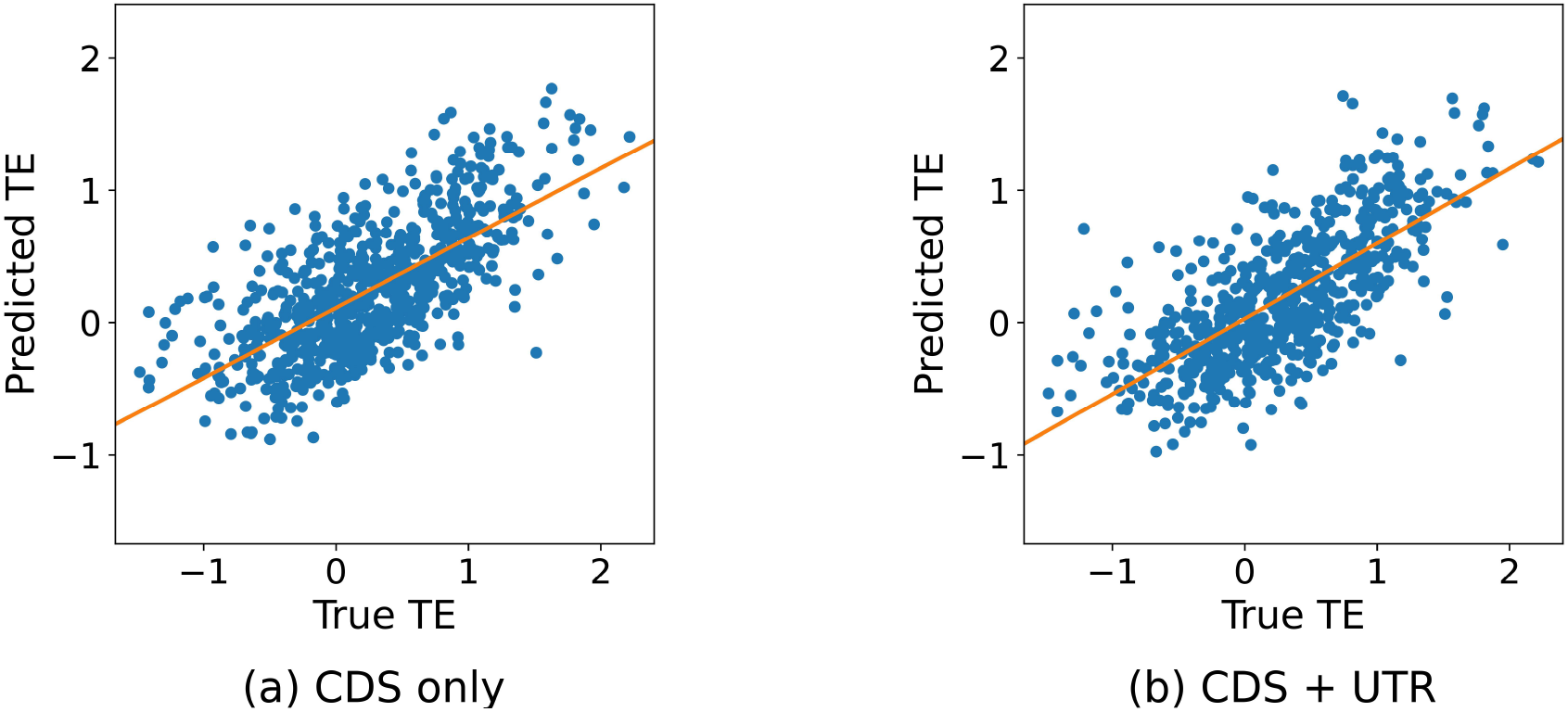
Correlation of finetuned `seq2ribo` predictions with experimental TE on the Mean TE test set. (a) CDS-only input. (b) CDS + UTR input.

These results show that the structured ribosome count profiles generated by `seq2ribo` provide a richer representation than direct sequence-to-TE models, enabling a simple task-specific head to reach higher accuracy. Stratified evaluation across ten TE-level bins shows that `seq2ribo` maintains stable prediction quality regardless of translational activity, with Shape *r* remaining consistently positive across all bins and cell lines (Section 1.18 of the Supplementary Text). Perturbation analysis confirms that TE predictions are robust to moderate noise, dropout, and scaling of the input profiles (Section 1.17 of the Supplementary Text). We also report correlations between raw sTASEP and polisher total ribosome load and experimental TE in Supplementary Table S15 with corresponding scatter plots in Supplementary Figures S46–S49.

### 3.4 Protein expression prediction

We evaluate `seq2ribo` on a protein expression task using an mRFP expression dataset (Nieuwkoop et al., 2023). Because this dataset provides a single set of expression measurements without cell-type-specific labels, we use it to test whether polishers trained on different cell-line Ribo-seq data can each serve as a general-purpose feature extractor for protein expression. For each of the four cell lines, we take the corresponding pretrained `seq2ribo` polisher, attach an expression head, and finetune on the same mRFP training split. As a baseline, we use the total predicted ribosome count from Translatomer as a scalar predictor of expression; this yields weak correlations across all cell lines (*r* ≤ 0.261).

As shown in Table 3, all four `seq2ribo` polishers achieve strong performance after finetuning, with Pearson correlations ranging from 0.830 (RPE-1) to 0.903 (iPSC). Notably, finetuned `seq2ribo` trained on iPSC, HEK293, or LCL Ribo-seq data surpasses the Pearson correlation of 0.85 reported by CodonBERT (Li et al., 2024) on the same dataset and splits. Figure 4 shows the tight agreement between predicted and measured expression for the iPSC and LCL polishers. The consistency across cell lines indicates that each polisher captures translational features that generalize to protein expression prediction, even though the polishers were trained on distinct Ribo-seq datasets. The modest variation in performance suggests that the choice of Ribo-seq training data introduces some cell-type bias, but does not prevent effective transfer. Stratified analysis shows that several polishers already predict higher ribosome load for more highly expressed sequences before expression-specific finetuning (Section 1.19 of the Supplementary Text). Perturbation analysis shows that expression predictions are comparably robust to noise as TE predictions, but more sensitive to dropout, and remain nearly invariant to scaling of the input profiles, indicating that the expression head relies primarily on the shape and relative distribution of ribosome density rather than on absolute count magnitudes (Section 1.17 of the Supplementary Text). Scatter plots of finetuned `seq2ribo` expression predictions for HEK293 and RPE-1, polisher total ribosome load versus measured expression for all four cell lines, and Translatomer predictions versus measured expression appear in Supplementary Figures S58–S60. Correlations between raw sTASEP and polisher total ribosome load and measured expression are reported in Supplementary Table S16.

**Table 3:** Protein expression prediction results across cell lines on the test set. We report Pearson correlation *r*. Bold indicates the best performance within each cell line.

| Cell line | Translat. (seq.) | seq2ribo |
| --- | --- | --- |
| iPSC | 0.052 | **0.903** |
| HEK293 | 0.000 | **0.883** |
| LCL | 0.261 | **0.873** |
| RPE-1 | 0.027 | **0.830** |

**Figure 4.**
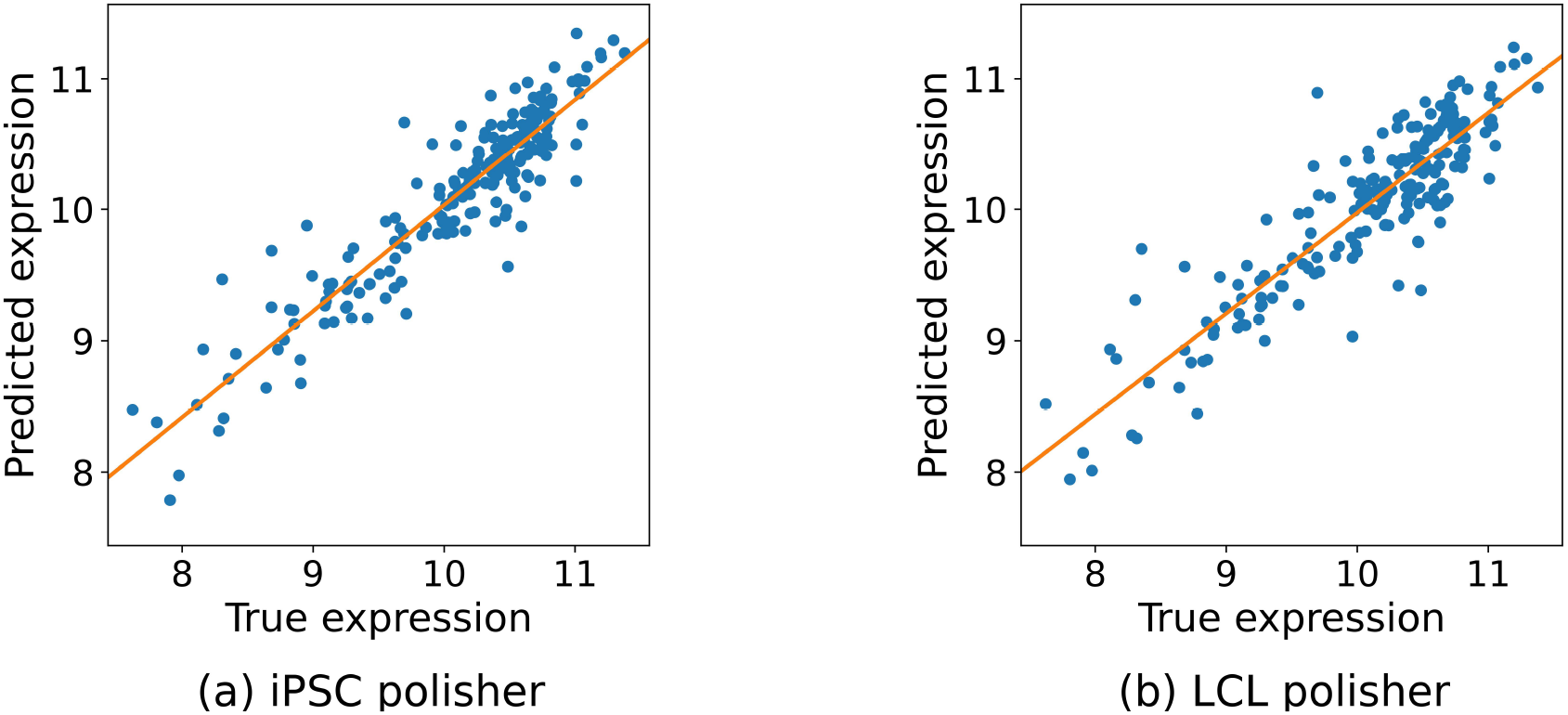
Correlation of finetuned `seq2ribo` predictions with measured protein expression on the mRFP test set. Each panel shows the result of finetuning the expression head on top of a different cell-line-specific polisher: (a) iPSC and (b) LCL.

### 3.5 Ablation studies on seq2ribo components

To deconstruct the `seq2ribo` pipeline and quantify the contribution of its core inputs, we conduct a set of ablation studies on the polisher module using the iPSC dataset. We systematically remove key inputs, namely the sTASEP simulation counts, the structural features and the sequence, and measure the effect on predictive accuracy. All models use the same architecture and optimization settings as the full polisher and are trained with the Poisson negative log-likelihood (NLL) loss on the iPSC training split, with checkpoints selected by validation loss. Detailed ablation procedures appear in Section 1.14 of the Supplementary Text. Table 4 summarizes the test set performance across all metrics. The MAE metrics in Table 4 are per-transcript-scaled; unscaled values appear in Supplementary Table S13.

**Table 4:** Summary of polisher feature ablation results on the iPSC test set. Lower is better for all MAE metrics, while higher is better for Pearson correlation. Bold indicates the best performance in each row.

|  | Seq. | sTASEP | Struc. | Seq+sTASEP | Seq+Struc. | sTASEP+Struc. | seq2ribo |
| --- | --- | --- | --- | --- | --- | --- | --- |
| *Ribosome profile prediction* |  |  |  |  |  |  |  |
| Tx-level $r$ | 0.898 | 0.216 | 0.205 | 0.913 | 0.903 | 0.258 | **0.920** |
| Shape $r$ | 0.129 | 0.058 | 0.026 | 0.120 | 0.102 | 0.051 | **0.162** |
| Elemwise $r$ | 0.554 | 0.102 | 0.084 | 0.563 | 0.546 | 0.128 | **0.598** |
| pertranscriptMAE | **1.28** | 1.29 | 1.32 | 1.29 | 1.30 | 1.30 | 1.39 |
| elemwiseMAE | **3.97** | 4.48 | 4.64 | 4.13 | 4.22 | 4.51 | 4.14 |
| codonMAE | 3 555 | 11 476 | 14 188 | **3 548** | 3 732 | 11 356 | 12 028 |
| pairMAE | 15 509 | 22 993 | 14 078 | 19 357 | **9 466** | 30 657 | 44 153 |
| angleMAE | **12 497** | 21 109 | 20 461 | 24 057 | 18 131 | 13 259 | 85 032 |
| bucketMAE | 68 032 | 554 629 | 411 693 | 46 308 | **31 270** | 214 407 | 287 231 |
| *Translation efficiency prediction* |  |  |  |  |  |  |  |
| Pearson $r$ | 0.654 | 0.578 | 0.388 | 0.660 | 0.624 | 0.584 | **0.688** |

All ablated models achieve comparable pertranscriptMAE (range 1.28–1.39), but they differ substantially in the primary correlation metrics. Sequence alone achieves Tx-level *r* of 0.898 and Shape *r* of 0.129, demonstrating strong baseline performance from codon identity. However, adding sTASEP simulation consistently improves both metrics: Seq+sTASEP raises Tx-level *r* to 0.913 and Shape *r* to 0.120, and the full `seq2ribo` model with all three inputs achieves the highest Tx-level *r* (0.920) and Shape *r* (0.162). The contribution of sTASEP is even more apparent when sequence is unavailable: Among sequence-free models, sTASEP alone attains a Tx-level *r* of 0.216, outperforming the structure-only model (0.205), and their combination raises this to 0.258, confirming that the simulation encodes translational information that is complementary to but independent of learned sequence representations. Together, these results support the hybrid two-stage design: the sTASEP simulation provides a mechanistic prior that the polisher exploits alongside sequence and structural features to produce the most accurate positional profiles.

We extend the ablation to translation efficiency prediction by attaching the TE head to each ablated polisher and finetuning on the mean TE task (procedure details in Section 1.15 of the Supplementary Text). As shown in Table 4, the full `seq2ribo` model achieves the highest Pearson correlation. Sequence alone and sTASEP alone are both strong individual predictors, each substantially outperforming structural features alone. All three components contribute to accuracy. This pattern mirrors the profile ablation and confirms that the mechanistic simulation, sequence, and structural features each provide complementary signals for TE prediction.

## 4 Discussion

We introduce `seq2ribo`, a hybrid framework that couples a structure-aware TASEP simulator with a Mamba-based polisher to predict ribosome dynamics from sequence alone. sTASEP encodes a mechanistic prior on ribosome traffic through fitted parameters for codon identity, local base pairing, structural angles, and positional context. The polisher refines these profiles, learning residual patterns the additive simulator cannot capture. This two-stage design preserves a physically interpretable core while achieving data-driven accuracy. sTASEP consistently improves on a classical TASEP baseline, reducing structural errors by up to 95.6%, and the full pipeline further improves positional agreement beyond either component alone. `seq2ribo` is the only method tested that produces ribosome profiles with genuine positional fidelity: transcript-level Pearson correlations reach 0.920 (versus 0.210 for the best simulator-only baseline and 0.055 for Translatomer), and within-transcript shape correlations reach 0.186, where every baseline returns near-zero or negative values.

A second key feature is that `seq2ribo` operates from sequence alone, making it directly applicable to de novo mRNA design where experimental covariates are unavailable. The predicted ribosome profiles serve as strong features for downstream tasks: `seq2ribo` outperforms RiboNN in the CDS-only setting across all four translation efficiency tasks, with gains of up to 30% in Pearson correlation, and reaches correlations up to 0.732 when UTR information is included. On a protein expression dataset, all four cell-line polishers achieve Pearson correlations above 0.83 after finetuning, indicating that `seq2ribo` captures translational dynamics relevant to protein output, not only ribosome positioning. Because the mRFP dataset lacks cell-type-specific labels, this evaluation primarily reflects sequence-intrinsic determinants of expression. However, the variation across polishers suggests that each encodes cell-line-specific translational features. As cell-type-resolved proteomics data become available, the framework can be extended by pairing each polisher with a dedicated expression head trained on matched labels, enabling disentanglement of sequence-intrinsic effects from cell-type-specific regulatory contributions.

Beyond these tasks, `seq2ribo` can generate synthetic Ribo-seq data for arbitrary sequences, including de novo designs or isoforms lacking experimental data. This enables in silico screening of candidate mRNA libraries to optimize ribosome traffic patterns—maximizing ribosome load or minimizing collision-prone motifs—before experimental synthesis, bridging the gap between sequence design and functional validation in synthetic biology.

Our study has limitations suggesting future directions. We currently derive structural features from predicted 2D RNA secondary structures, which do not fully capture 3D folding and tertiary interactions; future iterations could incorporate 3D predictions or co-transcriptional folding dynamics. We train separate models per cell line to maximize specificity; our cross-cell-line evaluation confirms that matched models substantially outperform transferred ones while revealing partial transferability that motivates a unified model capable of zero-shot transfer across tissues. We aim to extend the framework to other species to test cross-species generalization. The three-bucket positional parameterization captures coarse positional regimes but may be too coarse for very long transcripts; fixed-length buckets with regularization could provide finer resolution at the cost of additional parameters.

## Funding

This work was supported in part by the US National Science Foundation [III-2232121] and the US National Institutes of Health [R01HG012470].

## Data availability

The ribosome profiling data used in this study were obtained from the GWIPS-viz database (https://gwips.ucc.ie/). Specifically, we used aggregate A-site coverage tracks for the iPSC, HEK293, LCL, and RPE-1 cell lines. Gene annotations were obtained from GENCODE release v49 (https://www.gencodegenes.org/). The translation efficiency labels were obtained from Zheng et al. (2025). The mRFP protein expression dataset was obtained from Nieuwkoop et al. (2023). All processed datasets, including the matched sequences, structural features, and ribosome counts used for training and testing, are available at https://github.com/Kingsford-Group/seq2ribo.

## Code availability

The source code for `seq2ribo`, including the sTASEP simulator, the Mamba-based polisher, the task-specific finetuning pipelines for translation efficiency and protein expression, and scripts for data preprocessing, training, and evaluation, is available on GitHub at https://github.com/Kingsford-Group/seq2ribo. The repository also contains the pretrained model checkpoints for all four cell lines and the notebooks used to generate the figures in this manuscript. At inference time, profiling a single mRNA sequence requires seconds to a few minutes for the sTASEP simulation, depending on transcript length and the number of trajectories, followed by seconds for the polisher and any downstream prediction head. Full details on hardware, training times, and computational complexity are provided in Section 1.24 of the Supplementary Text.

## Author contributions statement

G.K. and C.K. conceived the study and designed the algorithms. G.K. implemented the `seq2ribo` framework, performed the data processing, trained the models, and conducted the experiments. G.K. and C.K. analyzed the results. G.K. wrote the manuscript with input from C.K.

## Acknowledgments

We thank Dr. David Koes for valuable discussions, Dr. Guillaume Marçais for help with system setup, Dr. Marçais and Shiyi Du for their careful review of the manuscript, and Jialin He for her feedback on training the sequence-only Translatomer model.

## Conflicts of interest

C.K. is a co-founder of Ocean Genomics, Inc.

## Notes

### Summary of Updates

Figures 3 and 4 revised; Tables 1, 2 and 3 revised; Supplemental files updated.

