## Supplementary Material for "seq2ribo: Structure-aware integration of machine learning and simulation to predict ribosome location profiles from RNA sequences"

Gün Kaynar and Carl Kingsford
Ray and Stephanie Lane Computational Biology Department
School of Computer Science
Carnegie Mellon University
{gkaynar,carlk}@cs.cmu.edu

### 1. Supplementary Text

#### 1.1 Data acquisition and preprocessing

We obtain global aggregate A-site ribosome counts from the GWIPS-viz Ribo-seq resource [16]. GWIPS-viz provides transcript-level A-site coverage tracks that summarize multiple Ribo-seq experiments and report the estimated A-site occupancy at each nucleotide position along annotated transcripts. The source data contain A-site ribosome counts for human transcripts across multiple conditions. We extract and process data for four distinct cell lines: iPSC [4], LCL [3], HEK293 [23], and RPE-1 [10].

We map these coverage tracks to coding sequences using GENCODE v49 [17] annotations on the GRCh38 reference genome. For each transcript in GENCODE, we identify the set of coding exons, order them along the genome, and construct the genomic interval that spans the coding sequence (CDS) from the annotated start codon to the annotated stop codon. We use the GRCh38/hg38 reference genome to extract the corresponding nucleotide sequence between these coding start and end coordinates. This yields a coding DNA sequence for each GENCODE transcript together with its genomic coordinates.

Next, we link the GWIPS-viz transcript-level A-site coverage tracks to the GENCODE v49 coding sequences. We use the transcript identifiers provided by GWIPS-viz and GENCODE to build a one-to-one mapping from GWIPS transcripts to GENCODE transcripts. We then intersect the mapped set of GENCODE transcripts with the GWIPS coverage tracks and keep only those transcripts for which we have both a valid coding sequence and a corresponding A-site coverage profile.

In the ribosome profile prediction task, for each matched transcript, we restrict the analysis to the annotated CDS. We discard intronic regions and untranslated regions and do not use any positions from the 5′ or 3′ untranslated regions. Likewise, we do not include any counts from introns or intergenic regions. For each coding transcript, we align the nucleotide-level A-site coverage from GWIPS-viz to the coding sequence that we extract from hg38 using the GENCODE coordinates. We require that the coverage track spans the entire coding region without gaps that would disrupt the reading frame. Transcripts for which we cannot establish a consistent mapping between the coverage track and the coding sequence are removed.

We then convert the nucleotide resolution A-site coverage into codon-resolution counts. For each coding sequence, we identify the reading frame from the annotated start codon and partition the coding sequence into contiguous codons of length three. For each codon, we take the sum of A-site counts at three nucleotides of that codon as the codon-level A-site count. We filter out coding sequences whose length is not a multiple of three, sequences that contain ambiguous nucleotides, sequences that contain an internal stop codon within the annotated coding region, and sequences that do not start with a start codon. We also discard any sequence for which the mapping between the A-site coverage and the coding sequence is inconsistent. We discard coding sequences whose total A-site ribosome count is zero, as these provide no information about occupancy.

After applying these filtering steps separately for each cell line, we obtain datasets used for training and evaluation. For each cell line, we split the transcripts into training (80%), validation (10%), and test (10%) sets, stratified by total A-site count. The transcript counts are:

- **iPSC:** 71,669 total transcripts (57,335 training, 7,167 validation, 7,167 test).
- **HEK293:** 60,772 total transcripts (48,617 training, 6,077 validation, 6,078 test).
- **RPE-1:** 66,130 total transcripts (52,904 training, 6,613 validation, 6,613 test).
- **LCL:** 73,210 total transcripts (58,568 training, 7,321 validation, 7,321 test).

We then remove from each test split any transcript whose coding sequence is identical to a coding sequence in the corresponding training split, which removes isoforms and other duplicate coding sequences across splits. After this filtering step, the test set sizes become 5,037 for iPSC, 4,218 for HEK293, 4,558 for RPE-1, and 5,133 for LCL.

For the translation efficiency prediction tasks, we cross-referenced our ribosome profiling transcripts with experimental TE values from Zheng et al. [25]. We matched transcripts based on common identifiers to create task-specific datasets for mean TE and cell-specific TE. The final dataset sizes for these tasks are:

- **Mean TE:** 8,096 training, 1,008 validation, and 1,020 test transcripts.
- **HEK293 TE:** 7,819 training, 984 validation, and 967 test transcripts.
- **RPE-1 TE:** 8,002 training, 993 validation, and 997 test transcripts.
- **LCL TE:** 8,045 training, 1,032 validation, and 987 test transcripts.

After applying the same duplicate-removal filtering to the TE test splits, the test set sizes become 733 for mean TE, 709 for HEK293 TE, 672 for RPE-1 TE, and 706 for LCL TE.

#### 1.2 Structural feature assembly

We use $D$ to denote the set of coding sequences we analyze, as defined in Methods Section 2.2 of the main text. We construct codon-level structural features $P_i$, $A_i$, and $B_i$ from the coding sequences in $D$ in three stages. We first compute a minimum free energy secondary structure and a two-dimensional layout for each transcript. We then derive codon resolution pair and angle features from this layout. Finally, we assign position buckets along each coding sequence.

For each CDS, we use the ViennaRNA package [14] to predict a minimum free energy secondary structure and obtain a dot-bracket representation. We then run the Turtle [24] graph-based layout algorithm in ViennaRNA to assign two-dimensional coordinates to every nucleotide in the sequence. This results in a secondary structure and a planar embedding with one coordinate pair for each nucleotide position.

We derive the codon-level pair feature from the base pairing pattern in the minimum free energy structure. For each nucleotide, we record whether it participates in a base pair. For each transcript $i$ and codon position $j$, we compute the number of nucleotides within that codon that are paired in the structure. This count ranges from zero to three. We set

$$p_i^j \in \{0, 1, 2, 3\}$$

to this count and define the pair count sequence

$$P_i = \left(p_i^1, \ldots, p_i^{N_i}\right).$$

This sequence is the pair feature that we use in the sTASEP simulator and in the polisher.

We derive the codon-level angle feature from the two-dimensional layout. Let $(x_k, y_k)$ denote the coordinates of nucleotide position $k$ along the coding sequence. For each interior nucleotide position $k$, we compute the interior vertex angle $\theta_k$ in radians formed by the vectors from $(x_k, y_k)$ to its two neighbors. For the first and last nucleotide, we set $\theta_k$ to zero.

We then aggregate these nucleotide-level angles to codon resolution. For transcript $i$ and codon position $j$, we define the raw codon angle

$$u_i^j = \theta_{k_1} + \theta_{k_2} + \theta_{k_3},$$

where $k_1$, $k_2$ and $k_3$ are the nucleotide indices that belong to codon $C_i^j$. This yields a continuous codon-level geometry signal $u_i^j$.

To obtain a feature that describes changes in local backbone geometry, we convert these raw angles into codon-wise angle differences. For $j \geq 2$ we define

$$\delta_i^j = u_i^j - u_i^{j-1},$$

and we set $\delta_i^1 = 0$.

We discretize the continuous values $\delta_i^j$ into four categories that represent no change, small change, medium change, and large change, and describe this discretization in Supplementary Section 1.3. We compute empirical thresholds from the distribution of $\delta_i^j$ values over all transcripts in $D_{\text{train}}$. We then assign a discrete angle category

$$a_i^j \in \{0, 1, 2, 3\}$$

to each codon position by thresholding $\delta_i^j$ according to these four bins. This defines the angle sequence

$$A_i = \left(a_i^1, \ldots, a_i^{N_i}\right),$$

which we use as the angle feature in sTASEP and in the polisher.

We define the bucket feature $B_i$ as a coarse positional encoding along the coding sequence. For a transcript $i$ with length $N_i$, we divide the codon indices into three segments of equal size up to rounding. Codons in the first third of the sequence receive bucket value 0, codons in the middle third receive bucket value 1, and codons in the final third receive bucket value 2. This yields

$$b_i^j \in \{0, 1, 2\} \quad \text{for } j = 1, \ldots, N_i,$$

and the bucket sequence

$$B_i = \left(b_i^1, \ldots, b_i^{N_i}\right).$$

We precompute and cache all pair, angle, and bucket features for every transcript to ensure reproducibility and to avoid repeated folding and layout computations.

**Rationale for the three-bucket design.** The three-bucket design reflects the observation that ribosome kinetics differ systematically across broad positional regimes of the coding sequence. The $5'$ region exhibits a "ramp" of slower elongation associated with rare codons and mRNA structure that is thought to regulate ribosome spacing and prevent downstream collisions [5, 22]. The central body of the CDS shows comparatively steady-state elongation. The $3'$ region shows distinct translational properties: in eukaryotes, the last $\sim$50 codons display the highest translational efficiency [22], while the stop codon region involves termination and ribosome recycling dynamics [12]. Ribosome profiling has confirmed that ribosome density varies systematically along the CDS, with a general decreasing trend consistent with initiation as the rate-limiting step [1, 13]. Three buckets capture these coarse positional regimes with only three additional parameters in the sTASEP model.

Our fitted parameters confirm this biological rationale. In iPSC and RPE-1, the first third of the CDS carries the lowest additional wait time ($\gamma_0 = 0.431$ and $0.440$, respectively), while the middle and final thirds are slower ($\gamma_1 = 0.536$, $\gamma_2 = 0.532$ for iPSC; $\gamma_1 = 0.528$, $\gamma_2 = 0.532$ for RPE-1), consistent with known ribosome density gradients along the CDS [13, 22]. These patterns emerge from data-driven fitting without any prior biological constraints on the parameter values, validating the three-bucket design as biologically meaningful (see Supplementary Table S14 and Supplementary Figures S20–S22 for the full parameter comparison).

An alternative design using fixed-length, variable-number buckets could capture finer positional patterns, particularly in long transcripts where the proportional thirds may be too coarse. We chose the proportional three-bucket design as a trade-off, because sTASEP fits bucket wait times through iterative Monte Carlo optimization, and increasing the number of bucket parameters risks overfitting to transcript-length-specific patterns in the training data. The three-bucket parameterization captures the dominant positional effects with minimal additional complexity. A regularized version of fixed-length buckets could potentially improve performance for very long transcripts and is a worthwhile direction for future investigation.

#### 1.3 Angle bin discretization

We discretize the continuous codon-level angle differences into four categories by estimating bin boundaries from the training set and then applying these boundaries unchanged to the validation and test sets. This procedure ensures that the structural features for the evaluation data do not influence the design of the bins.

As described above, we first compute a continuous codon-level geometry signal for each coding sequence. For transcript $i$ and codon position $j$, we obtain a raw codon angle $u_i^j$ from the two-dimensional layout and define the codon-wise angle difference

$$\delta_i^j = u_i^j - u_i^{j-1} \quad \text{for } j \geq 2,$$

with $\delta_i^1 = 0$. We store these values for each transcript in a geometry map.

To derive bin boundaries, we collect all codon-level angle differences from the training set. We iterate over the geometry map for $D_{\text{train}}$ and extract the vectors of angle differences for each transcript. We concatenate these vectors into a single array. This yields a collection of continuous values

$$\{\delta_i^j : i \in D_{\text{train}}, 1 \leq j \leq N_i\}$$

that summarizes the distribution of codon-wise angle changes in the training data.

We then compute bin edges for $K_{\text{angle}} = 4$ angle categories. To robustly define the range of angle changes and exclude extreme outliers, we select two cumulative probabilities, $p_{\text{min}} = 0$ and $p_{\text{max}} = 0.995$. We calculate the corresponding values from the training distribution $\delta$:

$$\ell = \text{Quantile}(\delta, p_{\text{min}}), \quad \text{(the minimum value)}$$

$$h = \text{Quantile}(\delta, p_{\text{max}}). \quad \text{(the 99.5th percentile)}$$

We then divide the range $[\ell, h]$ into discrete bins where $\delta$ ranges over all training set values. In practice, $\ell$ is close to the minimum and $h$ captures the bulk of the distribution up to the 99.5th percentile. If $h \leq \ell$, we enforce $h > \ell$ by adding a small constant.

We then construct $K_{\text{angle}} + 1 = 5$ bin edges by linearly spacing them between $\ell$ and $h$:

$$e_0, e_1, e_2, e_3, e_4 \quad \text{with} \quad e_0 = \ell, \quad e_4 = h, \quad e_{s+1} - e_s = \frac{h - \ell}{4} \text{ for } s = 0, \ldots, 3.$$

Finally, we set $e_4$ to the maximum of $e_4$ and the global maximum of the training set values to ensure that the last bin covers all observed angle differences in the training data. This yields a fixed set of monotonically increasing bin edges

$$e_0 < e_1 < e_2 < e_3 < e_4$$

that we use for all subsequent discretization steps.

We assign a discrete angle category to each codon by thresholding its codon-level angle difference against these bin edges. For transcript $i$ and codon $j$ we define

$$a_i^j = s \quad \text{if} \quad e_s \leq \delta_i^j < e_{s+1} \quad \text{for } s = 0, 1, 2,$$

and we assign $a_i^j = 3$ when $\delta_i^j \geq e_3$. This mapping yields

$$a_i^j \in \{0, 1, 2, 3\}$$

for each codon position and defines the angle sequence

$$A_i = \left(a_i^1, \ldots, a_i^{N_i}\right)$$

that we use in the sTASEP simulator and in the polisher.

We compute the bin edges $e_0, \ldots, e_4$ once using only the training set and then keep them fixed. We apply the same boundaries to discretize the codon-level angle differences for the validation and test transcripts. Because the validation and test angle features are assigned using thresholds derived solely from the training distribution, this procedure avoids any information leakage from the evaluation data into the feature construction.

##### 1.4 Evaluation metrics

We evaluate models with nine metrics that summarize agreement between predicted and observed A-site counts at different levels of aggregation. We treat two Pearson correlation metrics as primary indicators of prediction quality. We also report a third Pearson correlation metric and six mean absolute error (MAE) summaries. For any set of transcripts $\mathcal{S}$ and any model that outputs a codon-level A-site count sequence, we denote the observed count at codon $j$ of transcript $i$ by $y_i^j$. We denote the corresponding predicted count generically by $\hat{y}_i^j$ in the formulas below. Note that when evaluating the sTASEP simulator, $\hat{y}_i^j$ refers to the simulated output, and when evaluating the polisher, $\hat{y}_i^j$ refers to the refined output.

**Tx-level $r$:** Transcript-level Pearson correlation measures how well the model captures differences in overall expression across transcripts. For each transcript $i \in \mathcal{S}$, we compute the total observed and predicted counts

$$Y_i = \sum_{j=1}^{N_i} y_i^j, \qquad \hat{Y}_i = \sum_{j=1}^{N_i} \hat{y}_i^j.$$

We then compute the Pearson correlation of these transcript-level sums:

$$\text{Tx-level } r = \frac{\sum_{i \in \mathcal{S}} (Y_i - \bar{Y})(\hat{Y}_i - \bar{\hat{Y}})}{\sqrt{\sum_{i \in \mathcal{S}} (Y_i - \bar{Y})^2} \sqrt{\sum_{i \in \mathcal{S}} (\hat{Y}_i - \bar{\hat{Y}})^2}},$$

where $\bar{Y}$ and $\bar{\hat{Y}}$ are the means of $\{Y_i\}$ and $\{\hat{Y}_i\}$ over $\mathcal{S}$.

**Shape $r$:** Shape Pearson correlation measures how well the model captures the positional profile of ribosome occupancy along individual coding sequences. For each transcript $i \in \mathcal{S}$, we compute the Pearson correlation between the observed and predicted count vectors. Let

$$\bar{y}_i = \frac{1}{N_i} \sum_{j=1}^{N_i} y_i^j, \qquad \bar{\hat{y}}_i = \frac{1}{N_i} \sum_{j=1}^{N_i} \hat{y}_i^j,$$

and define the per-transcript correlation

$$r_i = \frac{\sum_{j=1}^{N_i} (y_i^j - \bar{y}_i)(\hat{y}_i^j - \bar{\hat{y}}_i)}{\sqrt{\sum_{j=1}^{N_i} (y_i^j - \bar{y}_i)^2} \sqrt{\sum_{j=1}^{N_i} (\hat{y}_i^j - \bar{\hat{y}}_i)^2}}.$$

We then average across transcripts:

$$\text{Shape } r = \frac{1}{|\mathcal{S}|} \sum_{i \in \mathcal{S}} r_i.$$

**Elemwise $r$:** Elementwise Pearson correlation measures linear agreement between predicted and observed counts across all codon positions in all transcripts. Unlike Shape $r$, which gives equal weight to each transcript and isolates within-transcript positional agreement, Elemwise $r$ pools all codon positions across all transcripts into a single correlation and is therefore influenced by differences in transcript length and ribosome load, which can cause highly loaded transcripts to dominate the metric. We retain Elemwise $r$ because the polisher's Poisson NLL loss operates at the codon level and thus most directly optimizes elementwise agreement, making this metric the natural reflection of what the model is trained to do. However, because it does not control for transcript-level differences, we treat Tx-level $r$ and Shape $r$ as the primary evaluation metrics. Let

$$M = \sum_{i \in \mathcal{S}} N_i,$$

and let $\{y_k\}_{k=1}^{M}$ and $\{\hat{y}_k\}_{k=1}^{M}$ denote the concatenated observed and predicted counts obtained by flattening $\{y_i^j\}$ and $\{\hat{y}_i^j\}$ over all $(i, j)$. Define the means

$$\bar{y} = \frac{1}{M} \sum_{k=1}^{M} y_k, \qquad \bar{\hat{y}} = \frac{1}{M} \sum_{k=1}^{M} \hat{y}_k.$$

We compute

$$\text{Elemwise } r = \frac{\sum_{k=1}^{M} (y_k - \bar{y})(\hat{y}_k - \bar{\hat{y}})}{\sqrt{\sum_{k=1}^{M} (y_k - \bar{y})^2} \sqrt{\sum_{k=1}^{M} (\hat{y}_k - \bar{\hat{y}})^2}}.$$

**pertranscriptMAE:** Per transcript mean absolute error first computes a mean absolute error for each transcript and then averages across transcripts:

$$\text{pertranscriptMAE} = \frac{1}{|\mathcal{S}|} \sum_{i \in \mathcal{S}} \left( \frac{1}{N_i} \sum_{j=1}^{N_i} |y_i^j - \hat{y}_i^j| \right).$$

This metric gives equal weight to each transcript rather than to each codon.

**elemwiseMAE:** Elementwise mean absolute error sums the absolute error across all codon positions and normalizes by the total number of codons:

$$\text{elemwiseMAE} = \frac{\sum_{i \in \mathcal{S}} \sum_{j=1}^{N_i} |y_i^j - \hat{y}_i^j|}{\sum_{i \in \mathcal{S}} N_i}.$$

This metric gives equal weight to each codon in the dataset.

**codonMAE:** Codon mean absolute error aggregates errors by codon identity, in line with the codon-specific wait time parameters of the sTASEP simulator. Let $\mathcal{C}$ denote the set of 61 non-stop codons, and let $\text{codon}(C_i^j) \in \mathcal{C}$ denote the codon at position $j$ of transcript $i$. For each codon type $c \in \mathcal{C}$, we compute the total predicted and observed counts

$$\hat{Y}_c = \sum_{i \in \mathcal{S}} \sum_{j:\text{codon}(C_i^j)=c} \hat{y}_i^j, \qquad Y_c = \sum_{i \in \mathcal{S}} \sum_{j:\text{codon}(C_i^j)=c} y_i^j,$$

and define

$$\text{codonMAE} = \frac{1}{|\mathcal{C}|} \sum_{c \in \mathcal{C}} \left| Y_c - \hat{Y}_c \right|.$$

**pairMAE:** Pair mean absolute error aggregates errors by base pairing category, which corresponds to the pair wait time parameters in sTASEP. For each transcript we have $p_i^j \in \{0, 1, 2, 3\}$, which records how many nucleotides in codon $j$ participate in base pairing. For each pair category $r \in \{0, 1, 2, 3\}$ we compute

$$\hat{Y}_r^{\text{pair}} = \sum_{i \in \mathcal{S}} \sum_{j:p_i^j=r} \hat{y}_i^j, \qquad Y_r^{\text{pair}} = \sum_{i \in \mathcal{S}} \sum_{j:p_i^j=r} y_i^j,$$

and define

$$\text{pairMAE} = \frac{1}{4} \sum_{r=0}^{3} \left| Y_r^{\text{pair}} - \hat{Y}_r^{\text{pair}} \right|.$$

**angleMAE:** Angle mean absolute error aggregates errors by discretized local backbone angle change, which corresponds to the angle wait time parameters in sTASEP. For each transcript, we have $a_i^j \in \{0, 1, 2, 3\}$, which encodes the local angle change at codon $j$. For each angle category $s \in \{0, 1, 2, 3\}$ we compute

$$\hat{Y}_s^{\text{angle}} = \sum_{i \in \mathcal{S}} \sum_{j:a_i^j=s} \hat{y}_i^j, \qquad Y_s^{\text{angle}} = \sum_{i \in \mathcal{S}} \sum_{j:a_i^j=s} y_i^j,$$

and define

$$\text{angleMAE} = \frac{1}{4} \sum_{s=0}^{3} \left| Y_s^{\text{angle}} - \hat{Y}_s^{\text{angle}} \right|.$$

**bucketMAE:** Bucket mean absolute error aggregates errors by coarse position along the coding sequence, which corresponds to the bucket wait time parameters in sTASEP. For each transcript, we have $b_i^j \in \{0, 1, 2\}$, which marks the first third, middle third, and final third of the coding sequence. For each bucket $t \in \{0, 1, 2\}$ we compute

$$\hat{Y}_t^{\text{bucket}} = \sum_{i \in \mathcal{S}} \sum_{j : b_i^j = t} \hat{y}_i^j, \qquad Y_t^{\text{bucket}} = \sum_{i \in \mathcal{S}} \sum_{j : b_i^j = t} y_i^j,$$

and define

$$\text{bucketMAE} = \frac{1}{3} \sum_{t=0}^{2} \left| Y_t^{\text{bucket}} - \hat{Y}_t^{\text{bucket}} \right|.$$

Higher values are better for the three correlation metrics, and lower values are better for the six MAE metrics. Together, these metrics characterize prediction accuracy at the level of individual codons, full transcripts, codon identities, structural and positional groups, and the within-transcript positional profile.

**Scaled MAE metrics.** We report scaled and unscaled (raw) variants of all six MAE metrics defined above. For a given transcript $i$, let

$$Y_i = \sum_{j=1}^{N_i} y_i^j, \qquad \hat{Y}_i = \sum_{j=1}^{N_i} \hat{y}_i^j$$

denote the total observed and predicted ribosome loads. The scaled predicted counts are

$$\hat{y}_i^{j,\text{sc}} = \begin{cases} \hat{y}_i^j \cdot \dfrac{Y_i}{\hat{Y}_i} & \text{if } \hat{Y}_i > 0, \\ 0 & \text{otherwise,} \end{cases}$$

so that the total predicted ribosome load on each transcript matches the total observed ribosome load exactly. The scaled MAE metrics are computed by substituting $\hat{y}_i^{j,\text{sc}}$ for $\hat{y}_i^j$ in the formulas above. By equalizing the ribosome load between ground truth and predicted profiles, these scaled metrics isolate errors in the positional distribution of ribosome occupancy from errors in overall transcript-level magnitude.

#### 1.5 sTASEP simulation schedule and parameter fitting

We implement the sTASEP dynamics described in Section 2.3 of the main text with a Monte Carlo simulation and an outer loop that updates wait time parameters.

For each transcript $i$ in $D_{\text{train}}$, we run 100 independent sTASEP trajectories with a fixed set of wait time parameters. In each trajectory, we start from an empty lattice. We simulate until we observe 10 termination events. This defines a burn-in period that lets the system reach a steady operating regime. After burn-in, at each simulation step, we stop the trajectory with probability 0.01. At the stopping time, we record the positions of all ribosomes and convert these positions into predicted A-site counts $\hat{y}_i^j$ at each codon position. We aggregate these counts across all trajectories and transcripts to obtain the simulator output $\hat{Y}_i$ for the current parameters.

We collect all wait time parameters into vectors,

$$\vec{\tau} \text{ for codons,} \qquad \vec{\alpha} \text{ for pair categories,} \qquad \vec{\beta} \text{ for angle categories,} \qquad \vec{\gamma} \text{ for bucket categories.}$$

We initialize every element in these vectors to 0.5 and then refine them by iterating between simulation and parameter updates.

Because the sTASEP uses a single initiation rate for all transcripts, the simulator produces ribosome profiles whose overall magnitude is governed by this shared rate rather than by transcript-specific expression levels. If we were to compare simulated and observed counts directly, the gradient proxies would be dominated by transcripts with high ribosome loads and the parameters would be driven toward reproducing the average count level rather than learning the correct positional distribution. To avoid this, we scale the simulated counts on each transcript to match the observed ribosome load before computing errors. Concretely, for each transcript $i$, we replace the raw simulated counts $\hat{y}_i^j$ with their scaled versions $\hat{y}_i^{j,\text{sc}} = \hat{y}_i^j \cdot Y_i / \hat{Y}_i$, where $Y_i = \sum_j y_i^j$ and $\hat{Y}_i = \sum_j \hat{y}_i^j$ are the total observed and simulated ribosome loads. All aggregated counts and gradient proxies described below use these scaled counts. This ensures that the parameter updates are driven entirely by discrepancies in the positional shape of the ribosome profile.

After each round of simulations, we compare scaled simulated counts with observed counts and compute gradient proxies at the level of codon, pair, angle, and bucket groups. For codon-specific parameters, we aggregate counts by codon identity. For each non-stop codon $c$, we compute the total scaled simulated and observed counts

$$\hat{Y}_c^{\text{sc}} = \sum_{i \in D_{\text{train}}} \sum_{j: C_i^j = c} \hat{y}_i^{j,\text{sc}}, \qquad Y_c = \sum_{i \in D_{\text{train}}} \sum_{j: C_i^j = c} y_i^j,$$

and the number of occurrences

$$n_c = \sum_{i \in D_{\text{train}}} \sum_{j: C_i^j = c} 1.$$

We then define a gradient proxy

$$\Delta_c = \frac{\hat{Y}_c^{\mathrm{sc}} - Y_c}{n_c}$$

and update the codon wait time

$$\tau_c \leftarrow \max(\tau_c - \eta_\tau \cdot \Delta_c, \varepsilon),$$

where $\eta_\tau$ is a learning rate and $\varepsilon = 10^{-12}$ is the minimum allowed wait time. We set $\eta_\tau$ to 0.01. When the simulator overestimates the occupancy of codon $c$, so that $\hat{Y}_c^{\mathrm{sc}} > Y_c$, this update decreases $\tau_c$, which increases the elongation rate through $r_i^j$ and reduces ribosome dwell time at that codon in the next simulation round. For pair, angle, and bucket parameters, we use centered updates and describe them in the next section.

We run this simulation and update procedure for 100 outer iterations. The final simulator $f$ uses the wait time parameters obtained after the last iteration and produces the simulated A-site count sequences $\hat{Y}_i$. All ribosome profile prediction results that we report in this work use this fixed parameter set and are evaluated on $D_{\mathrm{test}}$.

##### 1.5.1 Centered updates for structural wait time parameters

We use centered updates for the pair, angle, and bucket wait time parameters to separate relative effects from a global shift within each group of categories. We illustrate the procedure for the pair parameters, and we use the same scheme for the angle and bucket parameters.

For the pair categories we maintain a category-specific vector $\vec{\alpha} = (\alpha_r)$ and a scalar bias term $\alpha_{\mathrm{bias}}$. During one outer simulation iteration, the wait time contribution for pair category $r$ is

$$\alpha_r + \alpha_{\mathrm{bias}}.$$

After each iteration, we compute gradient proxies $\Delta_r$ for each pair category. Let $\mathcal{R}$ denote the set of pair categories $\{0, 1, 2, 3\}$. For each category $r \in \mathcal{R}$, we aggregate simulated and observed counts over all positions with $p_i^j = r$ in the training set and normalize by the number of such positions. This gives a category-specific proxy $\Delta_r$ that measures the average over prediction or under prediction for that category. Concretely, we define

$$\hat{Y}_r^{\mathrm{pair}} = \sum_{i \in D_{\mathrm{train}}} \sum_{j : p_i^j = r} \hat{y}_i^j, \qquad Y_r^{\mathrm{pair}} = \sum_{i \in D_{\mathrm{train}}} \sum_{j : p_i^j = r} y_i^j,$$

$$n_r = \sum_{i \in D_{\mathrm{train}}} \sum_{j : p_i^j = r} 1, \qquad \Delta_r = \frac{\hat{Y}_r^{\mathrm{pair}} - Y_r^{\mathrm{pair}}}{n_r}.$$

We then compute the mean gradient proxy

$$\bar{\Delta}_\alpha = \frac{1}{|\mathcal{R}|} \sum_{r \in \mathcal{R}} \Delta_r.$$

We update the relative pair parameters and the global bias as

$$\alpha_r \leftarrow \alpha_r - \eta_\alpha \cdot \left(\Delta_r - \bar{\Delta}_\alpha\right),$$

$$\alpha_{\text{bias}} \leftarrow \alpha_{\text{bias}} - \eta_\alpha \cdot \left(\lambda \cdot \bar{\Delta}_\alpha\right),$$

where $\eta_\alpha$ is a learning rate and $\lambda$ controls the speed of the bias update. We set $\eta_\alpha = 0.01$ and $\lambda = 0.5$ in all experiments.

The term $\Delta_r - \bar{\Delta}_\alpha$ adjusts the relative differences among pair categories. The mean $\bar{\Delta}_\alpha$ adjusts the shared bias term and thus captures a global shift across all pair categories.

At the end of the update step, we fold the bias into the category-specific parameters by setting

$$\alpha_r \leftarrow \alpha_r + \alpha_{\text{bias}} \quad \text{for all } r \in \mathcal{R},$$

and we reset

$$\alpha_{\text{bias}} \leftarrow 0.$$

The next simulation iteration then uses the updated pair wait times $\alpha_r$ directly.

We apply the same centered update and bias folding scheme to the angle wait times $\vec{\beta}$ with learning rate $\eta_\beta = 0.01$ and to the bucket wait times $\vec{\gamma}$ with learning rate $\eta_\gamma = 0.01$. For each group, we maintain a separate bias term and compute the corresponding mean gradient proxy in the same way as for the pair parameters.

##### 1.6 Mamba-based polisher architecture and training

We provide the full architecture and training setup. For a transcript $i$ with length $N_i$, we construct five input channels.

- Codon embedding: We map codon sequences to an index set of size 65 that includes 61 sense codons, three stop codons, and one padding symbol. We apply a learnable embedding layer that maps this index to a 192 dimensional vector, which yields

$$E_i^{\text{codon}} \in \mathbb{R}^{N_i \times 192}.$$

- Structural (Geometric) embeddings. We embed the structural feature sequences in the same way. The pair feature and the angle feature each take four discrete values, and the bucket feature takes three discrete values. We embed each feature into 192 dimensions to obtain

$$E_i^{\text{pair}}, E_i^{\text{angle}}, E_i^{\text{bucket}} \in \mathbb{R}^{N_i \times 192}.$$

- Simulation embedding. We embed the simulated A-site counts from the sTASEP simulator $f$ with a learned linear projection from dimension one to 192,

$$E_i^{\text{sim}} \in \mathbb{R}^{N_i \times 192}.$$

We sum the five embeddings position-wise to obtain the initial hidden representation

$$H_i^{(0)} = E_i^{\text{codon}} + E_i^{\text{pair}} + E_i^{\text{angle}} + E_i^{\text{bucket}} + E_i^{\text{sim}},$$

which we feed into the Mamba backbone. The backbone consists of four stacked Mamba blocks with model dimension $d_{\text{model}} = 192$, state dimension $d_{\text{state}} = 16$, convolution width $d_{\text{conv}} = 4$, and expansion factor 2 as in Gu and Dao [9]. For block index $\ell = 1, \ldots, 4$ we compute

$$Z_i^{(\ell)} = \text{LN}\big(H_i^{(\ell-1)}\big), \quad U_i^{(\ell)} = \text{Mamba}\big(Z_i^{(\ell)}\big), \quad H_i^{(\ell)} = H_i^{(\ell-1)} + U_i^{(\ell)}.$$

The Mamba layers share parameters across positions and operate along the sequence length. After the fourth Mamba block, we apply a position-wise feed-forward head consisting of a linear transformation from dimension 192 to 192, the Gaussian Error Linear Unit (GELU) nonlinearity, dropout with rate 0.1, and a final linear transformation from 192 to a single scalar per codon position. To enforce non-negative predicted counts, we apply a Softplus nonlinearity (with default parameter $\beta = 1$) to the output. This produces the polished sequence

$$\tilde{Y}_i = g\big(C_i, P_i, A_i, B_i, \hat{Y}_i\big) = \big(\tilde{y}_i^1, \ldots, \tilde{y}_i^{N_i}\big).$$

We train the polisher on $D_{\text{train}}$ using multiple simulator trajectories per transcript as inputs. For each transcript, we run the trained sTASEP simulator $f$ 32 times and treat each simulation output as an input sequence to $g$. All 32 simulations share the same ground truth A-site counts $Y_i$. During training, we minimize a Poisson negative log-likelihood (NLL) loss. Let $\tilde{y}_i^j$ denote the polisher output at codon $j$ of transcript $i$ before the Softplus activation, and let $y_i^j$ denote the observed count. The per-position loss is

$$\ell_i^j = \exp(\tilde{y}_i^j) - y_i^j \cdot \tilde{y}_i^j + \log(y_i^j!),$$

which corresponds to the negative log-likelihood of a Poisson distribution with log-rate $\tilde{y}_i^j$. We average this loss over all unmasked codon positions in each batch:

$$\mathcal{L}_{\text{NLL}} = \frac{\sum_{i \in \mathcal{B}} \sum_{j=1}^{N_i} \ell_i^j}{\sum_{i \in \mathcal{B}} N_i}.$$

For the HEK293 cell line, we observed that the ribosome profiling data are substantially sparser than in the other cell lines, with a larger fraction of codon positions having zero observed counts. To prevent the model from learning to predict zero everywhere, we upweight the loss at positions with non-zero observed counts by a factor of 5, replacing $\ell_i^j$ with $5 \cdot \ell_i^j$ whenever $y_i^j > 0$. We do not apply this weighting for the other three cell lines.

We optimize the parameters of $g$ with the Adam optimizer with learning rate $10^{-4}$ and weight decay $2 \times 10^{-5}$. We apply gradient clipping with a maximum global norm of 1.0. After each training epoch, we compute the Poisson NLL on $D_{\text{val}}$ and retain the checkpoint with the lowest validation loss. We reduce the learning rate by a factor of 0.5 when the validation loss does not improve for three consecutive epochs. All polisher and `seq2ribo` results that we report use this selected model and are evaluated on $D_{\text{test}}$.

#### 1.7 Downstream heads for translation efficiency and protein expression

##### 1.7.1 Translation efficiency

**CDS-only TE head:** For a transcript $i$, we first obtain the polished codon-level profile

$$\tilde{Y}_i = (\tilde{y}_i^1, \ldots, \tilde{y}_i^{N_i})$$

from the fixed sTASEP simulator and Mamba-based polisher. The TE head operates in two stages: a per-position transformation followed by pooling. We pass each position independently through a feed-forward network with three linear layers of width 256, 128, and 1. Each hidden layer applies a linear transformation followed by a GELU nonlinearity. This produces a learned scalar at each codon position. We then apply masked mean pooling over positions to obtain a single value per transcript. A final linear layer maps this pooled value to a scalar, and a sigmoid nonlinearity maps the output to the interval $[0, 1]$.

We map the experimental TE labels to $[0, 1]$ using a min-max transform computed from the observed range of TE values in the training set. We train the head parameters and finetune the polisher $g$ end-to-end with a mean squared error (MSE) loss between predicted and target values in this scaled space. At test time, we invert the transform to recover predictions in the original TE scale.

For each labeled transcript, we run the trained sTASEP simulator $f$ 100 times, polish each simulated profile with the Mamba-based polisher $g$, and obtain 100 codon-level profiles. All 100 profiles share the same experimental translation efficiency label and serve as inputs to the TE head during training. We optimize with Adam using a learning rate of $10^{-4}$, weight decay $10^{-4}$, and gradient clipping with a maximum global norm of 1.0. We train for 10 epochs with batch size 64 and select the best checkpoint by validation MSE in the original TE scale. At test time, we run the trained pipeline on each of the 100 simulated profiles for a given test sequence, obtain 100 TE predictions, and report the average as the final prediction for that sequence.

**UTR-aware TE head:** To enable a fair comparison with RiboNN in the setting where both $5'$ and $3'$ UTR sequences are available, we also train a UTR-aware variant of the `seq2ribo` TE head. This variant replaces the CDS-only feed-forward head with a multi-region architecture that processes the $5'$ UTR, the CDS count profile, and the $3'$ UTR through three independent Mamba sequence blocks.

For the $5'$ and $3'$ UTR regions, we tokenize each nucleotide sequence into single-nucleotide tokens (A, U, G, C) and embed them into a shared 128-dimensional space using a learnable nucleotide embedding. For the CDS region, we apply $\log(1 + x)$ to each polished codon count, project the scalar to 128 dimensions with a learned linear layer. Each of the three regions is then processed by its own Mamba block consisting of two Mamba layers with model dimension 128, state dimension 16, convolution width 4, and expansion factor 2. We apply masked mean pooling to each block's output to obtain one 128-dimensional summary per region.

In addition, we compute a count summary by taking the masked mean of $\log(1 + \tilde{y}_i^j)$ across CDS positions and projecting the resulting scalar through a linear layer and GELU nonlinearity to a 128-dimensional vector. We concatenate the four summaries (count summary, $5'$ UTR, CDS, $3'$ UTR) to form a 512-dimensional representation and pass it through a feed-forward network with hidden sizes 128 and 64, GELU nonlinearities, and dropout with rate 0.1. A final linear layer maps to a single scalar, and a sigmoid nonlinearity produces the output in $[0, 1]$.

We use the same min-max transform, MSE loss in scaled space, optimizer settings, and training schedule as the CDS-only head. The polisher backbone is finetuned end-to-end together with the UTR-aware head.

##### 1.7.2 Protein expression

For the mRFP protein expression dataset, we apply the same per-position-then-pool design as the CDS-only TE head. We pass each codon position independently through a feed-forward network with four linear layers of width 128, 64, 64, and 1, with GELU nonlinearities between hidden layers. We apply masked mean pooling over the resulting per-position scalars, and a final linear layer outputs a single protein expression prediction. Unlike the TE head, we do not apply a sigmoid nonlinearity, as the expression values do not require a bounded output range.

We generate 1,000 simulated ribosome count profiles for each training, validation, and test sequence in the mRFP dataset. We finetune the full model (polisher backbone and expression head) on the training sequences using their experimental protein expression values with an MSE loss. We optimize with Adam using a learning rate of $10^{-4}$, weight decay $2 \times 10^{-4}$, and gradient clipping with a maximum global norm of 1.0. We train for 3 epochs with batch size 512 and use the validation sequences for early stopping. At test time, we run the trained pipeline on each of the 1,000 simulated profiles for a given test sequence, obtain 1,000 expression predictions, and report the average as the final prediction for that sequence.

##### 1.8 The classical TASEP model

We implement a classical codon-level TASEP model as a baseline. This model uses the same discrete-time stochastic simulation framework as our sTASEP model, including the definitions of initiation, elongation, termination, and steric exclusion. In particular, initiation attempts place a ribosome at the first codon with a four codon footprint, elongation enforces a three codon forward footprint, and termination removes a ribosome when it advances onto the stop codon of the coding sequence.

The key difference lies in the parameterization of the elongation rate. In the classical TASEP model, the ribosome wait time at codon position $j$ of transcript $i$ depends only on the codon identity and ignores all structural or positional context:

$$w_i^j = \tau_{\text{codon}(C_i^j)}.$$

The elongation rate at that position is then

$$r_i^j = \frac{1}{\max\left(w_i^j, \varepsilon\right)},$$

with the same numerical value of $\varepsilon$ as in sTASEP. The only trainable parameters are the 61 codon-specific wait times $\vec{\tau} = (\tau_c)_{c \in \mathcal{C}}$, one for each non-stop codon. This model is therefore a special case of sTASEP where the structural parameter vectors $\vec{\alpha}$, $\vec{\beta}$, and $\vec{\gamma}$ are fixed to zero.

We optimize the parameters $\vec{\tau}$ using the same simulation-based gradient descent procedure that we use for sTASEP, but applied only to the codon parameters. After each outer simulation iteration, we aggregate simulated and observed counts by codon identity over the training set. For each non-stop codon $c \in \mathcal{C}$ we compute

$$\hat{Y}_c = \sum_{i \in D_{\text{train}}} \sum_{j: C_i^j = c} \hat{y}_i^j, \qquad Y_c = \sum_{i \in D_{\text{train}}} \sum_{j: C_i^j = c} y_i^j,$$

and the number of occurrences

$$n_c = \sum_{i \in D_{\text{train}}} \sum_{j: C_i^j = c} 1.$$

We define the gradient proxy

$$\Delta_c = \frac{\hat{Y}_c - Y_c}{n_c}$$

and update the codon wait time

$$\tau_c \leftarrow \max\left(\tau_c - \eta_\tau \cdot \Delta_c, \varepsilon\right),$$

where $\eta_\tau$ is a learning rate. We set $\eta_\tau = 0.01$. When the simulator overestimates the occupancy of codon $c$, so that $\hat{Y}_c > Y_c$, this update decreases $\tau_c$, which increases the elongation rate and reduces ribosome dwell time at that codon in the next simulation round. This update fits the codon wait times.

##### 1.9 Direct parameter estimation via observed occupancy

As an alternative to iterative simulation-based optimization, we also formulate a direct non-iterative estimation method to approximate the sTASEP wait time parameters. This method provides an analytical first guess based on the assumption that the average observed ribosome occupancy within any given feature bin is directly proportional to the wait time associated with that bin.

We perform this estimation in a single pass over the training set $D_{\text{train}}$. For each group of parameters, we aggregate two statistics for each bin: the total observed A-site counts and the total number of codon occurrences in that bin.

For codon-specific parameters, we let $\mathcal{C}$ denote the set of 61 non-stop codons. For each codon $c \in \mathcal{C}$ we compute

$$Y_c = \sum_{i \in D_{\text{train}}} \sum_{j : C_i^j = c} y_i^j, \qquad n_c = \sum_{i \in D_{\text{train}}} \sum_{j : C_i^j = c} 1,$$

and define the average observed occupancy

$$\bar{Y}_c = \frac{Y_c}{n_c}.$$

For the structural parameters, we proceed in the same way. For the pair categories we let $\mathcal{R} = \{0, 1, 2, 3\}$ and define

$$Y_r^{\text{pair}} = \sum_{i \in D_{\text{train}}} \sum_{j : p_i^j = r} y_i^j, \qquad n_r^{\text{pair}} = \sum_{i \in D_{\text{train}}} \sum_{j : p_i^j = r} 1,$$

$$\bar{Y}_r^{\text{pair}} = \frac{Y_r^{\text{pair}}}{n_r^{\text{pair}}}.$$

For the angle categories we let $\mathcal{S} = \{0, 1, 2, 3\}$ and define

$$Y_s^{\text{angle}} = \sum_{i \in D_{\text{train}}} \sum_{j : a_i^j = s} y_i^j, \qquad n_s^{\text{angle}} = \sum_{i \in D_{\text{train}}} \sum_{j : a_i^j = s} 1,$$

$$\bar{Y}_s^{\text{angle}} = \frac{Y_s^{\text{angle}}}{n_s^{\text{angle}}}.$$

For the bucket categories we let $\mathcal{T} = \{0, 1, 2\}$ and define

$$Y_t^{\text{bucket}} = \sum_{i \in D_{\text{train}}} \sum_{j : b_i^j = t} y_i^j, \qquad n_t^{\text{bucket}} = \sum_{i \in D_{\text{train}}} \sum_{j : b_i^j = t} 1,$$

$$\bar{Y}_t^{\text{bucket}} = \frac{Y_t^{\text{bucket}}}{n_t^{\text{bucket}}}.$$

These average occupancies provide unscaled proxies for the wait times in each codon, pair, angle, and bucket bin.

We then normalize the average occupancy vectors to obtain relative wait time parameters with a unit sum within each group. For codons, we define

$$\tau'_c = \frac{\bar{Y}_c}{\sum\limits_{c' \in \mathcal{C}} \bar{Y}_{c'}} \quad \text{for all } c \in \mathcal{C},$$

and collect them into a vector $\vec{\tau}' = (\tau'_c)_{c \in \mathcal{C}}$.

For the pair parameters, we define

$$\alpha'_r = \frac{\bar{Y}_r^{\text{pair}}}{\sum\limits_{r' \in \mathcal{R}} \bar{Y}_{r'}^{\text{pair}}} \quad \text{for all } r \in \mathcal{R},$$

and collect them into a vector $\vec{\alpha}' = (\alpha'_r)_{r \in \mathcal{R}}$.

For the angle parameters, we define

$$\beta'_s = \frac{\bar{Y}_s^{\text{angle}}}{\sum\limits_{s' \in \mathcal{S}} \bar{Y}_{s'}^{\text{angle}}} \quad \text{for all } s \in \mathcal{S},$$

and collect them into a vector $\vec{\beta}' = (\beta'_s)_{s \in \mathcal{S}}$.

For the bucket parameters, we define

$$\gamma'_t = \frac{\bar{Y}_t^{\text{bucket}}}{\sum\limits_{t' \in \mathcal{T}} \bar{Y}_{t'}^{\text{bucket}}} \quad \text{for all } t \in \mathcal{T},$$

and collect them into a vector $\vec{\gamma}' = (\gamma'_t)_{t \in \mathcal{T}}$.

These normalized vectors $\vec{\tau}'$, $\vec{\alpha}'$, $\vec{\beta}'$, and $\vec{\gamma}'$ provide an analytical approximation of the sTASEP wait time parameters that we derive directly from the observed occupancy patterns in the training data rather than from iterative simulations.

We evaluate these analytically derived parameters as an alternative initialization strategy for the simulation-based optimization of sTASEP and compare them with a simple uniform initialization where we set all entries of $\vec{\tau}$, $\vec{\alpha}$, $\vec{\beta}$, and $\vec{\gamma}$ to 0.5. We observe that initialization with the analytical first estimate does not produce a clear improvement in the final converged performance. Since the simulation-based optimization converges to a robust solution even from the uniform initialization, we use the uniform initialization as the starting point for the final sTASEP model training.

##### 1.10 Translatomer

We use Translatomer [11] as an external baseline for ribosome profile prediction. In its original form, Translatomer ingests genomic DNA in a fixed-width window together with RNA sequencing coverage as a sixth input channel and predicts ribosome footprint density at base pair resolution. To match our setting, in which the model must predict ribosome occupancy from sequence alone without access to RNA-seq data, we train a sequence-only variant that removes the RNA-seq channel from both the architecture and the inputs. Specifically, we replace the six-channel input convolution with a five-channel convolution over one-hot encoded DNA (A, T, C, G, and an ambiguous-base channel) and keep the rest of the architecture unchanged. We train four separate sequence-only Translatomer models, one per cell line, to predict A-site ribosome counts from genomic sequence alone.

**Training.** For each cell line, we prepare training data by mapping each transcript to its genomic coordinates using GENCODE v49 annotations. For each transcript, we identify the CDS exon intervals and tile the CDS with genomic windows of length 65,536 base pairs. If the CDS fits within a single window, we center the window on the CDS midpoint. For longer coding sequences, we tile with a stride of 32,768 base pairs and add one final window anchored at the CDS end to ensure full coverage. For transcripts on the reverse strand, we reverse-complement the input sequence and flip the target profile.

Within each window, we construct the target by mapping the observed nucleotide-level A-site counts from the CDS back to their genomic positions using the exon intervals. Intronic and non-CDS positions within the window receive a target of zero. We then bin the 65,536 base pair target into 1,024 bins of 64 base pairs each by summing the counts within each bin. The model outputs a predicted count profile of 1,024 bins per window.

We split the training transcripts into 90% training and 10% validation. We train with the Adam optimizer at a learning rate of $10^{-5}$, batch size 32, and dropout rate 0.1 for 10 epochs. We select the best checkpoint by validation loss.

**Inference and evaluation.** For each transcript in the test set, we identify all genomic windows of length 65,536 base pairs that cover the CDS and provide the corresponding genomic sequence window as input to Translatomer. For each window, the model outputs a ribosome count profile in 1,024 bins at 64 base pair resolution. We aggregate these predictions by averaging the outputs from all windows that cover the CDS of a given transcript, which yields a single binned count profile per transcript. We then map the binned count profile to a codon-level count vector along the CDS by assigning each codon to the bin that covers its genomic coordinates and interpolating as needed for boundary cases. This produces a codon-level predicted count vector $c_i$ for transcript $i$.

**Translation efficiency and protein expression prediction.** For the downstream TE and protein expression tasks, we use the total predicted ribosome count per transcript from Translatomer as a simple scalar predictor. Specifically, for each transcript $i$, we compute the sum of the codon-level predicted count vector $\sum_j c_i^j$ and use this total ribosome load as the predicted value. We then compute Pearson correlation between this total load and the experimental TE or protein expression labels on the test set. This mirrors the evaluation we perform for the `seq2ribo` polisher output before finetuning (i.e., without a task-specific head), and provides a direct comparison of how well each model's predicted ribosome profile captures translational output from sequence alone.

##### 1.11 RiboNN

For translation efficiency prediction, we compare the `seq2ribo` translation efficiency heads with RiboNN [25]. We evaluate RiboNN in two input settings and two training strategies.

**UTR+CDS (original setting).** In the original RiboNN setting, the model ingests the full transcript RNA sequence including the $5'$ UTR, CDS, and $3'$ UTR. We use the official implementation and train RiboNN on the transcripts in our training set that have experimental translation efficiency measurements. We tokenize each RNA sequence into codons and feed the tokenized sequence into the RiboNN encoder, which consists of convolutional and recurrent layers as described in [25]. The final hidden representation feeds into a regression head that outputs a single scalar translation efficiency prediction. We optimize RiboNN with MSE loss and use the validation set for early stopping.

**CDS-only.** To compare both models under the same input information, we also train a CDS-only variant of RiboNN. We strip the $5'$ and $3'$ UTR nucleotide sequences from each transcript and provide only the coding sequence as input, setting the UTR lengths to zero. The architecture and all training hyperparameters remain identical to the UTR+CDS setting. This ensures that any difference in performance between `seq2ribo` and RiboNN in the CDS-only setting reflects differences in the models rather than differences in the input information.

**Individual vs. multitask training.** We train RiboNN under two strategies. In the individual setting, we train a separate RiboNN model for each of the four TE tasks (Mean, HEK293, LCL, RPE-1) independently. In the multitask setting, following the original RiboNN framework, we train a single model jointly on all four TE tasks using multitask learning with shared encoder weights and task-specific output heads. Both strategies are evaluated under both input settings (CDS-only and UTR+CDS), yielding four RiboNN configurations in total.

In all settings, we use the same train, validation, and test splits and the same labels that we use for the `seq2ribo` translation efficiency heads so that all models see the same data. At test time, we compute Pearson correlation between predictions and experimental translation efficiency and compare these metrics across all configurations.

##### 1.12 Transformer polisher

To validate the selection of Mamba as the polisher backbone, we perform an ablation study that compares its performance with a standard transformer-based architecture. For this comparison, we train the Mamba-based and transformer-based polishers in an identical simplified setting.

**Dataset.** To expedite training, we use a reduced dataset that contains one sTASEP simulation output per transcript from the training set, the validation set, and the test set. For each transcript $i$, we draw a single simulation from the trained sTASEP simulator $f$ and treat the resulting codon-level counts $\hat{Y}_i = (\hat{y}_i^1, \ldots, \hat{y}_i^{N_i})$ as input to the ablation models.

**Input features.** We restrict the input to non-structural features and exclude the pair, angle, and bucket sequences. For each transcript $i$ and codon position $j$, we construct an input vector

$$x_i^j = \mathrm{Emb}\big(C_i^j\big) + W^{\mathrm{sim}} \, \log\big(1 + \hat{y}_i^j\big),$$

where Emb denotes a learnable codon embedding and $W^{\mathrm{sim}}$ denotes a linear projection that maps the log-transformed simulated A-site count to the shared hidden dimension. Both the Mamba and Transformer models operate on the same sequence of inputs $\{x_i^j\}_{j=1}^{N_i}$.

**Loss function.** We optimize both models with the same simplified two-term loss that combines pertranscriptMAE and elemwiseMAE defined in the evaluation metrics section. We denote these two losses by $\mathcal{L}_{\mathrm{trans}}$ and $\mathcal{L}_{\mathrm{elem}}$ and define

$$\mathcal{L}_{\mathrm{abl1}} = \mathcal{L}_{\mathrm{trans}} + \mathcal{L}_{\mathrm{elem}}.$$

**Transformer baseline architecture.** We implement the Transformer baseline as a standard transformer stack. We first add a sinusoidal positional encoding to the input sequence $\{x_i^j\}$. We then process the encoded sequence with $N = 4$ Transformer encoder layers, each with $H = 8$ attention heads and a feed-forward dimension of 512. A final multilayer perceptron head, identical to the output head in the Mamba polisher, maps the hidden representation at each codon position to a predicted A-site count.

**Mamba baseline architecture.** For the Mamba baseline, we use the same RiboPolisherMamba architecture that we describe in the main Methods section, but we restrict the inputs to codon embeddings and log-transformed sTASEP counts as described above. We train both the Mamba model and the Transformer model until convergence and select the checkpoint with the lowest validation loss $\mathcal{L}_{\mathrm{abl1}}$ for each architecture.

We then evaluate the selected checkpoints on the test set. The Mamba-based polisher achieves a pertranscriptMAE of 0.900 and an elemwiseMAE of 3.951. The Transformer polisher achieves a pertranscriptMAE of 0.945 and an elemwiseMAE of 4.473. The Mamba architecture therefore reduces pertranscriptMAE by 4.8% and elemwiseMAE by 11.7% relative to the Transformer baseline. Based on this head-to-head comparison, we choose Mamba as the core backbone for the `seq2ribo` polisher.

##### 1.13 Training with pertranscriptMAE loss

To isolate the effect of the negative log-likelihood (NLL) loss function that we use for the polisher, we train an ablated version of the `seq2ribo` polisher that optimizes only the pertranscriptMAE. The model architecture, input features, and training data remain the same as in the main polisher experiment. In this ablation, we replace the NLL loss with a single objective. We denote the per transcript mean absolute error by $\mathcal{L}_{\text{trans}}$ and define the ablation loss

$$\mathcal{L}_{\text{abl2}} = \mathcal{L}_{\text{trans}} = \text{pertranscriptMAE}.$$

We update the polisher parameters by backpropagating gradients from this single loss term. We also drive the learning rate scheduler and the early stopping criterion for model checkpointing exclusively by the pertranscriptMAE on the validation set. In particular, we use the validation value of $\mathcal{L}_{\text{trans}}$ as the monitored quantity in the ReduceLROnPlateau scheduler and for model selection. Although we optimize only this single metric, we continue to compute and log all six evaluation metrics at each epoch for both the training and validation sets. At every epoch, we record pertranscriptMAE, elemwiseMAE, codonMAE, pairMAE, angleMAE, and bucketMAE as defined in the evaluation metrics section. This allows us to observe the collateral effects of the narrow optimization strategy on the non-optimized codon-level and structural metrics and yields the training and validation curves shown in Supplementary Figures S1 and S2. Training the polisher on pertranscriptMAE alone successfully drives that specific metric down, but substantially degrades the model's ability to match aggregate structural and codon-level distributions. On the test set, this model achieves a pertranscriptMAE of 0.916 and an elemwiseMAE of 4.128, which are close in magnitude to the corresponding values of the NLL-trained model (0.994 and 4.641, respectively). However, the aggregate metrics collapse. The per codon and structural errors increase by several orders of magnitude: the per codon error reaches a codonMAE of 119,800, the positional error reaches a bucketMAE of 2,555,799, and the structural errors reach a pairMAE of 1,916,849 and an angleMAE of 1,916,849. For comparison, the NLL-trained model attains a codonMAE of 4,570 on the same test set. These results support the use of the NLL loss for polisher training.

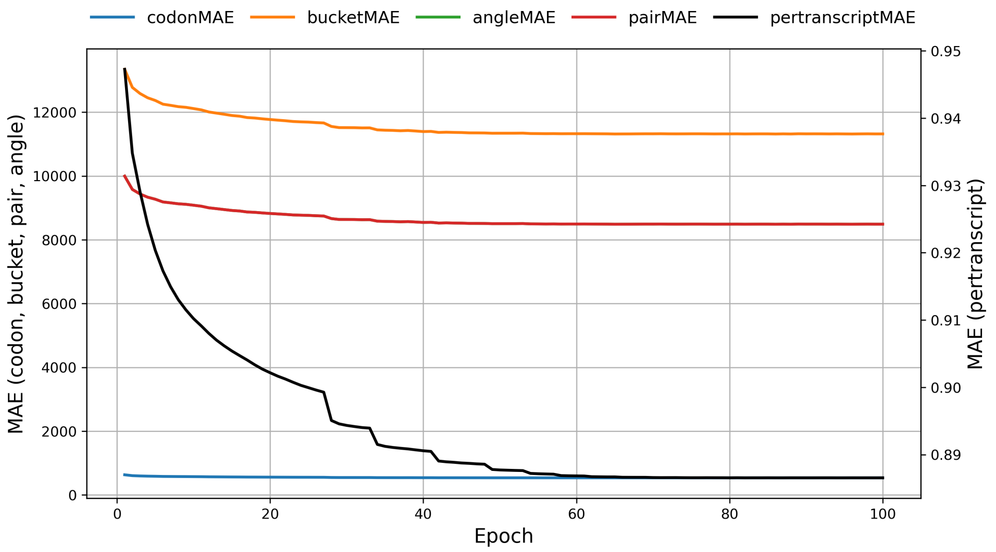


**Figure S1:** Training loss curve of `seq2ribo` polisher trained with $\mathcal{L}_{\text{abl2}}$.

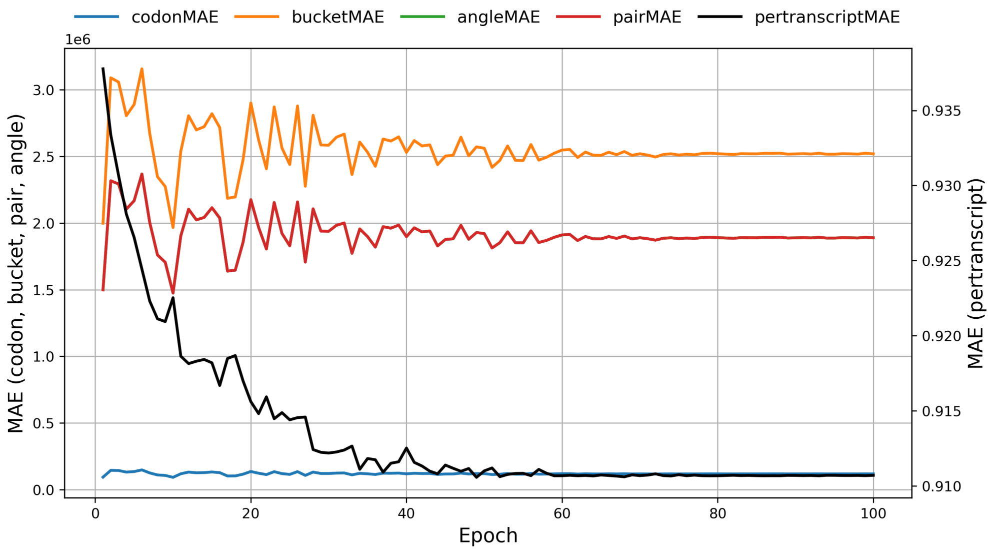


**Figure S2:** Validation loss curve of `seq2ribo` polisher trained with $\mathcal{L}_{\text{abl2}}$.

#### 1.14 Ablation studies on `seq2ribo` components

##### 1.14.1 Sequence and structural features only

To quantify the contribution of the sTASEP simulation as an input feature, we train an ablated polisher that does not receive any simulation counts. In this experiment, we force the model to predict A-site ribosome counts directly from the sequence and its associated structural features.

**Model architecture.** We use an ablated Mamba architecture that is identical to the main `seq2ribo` polisher, except that it does not contain the linear layer that embeds the simulated counts. The output layer uses a Softplus activation to produce non-negative predictions, consistent with the main polisher.

**Input features.** For each transcript $i$ and codon position $j$ we construct the input vector

$$x_i^j = E^{\text{codon}}\big(C_i^j\big) + E^{\text{pair}}\big(p_i^j\big) + E^{\text{angle}}\big(a_i^j\big) + E^{\text{bucket}}\big(b_i^j\big),$$

where $E^{\text{codon}}$, $E^{\text{pair}}$, $E^{\text{angle}}$, and $E^{\text{bucket}}$ denote the same embedding layers that we use in the full polisher. The sTASEP simulation output $\hat{Y}_i$ is completely removed from the input, and the model receives only codon identity and structural features.

**Loss function.** We optimize the ablated model with the same Poisson negative log-likelihood (NLL) loss used for the main polisher. We select the checkpoint with the lowest NLL on the validation set. We additionally compute and log all six evaluation metrics (pertranscriptMAE, elemwiseMAE, codonMAE, pairMAE, angleMAE, and bucketMAE) at each epoch for monitoring purposes.

We train this model until convergence and select the checkpoint with the lowest validation NLL. The training and validation loss curves for this experiment appear in Supplementary Figures S3 and S4. On the test set, the ablated model attains a pertranscriptMAE of 1.30 and an elemwiseMAE of 4.22, compared with the `seq2ribo` model values of 1.39 and 4.14, respectively. The codonMAE is 3,732, the pairMAE is 9,466, the angleMAE is 18,131, and the bucketMAE is 31,270. While the ablated model achieves competitive or even lower MAE values than the full `seq2ribo` model on several metrics, it lags behind the full model in correlation-based metrics, indicating that the sTASEP simulation input contributes primarily to capturing the shape and relative pattern of ribosome density profiles rather than reducing absolute count error.

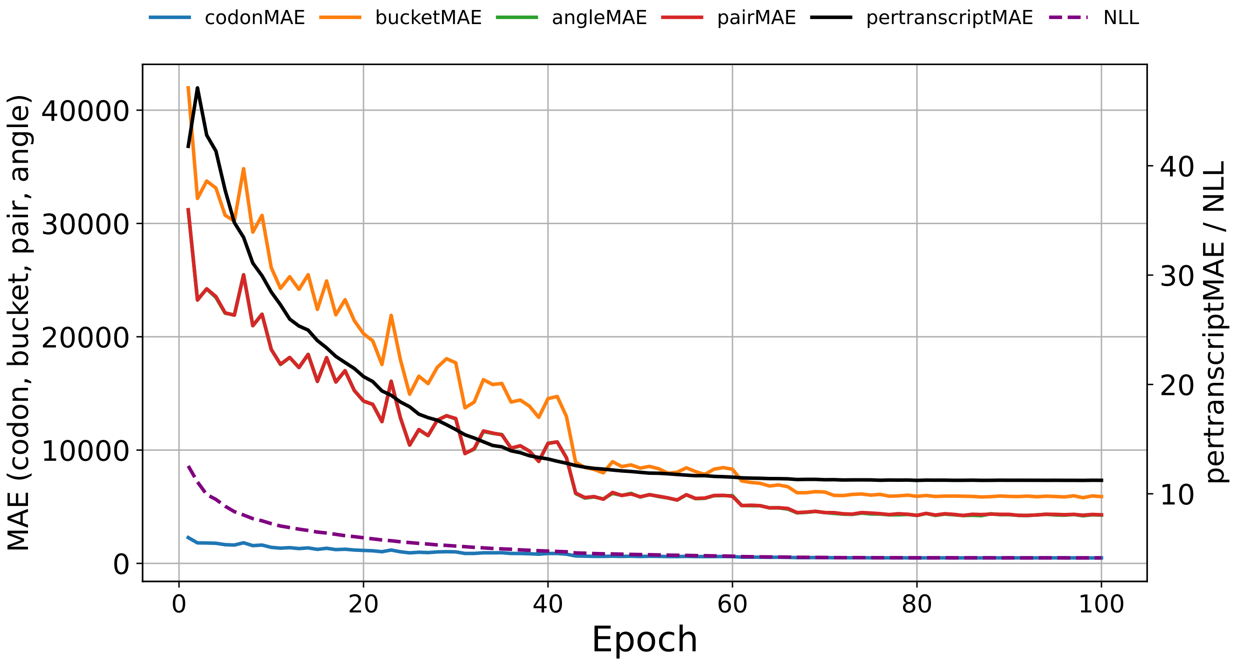


**Figure S3:** Training loss curve during the training of `seq2ribo` polisher trained with sequence and structural features only.

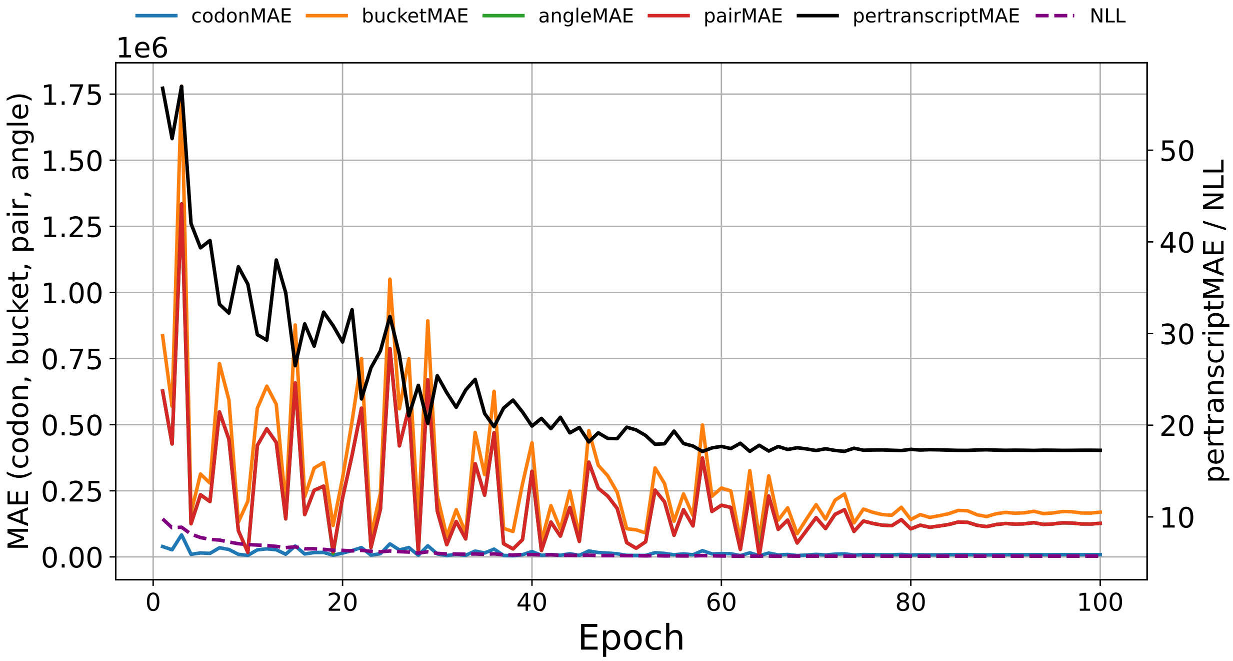


**Figure S4:** Validation loss curve during the training of `seq2ribo` polisher trained with sequence and structural features only.

###### 1.14.2 Sequence and sTASEP simulation only

To isolate the contribution of the static structural features, we train an ablated polisher that uses only the codon sequence and the sTASEP simulation output as inputs and excludes all structural feature embeddings.

**Model architecture.** This model is identical to the `seq2ribo` polisher, but it does not include the structural embedding layers for angle, pair, and bucket features. The output layer uses a Softplus activation to produce non-negative predictions.

**Input features.** For each transcript $i$ and codon position $j$ we construct the input vector

$$x_i^j = E^{\text{codon}}(C_i^j) + W^{\text{sim}} \log(1 + \hat{y}_i^j),$$

where $E^{\text{codon}}$ denotes the codon embedding layer and $W^{\text{sim}}$ denotes the linear projection that embeds the log-transformed sTASEP count at that position. The model does not receive the angle, pair, or bucket features as inputs.

**Loss function.** We optimize the model with the same Poisson negative log-likelihood (NLL) loss used for the main polisher. We select the checkpoint with the lowest NLL on the validation set. We additionally compute and log all six evaluation metrics at each epoch for monitoring purposes. Although the structural features are not inputs to the model, we still compute the structural evaluation metrics $\mathcal{L}_{\text{angle}}$, $\mathcal{L}_{\text{pair}}$, and $\mathcal{L}_{\text{bucket}}$ using the structural maps to assess the model's ability to match structural aggregates even without direct access to the structural annotations.

We train this model until convergence and select the checkpoint with the lowest validation NLL. The training and validation curves for this experiment appear in Supplementary Figures S5 and S6. On the test set, this model achieves a pertranscriptMAE of 1.29 and an elemwiseMAE of 4.13, comparable to the `seq2ribo` model values of 1.39 and 4.14. The codonMAE is 3,548. The aggregate structural metrics are a pairMAE of 19,357, an angleMAE of 24,057, and a bucketMAE of 46,308. Compared with the Seq+Struc. ablation (which includes structural features but omits the sTASEP input), the structural MAE values are higher, indicating that the structural feature embeddings help the model match structural aggregates. Nonetheless, the correlation-based metrics reveal that neither the Seq+sTASEP nor the Seq+Struc. ablation matches the full `seq2ribo` model in capturing the shape of the ribosome density profile, underscoring the complementary value of combining all three feature groups.

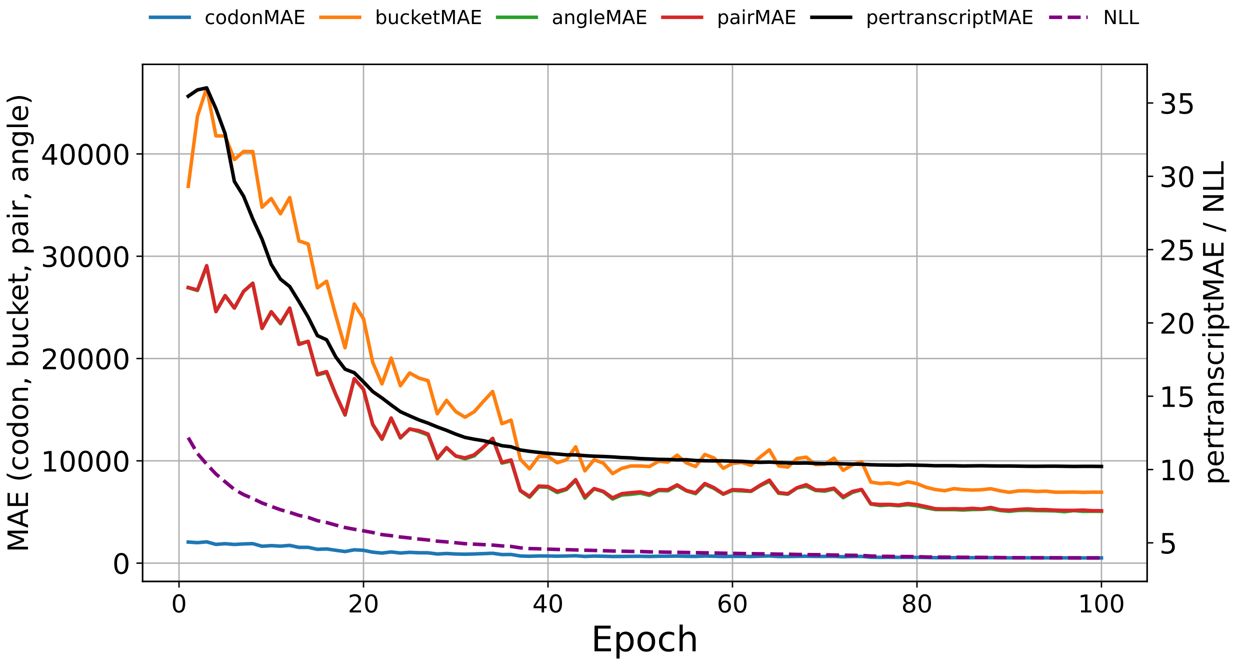


**Figure S5:** Training loss curve during the training of `seq2ribo` polisher trained with sequence and sTASEP simulation only.

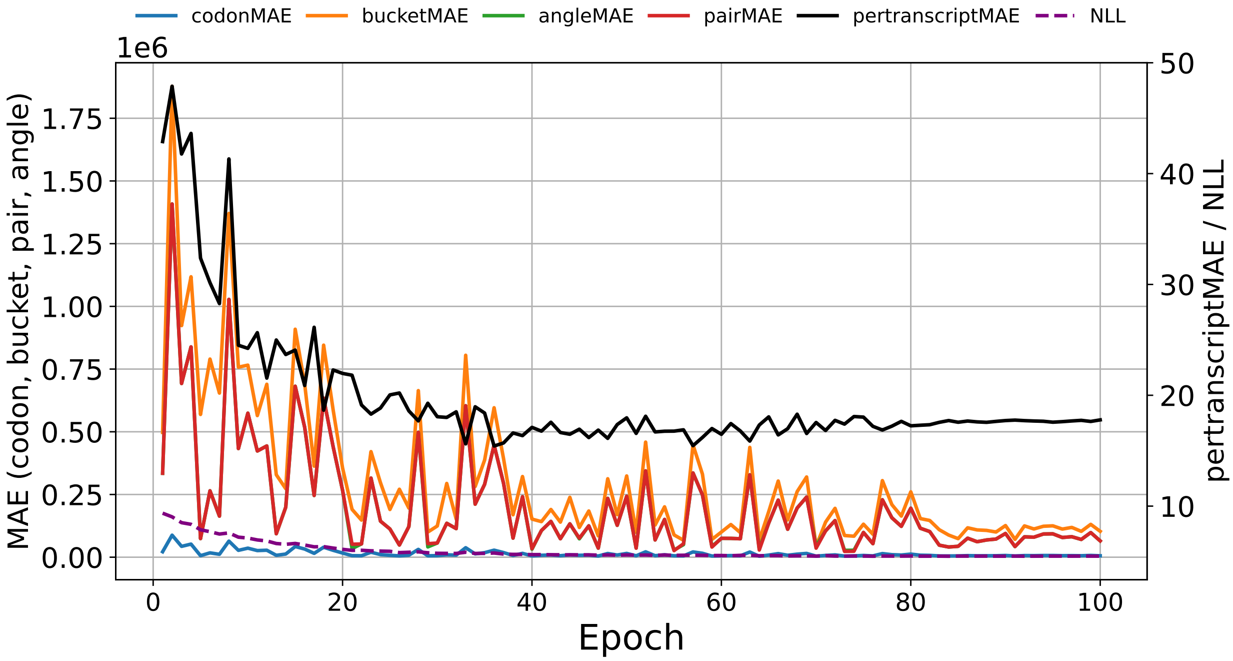


**Figure S6:** Validation loss curve during the training of `seq2ribo` polisher trained with sequence and sTASEP simulation only.

##### 1.14.3 Sequence-only

To isolate the combined contribution of the structural and simulation features, we train an ablated polisher that receives only the codon sequence as input. This experiment provides a sequence-only baseline for ribosome count prediction.

**Model architecture.** This model is identical to the main `seq2ribo` polisher, but it contains only a single embedding layer for codon identity. The output layer uses a Softplus activation.

**Input features.** For each transcript $i$ and codon position $j$ we construct the input vector

$$x_i^j = E^{\text{codon}}(C_i^j),$$

where $E^{\text{codon}}$ is the codon embedding layer. We remove all other inputs and do not provide the sTASEP simulation output or the angle, pair, and bucket features.

**Loss function.** We optimize the sequence-only model with the same Poisson NLL loss used for the main polisher. We select the checkpoint with the lowest NLL on the validation set. We additionally compute and log all six evaluation metrics at each epoch for monitoring purposes.

We train this model until convergence and select the checkpoint with the lowest validation NLL. The training and validation curves for this experiment appear in Supplementary Figures S7 and S8. On the test set, this model attains a pertranscriptMAE of 1.28 and an elemwiseMAE of 3.97, which are the lowest MAE values among all ablations on these two metrics. The codonMAE is 3,555, the pairMAE is 15,509, the angleMAE is 12,497, and the bucketMAE is 68,032. Despite achieving the lowest element-wise and per-transcript MAE, the sequence-only model attains the lowest correlation metrics among the sequence-containing ablations, suggesting that it learns a conservative predictor that minimises absolute error without accurately capturing the positional variation in ribosome density.

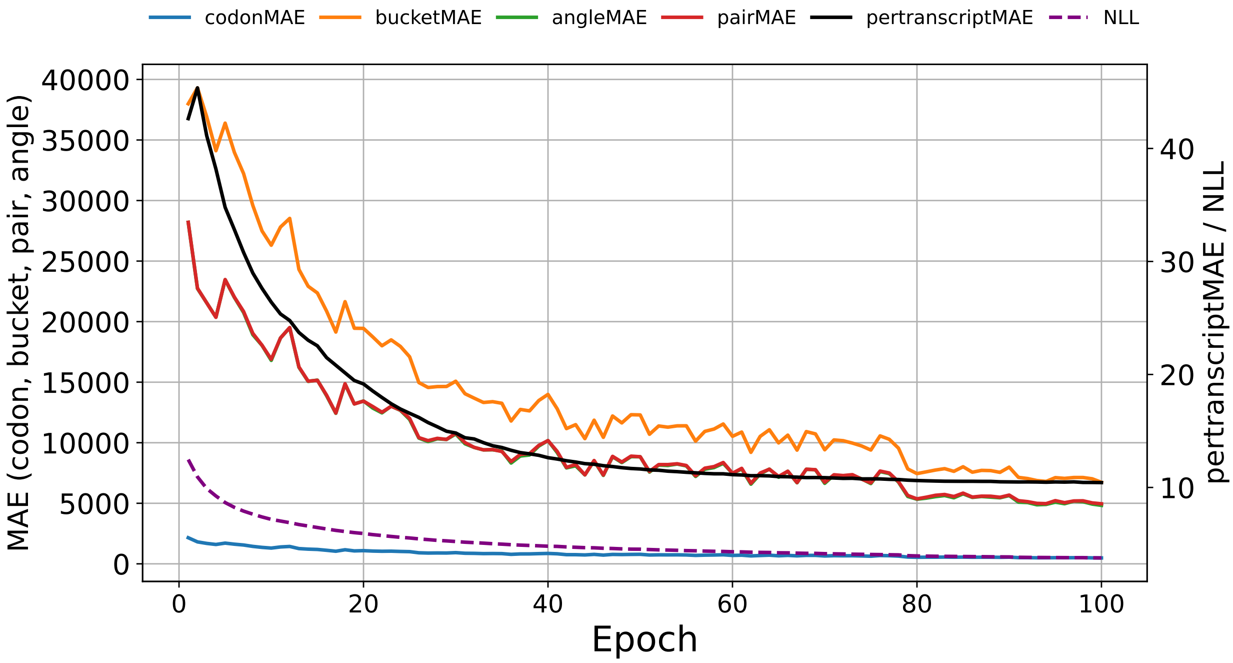


**Figure S7:** Training loss curve during the training of `seq2ribo` polisher trained with sequence-only.

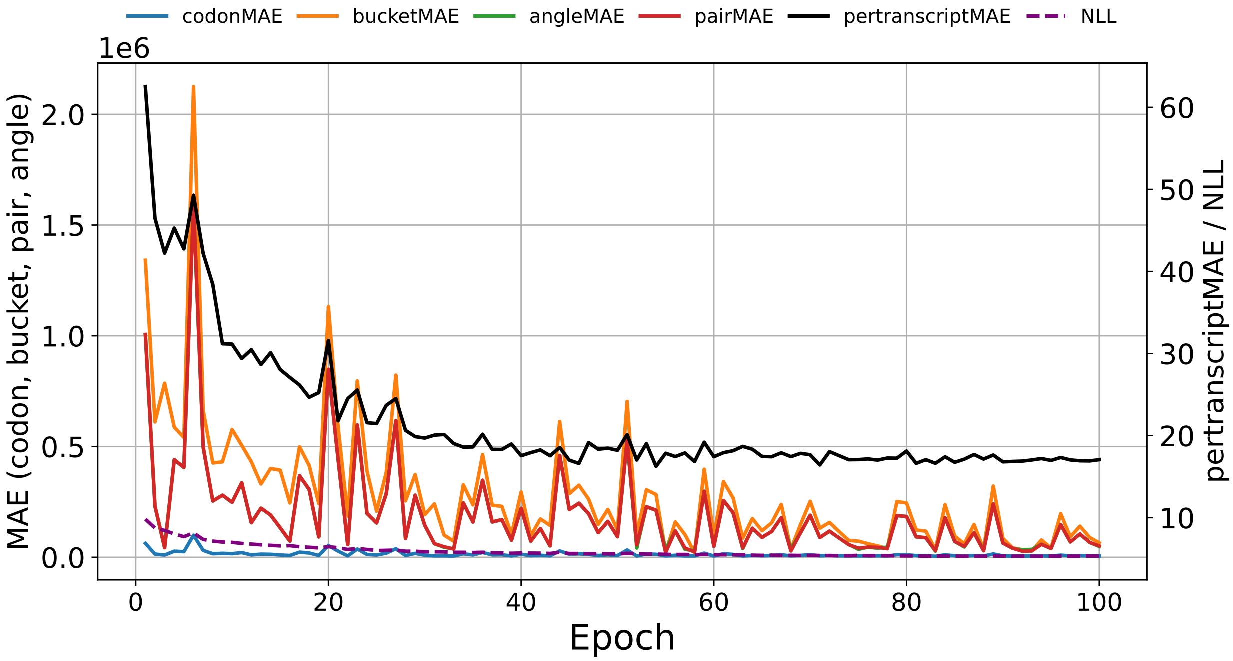


**Figure S8:** Validation loss curve during the training of `seq2ribo` polisher trained with sequence-only.

###### 1.14.4 sTASEP simulation and structural features only

To quantify the contribution of codon identity as an input feature, we train an ablated polisher that does not receive the codon sequence. In this experiment, we force the model to predict A-site ribosome counts using only the sTASEP simulation output and the static structural features.

**Model architecture.** We use an ablated Mamba architecture that is identical to the main `seq2ribo` polisher, except that it does not contain the codon embedding layer. The output layer uses a Softplus activation.

**Input features.** For each transcript $i$ and codon position $j$ we construct the input vector

$$x_i^j = W^{\text{sim}} \log\left(1 + \hat{y}_i^j\right) + E^{\text{pair}}\left(p_i^j\right) + E^{\text{angle}}\left(a_i^j\right) + E^{\text{bucket}}\left(b_i^j\right),$$

where $W^{\text{sim}}$ denotes the same linear projection that embeds the log-transformed sTASEP count in the full polisher, and $E^{\text{pair}}$, $E^{\text{angle}}$, and $E^{\text{bucket}}$ denote the same structural embedding layers that we use in the full model. The codon identity input $C_i^j$ is completely removed from the input, and the model receives only the simulation-derived signal and the structural annotations.

**Loss function.** We optimize the ablated model with the same Poisson NLL loss used for the main polisher. We select the checkpoint with the lowest NLL on the validation set. We additionally compute and log all six evaluation metrics at each epoch for monitoring purposes. Although codon identity is not an input to the model, we still compute codonMAE by aggregating the predicted and observed counts by codon type.

We train this model until convergence and select the checkpoint with the lowest validation NLL. The training and validation loss curves for this experiment appear in Supplementary Figures S9 and S10. On the test set, this model attains a pertranscriptMAE of 1.30 and an elemwiseMAE of 4.51, both higher than the `seq2ribo` model values of 1.39 and 4.14, respectively. The codonMAE is 11,356, the pairMAE is 30,657, the angleMAE is 13,259, and the bucketMAE is 214,407. Correlation-based metrics are substantially lower than those of any sequence-containing ablation: the Tx-level $r$ is 0.258, the Shape $r$ is 0.051, and the Elemwise $r$ is 0.128. Compared with the sTASEP-only ablation, adding structural features improves the Tx-level $r$ from 0.216 to 0.258 and reduces the angleMAE from 21,109 to 13,259, indicating that structural annotations provide a modest but consistent benefit even in the absence of codon identity. Nonetheless, the large gap relative to sequence-containing models confirms that codon identity is the single most informative feature group for ribosome density prediction.

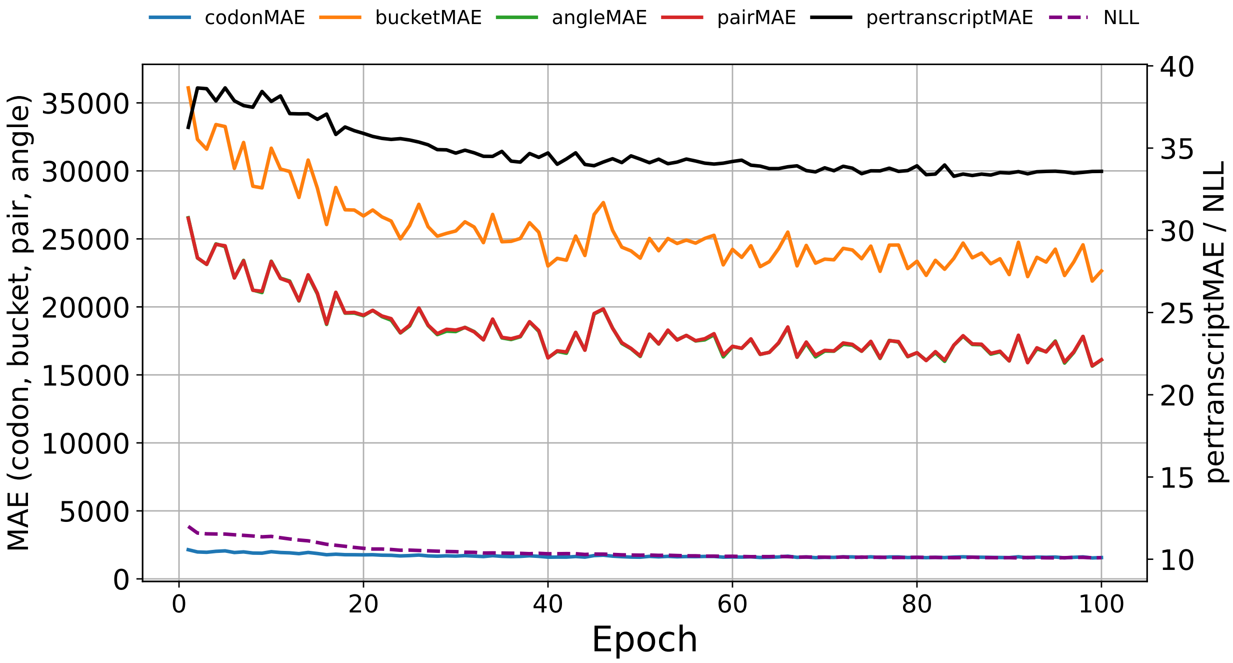


**Figure S9:** Training loss curve during the training of `seq2ribo` polisher trained with sTASEP simulation and structural features only.

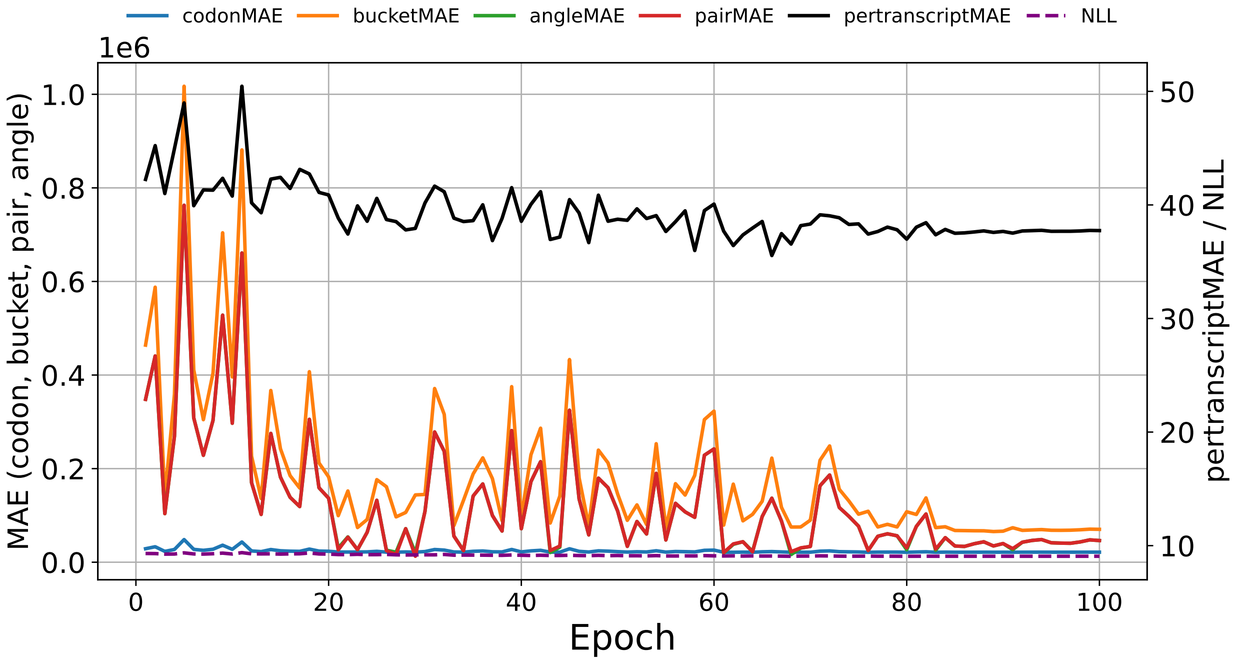


**Figure S10:** Validation loss curve during the training of `seq2ribo` polisher trained with sTASEP simulation and structural features only.

###### 1.14.5 sTASEP simulation only

To isolate the contribution of the static structural features, we train an ablated polisher that receives only the sTASEP simulation output as input and excludes both codon identity and structural feature embeddings.

**Model architecture.** We use an ablated Mamba architecture that is identical to the main `seq2ribo` polisher, except that it contains only the linear layer that embeds the simulated counts and does not include the codon or structural embedding layers. The output layer uses a Softplus activation.

**Input features.** For each transcript $i$ and codon position $j$ we construct the input vector

$$x_i^j = W^{\text{sim}} \log\bigl(1 + \hat{y}_i^j\bigr),$$

where $W^{\text{sim}}$ denotes the same linear projection that embeds the log-transformed sTASEP count in the full polisher. The model does not receive codon identity or the angle, pair, and bucket features as inputs.

**Loss function.** We optimize the model with the same Poisson NLL loss used for the main polisher. We select the checkpoint with the lowest NLL on the validation set. We additionally compute and log all six evaluation metrics at each epoch for monitoring purposes. Although the structural features are not inputs to the model, we still compute the structural evaluation metrics using the structural maps. Similarly, although codon identity is not an input, we still compute codonMAE from per-codon aggregates of predicted and observed counts.

We train this model until convergence and select the checkpoint with the lowest validation NLL. The training and validation curves for this experiment appear in Supplementary Figures S11 and S12. On the test set, this model achieves a pertranscriptMAE of 1.29 and an elemwiseMAE of 4.48. The codonMAE is 11,476, the pairMAE is 22,993, the angleMAE is 21,109, and the bucketMAE is 554,629. The correlation-based metrics are low: the Tx-level $r$ is 0.216, the Shape $r$ is 0.058, and the Elemwise $r$ is 0.102. Although these correlations are modest, the sTASEP-only model outperforms the structure-only ablation on all three correlation metrics (0.216 vs. 0.205, 0.058 vs. 0.026, 0.102 vs. 0.084), indicating that the simulation encodes translational information that is complementary to, and independent of, the structural annotations. The high bucketMAE (554,629) suggests that without codon identity or structural features the model cannot accurately recover structure-level count aggregates, consistent with the expectation that the sTASEP simulation captures positional translational dynamics rather than structural grouping effects.

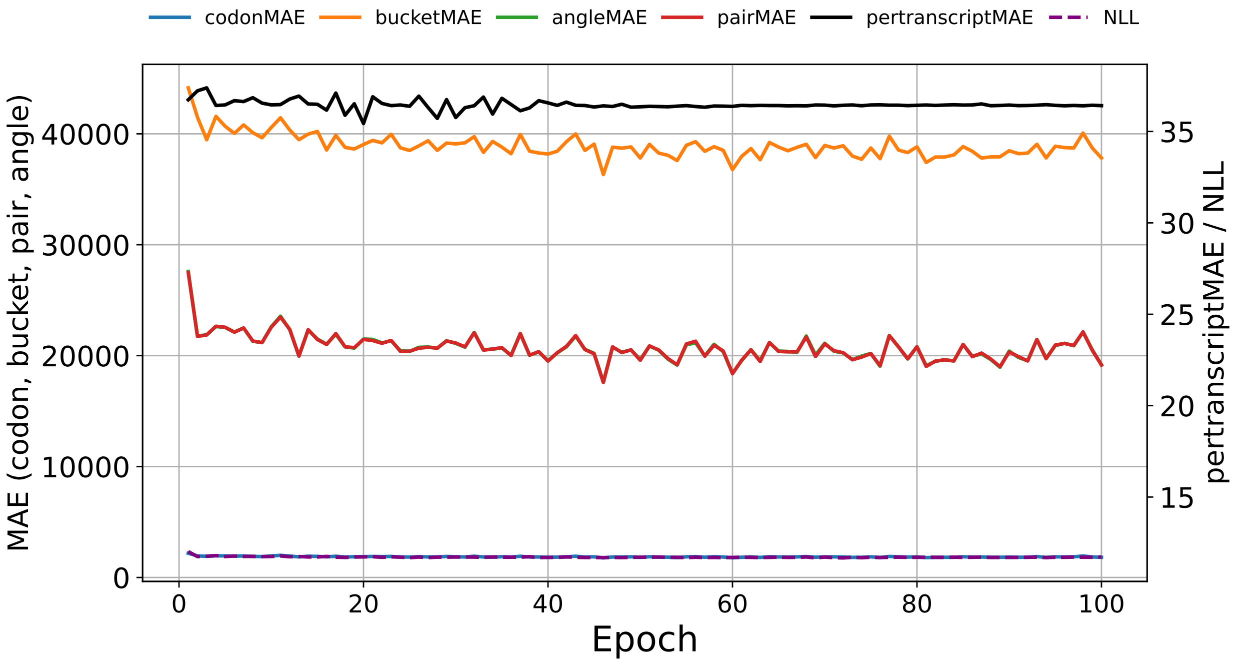


**Figure S11:** Training loss curve during the training of `seq2ribo` polisher trained with sTASEP simulation only.

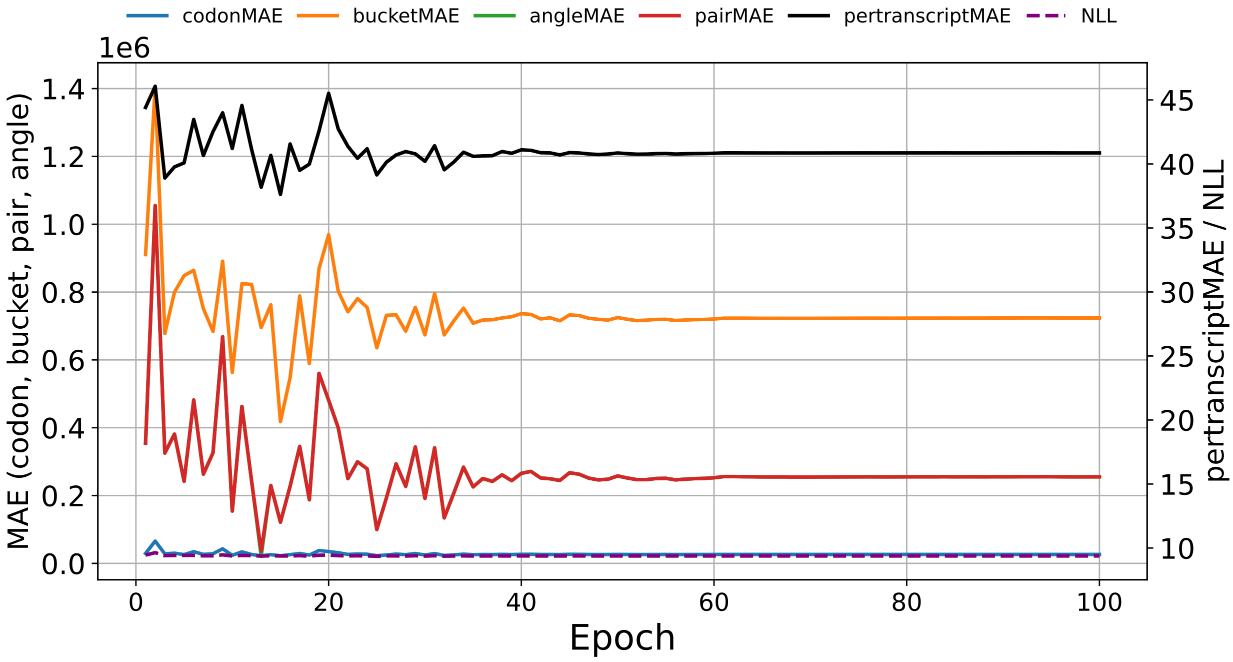


**Figure S12:** Validation loss curve during the training of `seq2ribo` polisher trained with sTASEP simulation only.

###### 1.14.6 Structural features only

To quantify the contribution of the sTASEP simulation output as an input feature in the absence of codon identity, we train an ablated polisher that removes both the codon sequence and the simulation counts and receives only the static structural features. In this experiment, we force the model to predict A-site ribosome counts directly from the structural annotations.

**Model architecture.** We use an ablated Mamba architecture that is identical to the main `seq2ribo` polisher, except that it does not contain the codon embedding layer or the linear layer that embeds the simulated counts. The output layer uses a Softplus activation.

**Input features.** For each transcript $i$ and codon position $j$ we construct the input vector

$$x_i^j = E^{\text{pair}}(p_i^j) + E^{\text{angle}}(a_i^j) + E^{\text{bucket}}(b_i^j),$$

where $E^{\text{pair}}$, $E^{\text{angle}}$, and $E^{\text{bucket}}$ denote the same structural embedding layers that we use in the full polisher. The sTASEP simulation output $\hat{Y}_i$ and the codon identity input $C_i^j$ are completely removed from the input, and the model receives only the structural features.

**Loss function.** We optimize the ablated model with the same Poisson NLL loss used for the main polisher. We select the checkpoint with the lowest NLL on the validation set. We additionally compute and log all six evaluation metrics at each epoch for monitoring purposes. Although codon identity is not an input to the model, we still compute codonMAE by aggregating the predicted and observed counts by codon type.

We train this model until convergence and select the checkpoint with the lowest validation NLL. The training and validation loss curves for this experiment appear in Supplementary Figures S13 and S14. On the test set, this model attains a pertranscriptMAE of 1.32 and an elemwiseMAE of 4.64, the highest values among all ablations on these two metrics. The codonMAE is 14,188, the pairMAE is 14,078, the angleMAE is 20,461, and the bucketMAE is 411,693. The correlation-based metrics are the lowest of any single-feature-group model: the Tx-level $r$ is 0.205, the Shape $r$ is 0.026, and the Elemwise $r$ is 0.084. Notably, although the structure-only model lacks both codon identity and simulation input, it achieves the lowest pairMAE (14,078) among all single-feature-group ablations, reflecting the direct correspondence between the pair structural annotations and the pairMAE evaluation metric. This result confirms that the structural features contribute primarily to matching structure-level count aggregates rather than to capturing the overall shape or positional variation of the ribosome density profile.

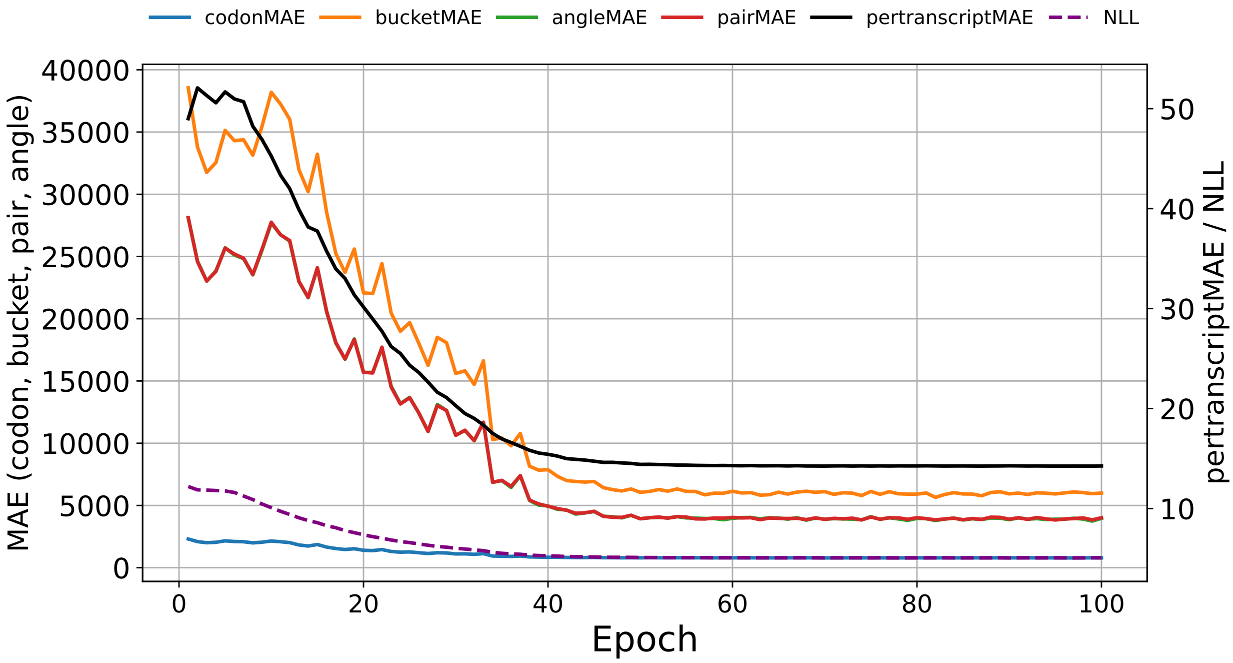


**Figure S13:** Training loss curve during the training of `seq2ribo` polisher trained with structural features only.

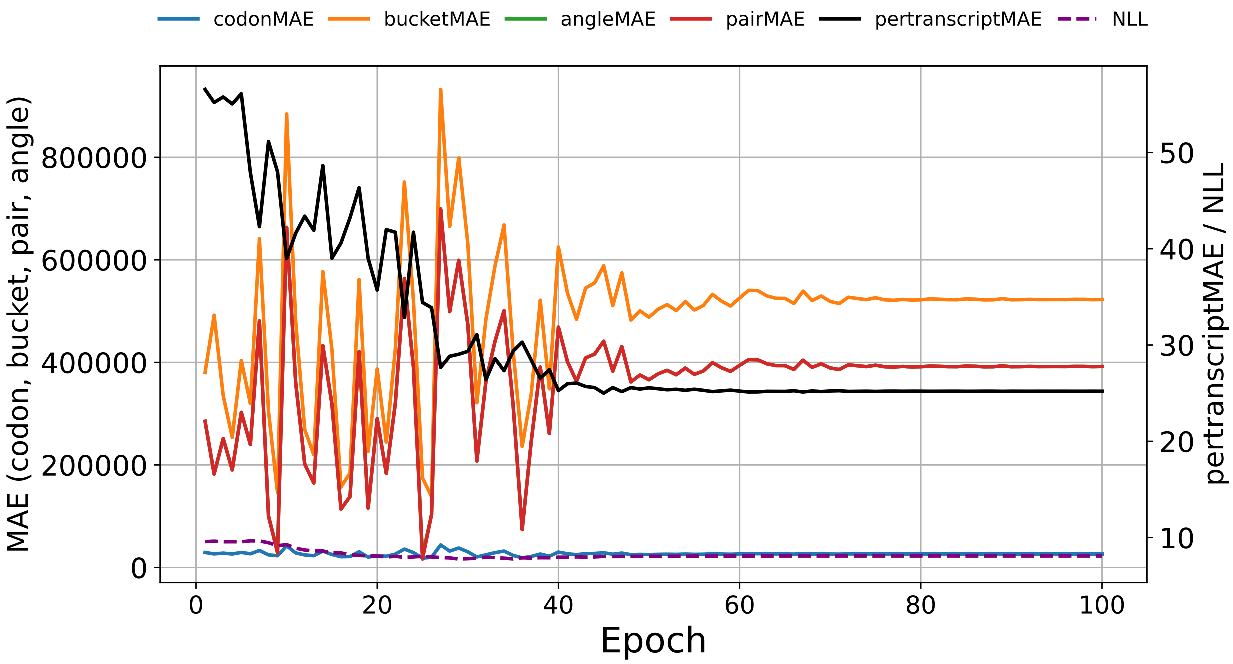


**Figure S14:** Validation loss curve during the training of `seq2ribo` polisher trained with structural features only.

##### 1.15 Translation efficiency prediction with ablated polishers

To assess the contribution of each input component to translation efficiency prediction, we finetune a TE head on top of each of the six ablated polisher variants described above. This extends the ribosome profile ablation to the downstream TE task and allows us to determine whether the same input components that improve profile prediction also improve TE prediction.

**Procedure.** For each ablated polisher (Sequence only, sTASEP only, Structure only, Sequence + sTASEP, Sequence + Structure, sTASEP + Structure), we take the polisher model from the corresponding ribosome profile ablation experiment and attach a TE prediction head. We follow the same finetuning procedure used for the full `seq2ribo` TE head.

**TE head architecture.** The TE head receives the full codon-level count profile produced by the polisher. It applies a position-wise feed-forward network that maps each scalar count through a three-layer MLP with hidden dimensions 256, 128, and 1, using GELU activations. Masked mean pooling over codon positions produces a fixed-length transcript representation, which is passed through a final linear layer and a sigmoid activation to output a scalar prediction in $[0, 1]$.

**TE label transformation.** We apply a min-max transform to map the experimental TE values to the $[0, 1]$ range for training. The transform parameters are computed from the training set and fixed for validation and test evaluation. At test time, predictions are mapped back to the original TE space before computing metrics.

**Training.** We finetune all model parameters (both the ablated polisher base and the TE head) jointly using MSE loss in the scaled $[0, 1]$ space. We use the Adam optimizer with a learning rate of $10^{-4}$, weight decay of $10^{-4}$, gradient clipping at 1.0, and a batch size of 64. We train for 10 epochs and select the best checkpoint by validation MSE in the original TE space. We apply the same CDS-hash-based split filtering to ensure no sequence overlap between splits.

**Results.** The full `seq2ribo` model achieves the highest correlation ($r = 0.688$). Among the two-feature ablations, Seq+sTASEP attains $r = 0.660$, followed by Seq+Struc. at $r = 0.624$ and sTASEP+Struc. at $r = 0.584$. The single-feature models rank as Sequence ($r = 0.654$), sTASEP ($r = 0.578$), and Structure ($r = 0.388$). Several patterns highlight the importance of the sTASEP simulation for TE prediction. First, as a standalone feature the sTASEP simulation achieves $r = 0.578$, substantially outperforming the structure-only baseline ($r = 0.388$) and approaching the sequence-only model ($r = 0.654$), despite encoding no codon identity information whatsoever. This demonstrates that the simulation captures a large portion of the translational dynamics relevant to TE from first principles alone. Second, every model that includes the sTASEP simulation outperforms its counterpart without it: adding sTASEP to sequence features raises the correlation from 0.654 to 0.660, adding it to structural features raises the correlation from 0.388 to 0.584, and the full model ($r = 0.688$) surpasses the Seq+Struc. ablation ($r = 0.624$) by a margin of 0.064. The largest such gain ($0.584 - 0.388 = 0.196$) is observed when sTASEP is added to structural features, confirming that the simulation provides a translational signal. Finally, the full `seq2ribo` model outperforms every ablation, with the margin over the best two-feature combination being $0.688 - 0.660 = 0.028$, demonstrating that all three feature groups contribute non-redundant information and that the hybrid approach is essential for maximizing TE prediction accuracy.

#### 1.16 Evaluation of generalization across sequence similarity

To ensure that `seq2ribo` learns generalizable rules of translation dynamics rather than memorizing specific sequences, we removed from each test split any transcript whose coding sequence was identical to a coding sequence in the training split. This step prevents data leakage by ensuring that identical isoforms or duplicate coding sequences do not appear in both the training and evaluation splits.

To quantify the generalization capability of `seq2ribo` on novel sequences, we computed the pairwise similarity between every transcript in the test set and the entire training corpus. We used a MinHash approximation of the Jaccard similarity index to handle the large dataset size. We encoded each transcript into $k = 5$ codon k-mers and generated MinHash signatures using 64 permutations. We then employed Locality Sensitive Hashing (LSH) to retrieve the nearest neighbor in the training set for each test transcript. The similarity score for a test sample is defined as the maximum estimated Jaccard similarity to any sequence in the training set.

Supplementary Figures S15–S18 display the pertranscriptMAE as a function of this maximum percent similarity. The individual data points (blue) represent test transcripts, and the orange line represents a moving average (window size $N = 401$) across the sorted similarity spectrum.

We observe that the error distribution remains stable across the full range of sequence similarities. The moving average trend line is flat, indicating that predictive accuracy is independent of a sequence's proximity to the training data. Notably, the model maintains high accuracy even on sequences with 0% similarity to the training set, demonstrating its ability to generalize to completely novel genetic contexts. Furthermore, the model does not exhibit a drop in error for the samples with high similarity compared to highly dissimilar sequences. This confirms that `seq2ribo` relies on learned structural and sequence motifs rather than memorization to generate predictions.

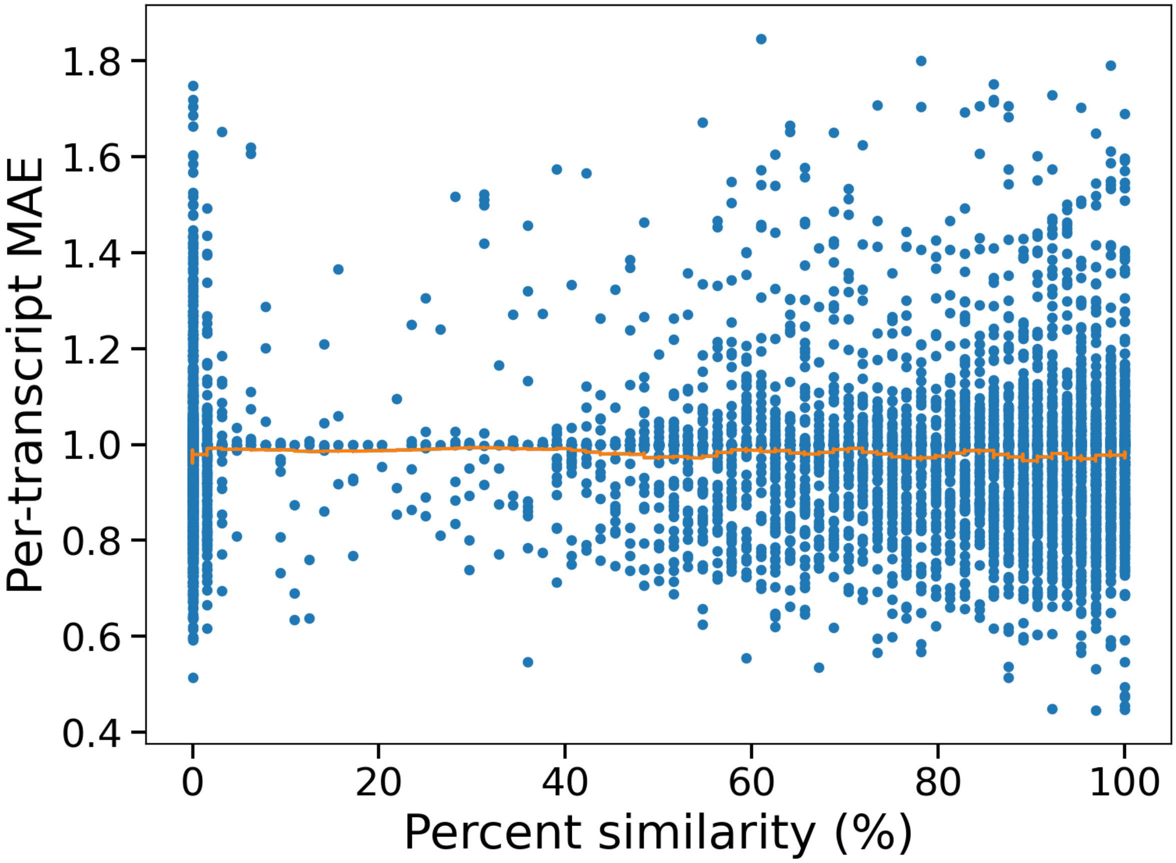


**Figure S15:** pertranscriptMAE versus percent similarity between each iPSC test transcript and its closest training transcript. Points show individual test transcripts. The orange curve shows the mean pertranscript-MAE within similarity bins.

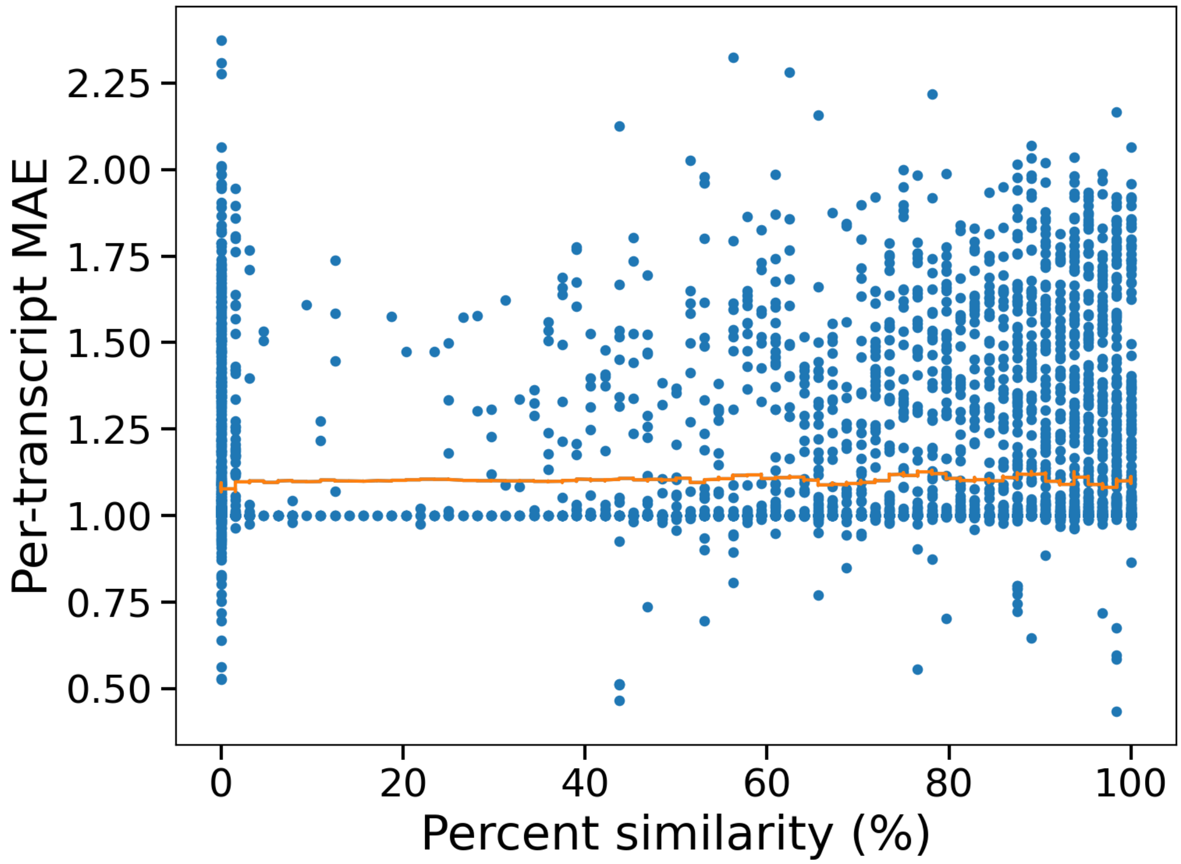


**Figure S16:** pertranscriptMAE versus percent similarity between each HEK293 test transcript and its closest training transcript. Points show individual test transcripts. The orange line indicates the moving average of the error.

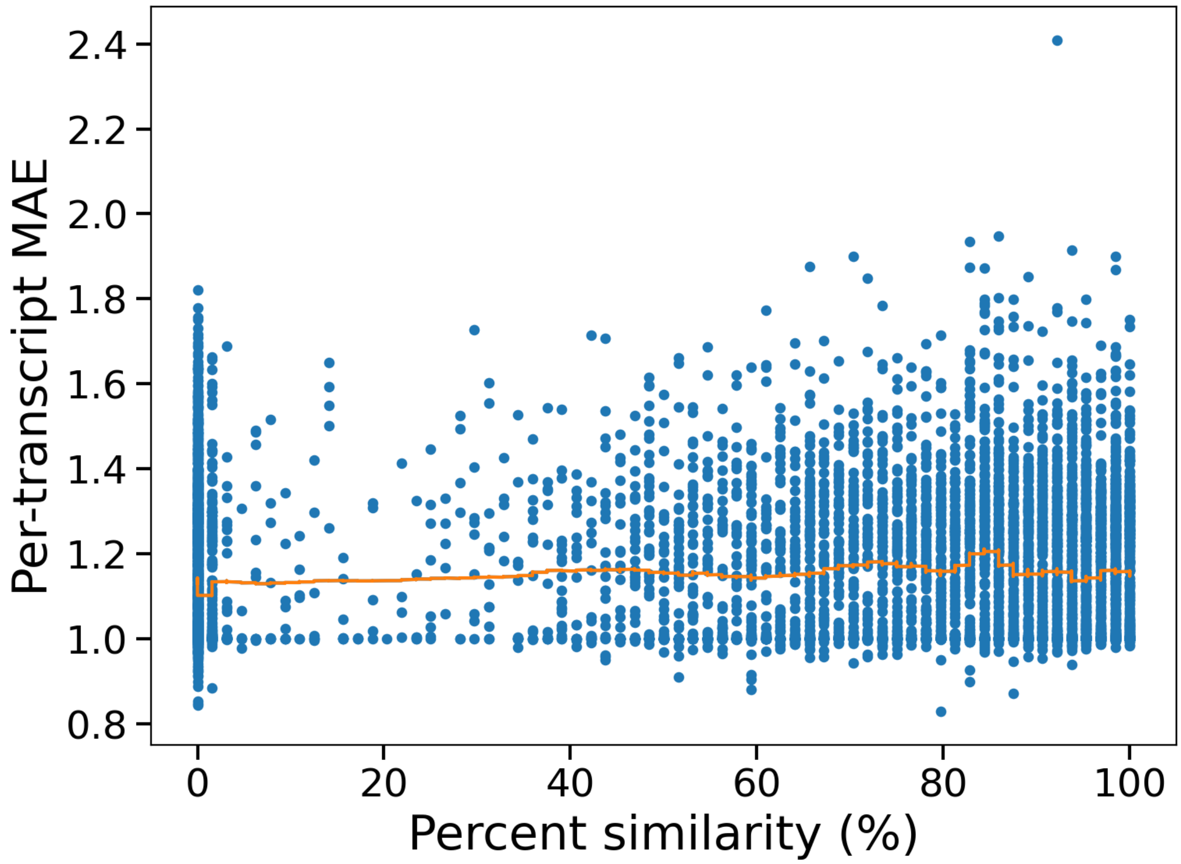


**Figure S17:** pertranscriptMAE versus percent similarity between each LCL test transcript and its closest training transcript. Points show individual test transcripts. The orange line indicates the moving average of the error.

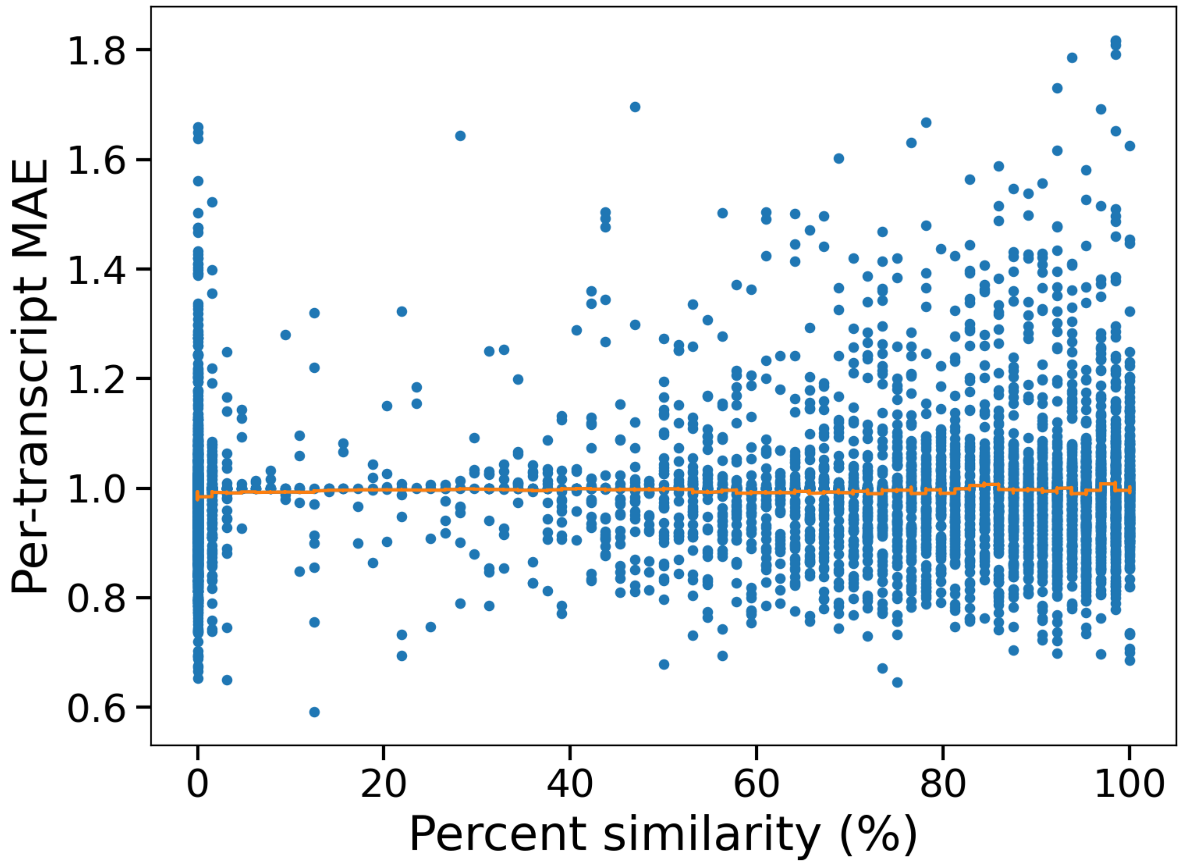


**Figure S18:** pertranscriptMAE versus percent similarity between each RPE-1 test transcript and its closest training transcript. Points show individual test transcripts. The orange line indicates the moving average of the error.

##### 1.17 Robustness of downstream predictions to ribosome profile perturbations

To evaluate the sensitivity of the finetuned TE and expression heads to perturbations in the predicted ribosome count profiles, we design three controlled perturbation experiments. In each experiment, we take the polished ribosome profiles produced by `seq2ribo` on the test set and apply a systematic perturbation before feeding the modified profiles into the downstream prediction heads. We report Pearson correlation between the perturbed predictions and the ground-truth labels.

**Gaussian noise.** We add independent Gaussian noise $\mathcal{N}(0, \sigma^2)$ to each codon-level count in the polished profile and clip negative values to zero. We evaluate eight noise levels: $\sigma \in \{0, 0.1, 0.25, 0.5, 1, 2, 5, 10\}$. As shown in Table S1, TE predictions are broadly robust across all cell lines, with most showing negligible degradation up to $\sigma = 5$. At the most extreme noise level ($\sigma = 10$), most cell lines exhibit only modest declines—iPSC drops from 0.688 to 0.669 and LCL remains essentially unchanged—although HEK293 shows a more pronounced decrease (from 0.570 to 0.517, a decline of $\sim$0.05). Expression predictions show greater sensitivity at high noise levels but remain remarkably stable at moderate perturbations: all cell lines retain correlations within 0.01 of their unperturbed values up to $\sigma = 2$. At $\sigma = 10$, RPE-1 expression shows the largest absolute decline (from 0.830 to 0.750), while LCL is the most resilient (from 0.873 to 0.856). Overall, these results indicate that the expression head relies more heavily on precise count magnitudes than the TE head, but both tolerate substantial noise before performance degrades meaningfully.

**Random dropout.** We randomly set a fraction of codon-level counts to zero, simulating missing or unreliable positions. We evaluate dropout fractions ranging from 0% to 90%. As shown in Table S2, TE predictions degrade gracefully, retaining correlations above 0.5 even at 70% dropout in iPSC and LCL. Expression predictions are substantially more sensitive: at 50% dropout, Pearson correlation drops to roughly 60–70% of the unperturbed value across cell lines (e.g., HEK293 from 0.883 to 0.573; RPE-1 from 0.830 to 0.358). This asymmetry suggests that the expression head depends on dense positional coverage: dropout creates zero-valued positions that dilute the mean-pooled transcript representation, whereas the TE head captures relative distributional features that are more resilient to sparse missing values.

**Uniform scaling.** We multiply all codon-level counts by a constant scale factor $s \in \{0.1, 0.25, 0.5, 0.8, 1.0, 1.2, 1.5, 2.0, 5.0, 10.0\}$. As shown in Table S3, TE predictions are largely invariant to scaling within the range 0.5×–2.0×, with performance changing by less than 0.02 in most cell lines. This invariance is expected because TE reflects translational efficiency per mRNA molecule rather than absolute ribosome load. However, at extreme upward scaling (5×–10×), HEK293 TE degrades sharply (from 0.570 to 0.419 and 0.202, respectively), suggesting that very large uniform amplification pushes count magnitudes into a regime where the head's learned mapping breaks down. The remaining cell lines show milder declines at these extremes. Expression predictions are similarly robust across all four cell lines, with Pearson correlations varying by no more than 0.005 across the full range of scale factors for iPSC, LCL, and RPE-1, and by less than 0.005 for HEK293. This near-complete invariance to uniform scaling indicates that the expression head has learned to rely primarily on the shape and relative distribution of ribosome density across codons rather than on absolute count magnitudes.

**Table S1:** Pearson correlation under varying levels of Gaussian noise ($\sigma$).

| Cell Line | Task | $\sigma = 0$ | $\sigma = 0.1$ | $\sigma = 0.25$ | $\sigma = 0.5$ | $\sigma = 1$ | $\sigma = 2$ | $\sigma = 5$ | $\sigma = 10$ |
|---|---|---|---|---|---|---|---|---|---|
| HEK293 | TE | 0.570 | 0.570 | 0.570 | 0.570 | 0.570 | 0.569 | 0.557 | 0.517 |
| | Expression | 0.883 | 0.883 | 0.883 | 0.882 | 0.882 | 0.878 | 0.854 | 0.779 |
| iPSC | TE | 0.688 | 0.688 | 0.688 | 0.688 | 0.688 | 0.686 | 0.681 | 0.669 |
| | Expression | 0.903 | 0.903 | 0.903 | 0.903 | 0.902 | 0.900 | 0.884 | 0.832 |
| LCL | TE | 0.697 | 0.697 | 0.697 | 0.697 | 0.697 | 0.697 | 0.697 | 0.697 |
| | Expression | 0.873 | 0.873 | 0.873 | 0.873 | 0.873 | 0.872 | 0.869 | 0.856 |
| RPE-1 | TE | 0.616 | 0.616 | 0.616 | 0.616 | 0.616 | 0.614 | 0.604 | 0.585 |
| | Expression | 0.830 | 0.830 | 0.829 | 0.829 | 0.828 | 0.825 | 0.805 | 0.750 |

**Table S2:** Pearson correlation under varying dropout fractions.

| Cell Line | Task | 0% | 5% | 10% | 20% | 30% | 50% | 70% | 90% |
|---|---|---|---|---|---|---|---|---|---|
| HEK293 | TE | 0.570 | 0.568 | 0.565 | 0.557 | 0.548 | 0.530 | 0.499 | 0.379 |
| | Expression | 0.883 | 0.853 | 0.822 | 0.762 | 0.700 | 0.573 | 0.432 | 0.245 |
| iPSC | TE | 0.688 | 0.685 | 0.682 | 0.679 | 0.675 | 0.658 | 0.625 | 0.502 |
| | Expression | 0.903 | 0.876 | 0.848 | 0.791 | 0.732 | 0.606 | 0.461 | 0.263 |
| LCL | TE | 0.697 | 0.694 | 0.692 | 0.686 | 0.681 | 0.663 | 0.624 | 0.499 |
| | Expression | 0.873 | 0.846 | 0.819 | 0.764 | 0.707 | 0.585 | 0.445 | 0.254 |
| RPE-1 | TE | 0.616 | 0.615 | 0.613 | 0.607 | 0.598 | 0.576 | 0.546 | 0.451 |
| | Expression | 0.830 | 0.747 | 0.680 | 0.573 | 0.488 | 0.358 | 0.247 | 0.133 |

**Table S3:** Pearson correlation under varying input scale factors.

| Cell Line | Task | 0.1× | 0.25× | 0.5× | 0.8× | 1.0× | 1.2× | 1.5× | 2.0× | 5.0× | 10× |
|---|---|---|---|---|---|---|---|---|---|---|---|
| HEK293 | TE | 0.555 | 0.560 | 0.562 | 0.566 | 0.570 | 0.574 | 0.577 | 0.576 | 0.419 | 0.202 |
| | Expression | 0.878 | 0.882 | 0.883 | 0.883 | 0.883 | 0.883 | 0.882 | 0.882 | 0.882 | 0.882 |
| iPSC | TE | 0.645 | 0.672 | 0.683 | 0.687 | 0.688 | 0.689 | 0.689 | 0.688 | 0.684 | 0.680 |
| | Expression | 0.903 | 0.904 | 0.904 | 0.903 | 0.903 | 0.903 | 0.903 | 0.903 | 0.903 | 0.903 |
| LCL | TE | 0.685 | 0.693 | 0.696 | 0.697 | 0.697 | 0.697 | 0.697 | 0.697 | 0.695 | 0.693 |
| | Expression | 0.872 | 0.873 | 0.873 | 0.873 | 0.873 | 0.873 | 0.873 | 0.873 | 0.873 | 0.873 |
| RPE-1 | TE | 0.565 | 0.593 | 0.609 | 0.615 | 0.616 | 0.617 | 0.617 | 0.617 | 0.612 | 0.606 |
| | Expression | 0.828 | 0.829 | 0.829 | 0.829 | 0.830 | 0.830 | 0.830 | 0.830 | 0.830 | 0.830 |

##### 1.18 Stratified performance of ribosome profile prediction by translation efficiency

To understand how ribosome profile prediction quality varies with translational activity, we partition the test transcripts in each cell line into ten equal-sized bins ordered by increasing translation efficiency. We evaluate all eight metrics within each bin. Results are shown in Tables S4–S7.

Several consistent trends emerge across cell lines. First, pertranscriptMAE decreases with increasing TE in all four datasets. This pattern indicates that highly translated transcripts, which carry more ribosome footprints, are easier for `seq2ribo` to model, likely because the higher count signal provides a better target for the polisher to learn from. Second, elemwiseMAE increases sharply in the highest TE bins, reflecting the larger absolute counts associated with highly translated transcripts. Third, positional correlation metrics (Shape $r$ and Elemwise $r$) tend to improve in higher-TE bins, particularly in iPSC and LCL, suggesting that `seq2ribo` more accurately captures within-transcript ribosome positioning when transcripts have abundant signal. Fourth, the structural metrics (codonMAE, pairMAE, angleMAE, bucketMAE) generally increase with TE, which is expected because these metrics are computed in absolute terms and scale with the magnitude of ribosome counts.

Taken together, the stratified analysis shows that `seq2ribo` provides its most reliable normalized predictions for highly translated transcripts and that prediction difficulty, in terms of both absolute error and positional accuracy, varies systematically with translational activity.

**Table S4:** Performance of `seq2ribo` across translation efficiency-level bins (iPSC test set). Bins are ordered by increasing translation efficiency.

| | **Bin 0** ($n$=74) | **Bin 1** ($n$=73) | **Bin 2** ($n$=73) | **Bin 3** ($n$=73) | **Bin 4** ($n$=73) | **Bin 5** ($n$=74) | **Bin 6** ($n$=73) | **Bin 7** ($n$=73) | **Bin 8** ($n$=73) | **Bin 9** ($n$=74) |
|---|---|---|---|---|---|---|---|---|---|---|
| pertranscriptMAE | 1.615 | 1.553 | 1.431 | 1.397 | 1.375 | 1.166 | 1.226 | 1.103 | 1.024 | 1.166 |
| elemwiseMAE | 1.489 | 2.227 | 2.695 | 2.798 | 3.544 | 5.495 | 6.896 | 10.384 | 13.808 | 20.871 |
| Shape $r$ | 0.066 | 0.096 | 0.098 | 0.117 | 0.090 | 0.183 | 0.155 | 0.209 | 0.244 | 0.219 |
| Tx-level $r$ | 0.548 | 0.441 | 0.858 | 0.534 | 0.800 | 0.744 | 0.674 | 0.579 | 0.851 | 0.748 |
| Elemwise $r$ | 0.120 | 0.225 | 0.343 | 0.173 | 0.225 | 0.257 | 0.292 | 0.239 | 0.610 | 0.354 |
| codonMAE | 4,112 | 5,534 | 3,662 | 3,945 | 4,109 | 5,446 | 5,567 | 7,227 | 5,542 | 9,156 |
| pairMAE | 17,767 | 20,276 | 5,923 | 9,613 | 15,176 | 28,438 | 30,648 | 69,601 | 14,272 | 46,402 |
| angleMAE | 17,069 | 33,349 | 14,264 | 18,753 | 15,174 | 51,830 | 18,488 | 35,022 | 24,814 | 33,301 |
| bucketMAE | 106,231 | 138,683 | 65,485 | 105,083 | 114,925 | 137,558 | 130,931 | 55,589 | 32,848 | 95,529 |

**Table S5:** Performance of `seq2ribo` across translation efficiency-level bins (HEK293 test set). Bins are ordered by increasing translation efficiency.

| | **Bin 0** ($n$=71) | **Bin 1** ($n$=71) | **Bin 2** ($n$=71) | **Bin 3** ($n$=71) | **Bin 4** ($n$=70) | **Bin 5** ($n$=71) | **Bin 6** ($n$=71) | **Bin 7** ($n$=71) | **Bin 8** ($n$=71) | **Bin 9** ($n$=71) |
|---|---|---|---|---|---|---|---|---|---|---|
| pertranscriptMAE | 1.983 | 1.980 | 1.978 | 1.960 | 1.970 | 1.975 | 1.954 | 1.937 | 1.929 | 1.702 |
| elemwiseMAE | 0.067 | 0.081 | 0.110 | 0.121 | 0.164 | 0.223 | 0.265 | 0.447 | 0.652 | 1.920 |
| Shape $r$ | 0.006 | 0.019 | 0.015 | 0.029 | 0.022 | 0.025 | 0.050 | 0.046 | 0.034 | 0.162 |
| Tx-level $r$ | −0.051 | −0.019 | 0.010 | 0.733 | 0.005 | 0.034 | 0.023 | 0.689 | 0.211 | 0.338 |
| Elemwise $r$ | 0.013 | 0.005 | 0.027 | 0.434 | 0.052 | 0.014 | 0.050 | 0.146 | 0.135 | 0.285 |
| codonMAE | 475 | 613 | 644 | 712 | 733 | 1,016 | 1,138 | 1,053 | 2,131 | 2,881 |
| pairMAE | 2,228 | 4,855 | 2,045 | 3,491 | 3,212 | 3,839 | 3,935 | 4,565 | 7,111 | 13,562 |
| angleMAE | 3,571 | 3,488 | 4,851 | 3,361 | 4,810 | 7,580 | 5,564 | 6,250 | 13,906 | 14,702 |
| bucketMAE | 8,131 | 10,449 | 11,348 | 11,015 | 13,512 | 18,086 | 19,526 | 20,474 | 32,810 | 39,578 |

**Table S6:** Performance of `seq2ribo` across translation efficiency-level bins (LCL test set). Bins are ordered by increasing translation efficiency.

| | **Bin 0** ($n$=71) | **Bin 1** ($n$=70) | **Bin 2** ($n$=71) | **Bin 3** ($n$=70) | **Bin 4** ($n$=71) | **Bin 5** ($n$=70) | **Bin 6** ($n$=71) | **Bin 7** ($n$=70) | **Bin 8** ($n$=71) | **Bin 9** ($n$=71) |
|---|---|---|---|---|---|---|---|---|---|---|
| pertranscriptMAE | 1.590 | 1.565 | 1.452 | 1.463 | 1.404 | 1.432 | 1.371 | 1.440 | 1.328 | 1.364 |
| elemwiseMAE | 4.172 | 5.452 | 14.286 | 19.006 | 19.157 | 22.518 | 28.968 | 48.787 | 70.934 | 184.857 |
| Shape $r$ | 0.104 | 0.120 | 0.137 | 0.153 | 0.143 | 0.178 | 0.220 | 0.122 | 0.241 | 0.226 |
| Tx-level $r$ | 0.457 | 0.565 | 0.799 | 0.853 | 0.735 | 0.803 | 0.940 | 0.847 | 0.932 | 0.355 |
| Elemwise $r$ | 0.133 | 0.270 | 0.330 | 0.212 | 0.377 | 0.344 | 0.473 | 0.364 | 0.474 | 0.416 |
| codonMAE | 5,545 | 5,915 | 16,066 | 16,569 | 10,891 | 14,429 | 15,206 | 28,409 | 39,875 | 100,898 |
| pairMAE | 28,599 | 11,948 | 41,283 | 130,214 | 9,726 | 24,175 | 68,941 | 132,239 | 140,548 | 523,374 |
| angleMAE | 10,829 | 28,475 | 43,806 | 95,459 | 49,524 | 72,053 | 33,863 | 63,779 | 197,485 | 257,313 |
| bucketMAE | 178,610 | 172,444 | 533,379 | 359,499 | 259,841 | 318,299 | 158,544 | 604,346 | 45,993 | 329,635 |

**Table S7:** Performance of `seq2ribo` across translation efficiency-level bins (RPE-1 test set). Bins are ordered by increasing translation efficiency.

| | **Bin 0** ($n$=68) | **Bin 1** ($n$=67) | **Bin 2** ($n$=67) | **Bin 3** ($n$=67) | **Bin 4** ($n$=67) | **Bin 5** ($n$=67) | **Bin 6** ($n$=67) | **Bin 7** ($n$=67) | **Bin 8** ($n$=67) | **Bin 9** ($n$=68) |
|---|---|---|---|---|---|---|---|---|---|---|
| pertranscriptMAE | 1.732 | 1.697 | 1.558 | 1.459 | 1.469 | 1.410 | 1.296 | 1.302 | 1.284 | 1.225 |
| elemwiseMAE | 0.801 | 1.376 | 2.058 | 1.882 | 2.569 | 3.298 | 3.835 | 5.294 | 9.700 | 57.766 |
| Shape $r$ | 0.071 | 0.043 | 0.090 | 0.135 | 0.124 | 0.079 | 0.134 | 0.134 | 0.136 | 0.250 |
| Tx-level $r$ | 0.324 | 0.222 | 0.481 | 0.642 | 0.660 | 0.835 | 0.758 | 0.790 | 0.704 | 0.142 |
| Elemwise $r$ | 0.145 | 0.100 | 0.177 | 0.320 | 0.184 | 0.241 | 0.366 | 0.317 | 0.374 | 0.200 |
| codonMAE | 1,379 | 4,107 | 2,938 | 2,352 | 3,318 | 3,989 | 2,946 | 2,998 | 5,183 | 63,807 |
| pairMAE | 8,654 | 18,614 | 12,433 | 4,880 | 7,230 | 18,288 | 7,816 | 2,843 | 18,467 | 248,165 |
| angleMAE | 8,317 | 10,313 | 11,630 | 7,058 | 6,227 | 18,986 | 11,972 | 5,327 | 15,199 | 179,606 |
| bucketMAE | 58,738 | 113,642 | 81,131 | 63,226 | 67,696 | 97,477 | 47,807 | 90,879 | 115,208 | 1,058,448 |

##### 1.19 Stratified ribosome profile prediction by protein expression

To assess whether the predicted ribosome profiles capture information relevant to protein expression prior to any expression-specific finetuning, we stratify the polisher's predicted total ribosome load by protein expression tertile on the mRFP test set. For each cell-line polisher, we bin the 219 test transcripts into three equal-sized groups ordered by increasing measured expression and compute the mean predicted ribosome load per bin.

As shown in Figure S19, the LCL polisher shows a strong monotonic increase in predicted load with expression (tertile means: 3,941 to 8,777; Spearman $\rho = 1.0$), indicating that the polisher trained only on Ribo-seq data already assigns higher ribosome occupancy to sequences that produce more protein. The iPSC and HEK293 polishers show the same monotonic trend (921 to 940 and 253 to 260, respectively; both $\rho = 1.0$), though with a much narrower range. The RPE-1 polisher increases from the lowest to the middle tertile (717 to 902) but does not increase further for the highest tertile (894; $\rho = 0.5$, $p = 0.67$). The raw sTASEP input is nearly constant across all bins and cell lines ($\sim$ 1,039–1,096), confirming that this signal comes from the polisher rather than the simulator. These patterns are broadly consistent with the downstream finetuning results, where the LCL and iPSC polishers achieve the highest expression prediction correlations ($r = 0.873$ and 0.903).

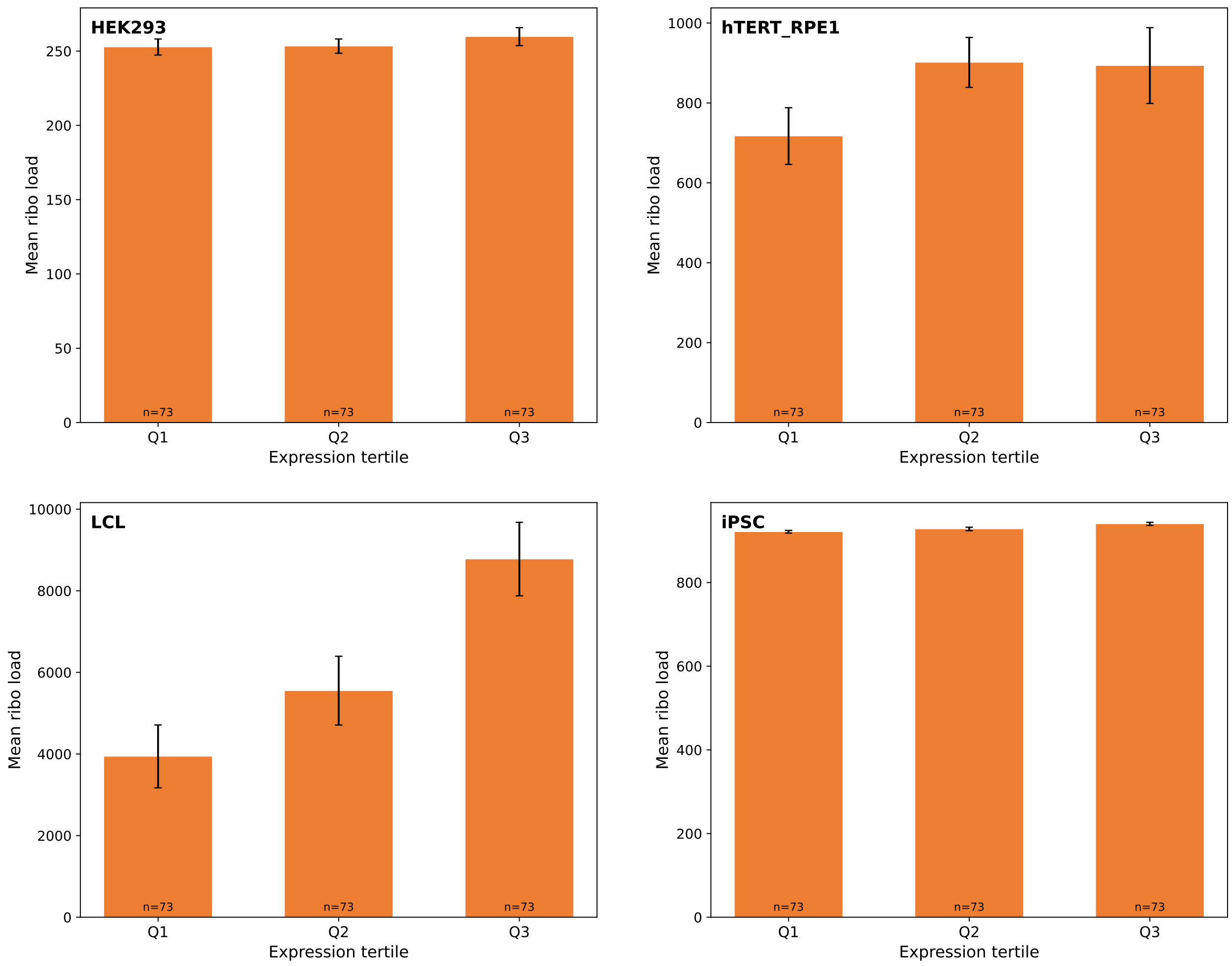


**Figure S19:** Mean predicted ribosome load from the `seq2ribo` polisher (without expression-specific finetuning) stratified by protein expression tertile on the mRFP test set. Each panel shows one cell-line polisher. Error bars indicate standard error of the mean. The LCL polisher shows a strong monotonic increase, indicating that expression-relevant signal emerges from Ribo-seq training alone.

#### 1.20 Interpretability of sTASEP parameters across cell lines

The fitted sTASEP parameters provide a window into cell-line-specific translational kinetics. We release the full parameter sets for all four cell lines in Supplementary Table S14 and visualise them in Figures S20–S22.

The 61 codon-specific wait times ($\tau$) show markedly different variability across cell lines (Figure S20). HEK293 parameters are nearly uniform (range 0.484–0.528), consistent with the sparse ribosome profiling signal available for this dataset. In contrast, LCL exhibits the widest variation (range 0.251–1.170), followed by iPSC (0.290–0.817) and RPE-1 (0.272–0.752). Despite these differences in range, pairwise Spearman correlations reveal positive agreement among the three cell lines (Figure S21): iPSC–LCL $\rho = 0.421$ ($p < 0.001$), iPSC–RPE-1 $\rho = 0.383$ ($p = 0.002$), and LCL–RPE-1 $\rho = 0.268$ ($p = 0.037$). HEK293 shows no significant correlation with LCL or RPE-1 ($\rho < 0.05$, $p > 0.7$), consistent with its near-uniform parameter profile. The shared codon-level structure across iPSC, LCL, and RPE-1 suggests a conserved core of elongation kinetics that likely reflects the human tRNA pool [7, 8, 18]. Cell-line-specific deviations from this shared profile may reflect differences in tRNA expression, aminoacyl-tRNA charging, or tRNA modification status that alter the effective decoding rate of specific codons [6, 15, 21].

The base-pairing penalty ($\alpha$) is most pronounced in LCL, where codons with all three nucleotides engaged in base pairing incur a wait time of 0.702, approximately 50% higher than codons with no pairing (0.468) (Figure S22). This is consistent with structured mRNA regions acting as physical barriers that slow ribosome translocation [2, 20]. The angle parameters ($\beta$) in iPSC and RPE-1 reveal that codons within continuous helical stretches (no angle change) have the highest wait times (0.562 in iPSC), while positions at structural junctions (larger angle changes) allow faster elongation. This supports the view that unbroken double-stranded regions present the greatest resistance to the elongating ribosome, whereas junctions between structural elements are more easily traversed.

The positional bucket parameters ($\gamma$) show that in iPSC and RPE-1, the first third of the CDS carries the lowest additional wait time (0.431 and 0.440, respectively), while the middle and final thirds are slower (Figure S22).

To further validate the biological relevance of the fitted codon wait times, we computed the Codon Stabilization Coefficient (CSC) for each of the 61 sense codons in each cell line. The CSC of a codon $c$ is defined as the Pearson correlation, taken across all transcripts, between the per-transcript frequency of $c$ and the per-transcript ribosome density [19]. We then computed the Spearman correlation between the fitted sTASEP codon wait times $\vec{\tau}$ and the CSC values. All four cell lines show a positive correlation between fitted wait times and CSC: LCL $\rho = 0.418$ ($p = 7.9 \times 10^{-4}$), HEK293 $\rho = 0.354$ ($p = 5.2 \times 10^{-3}$), iPSC $\rho = 0.343$ ($p = 6.8 \times 10^{-3}$), and RPE-1 $\rho = 0.308$ ($p = 1.6 \times 10^{-2}$). This means that codons whose presence in a transcript is associated with higher ribosome density are assigned longer wait times by sTASEP, consistent with the interpretation that these codons slow elongation and cause local ribosome accumulation. This relationship emerges from the simulation-based fitting procedure without any prior linking wait times to CSC, providing independent evidence that the fitted parameters capture translational kinetics. The scatter plots in Figure S23 show this relationship for each cell line.

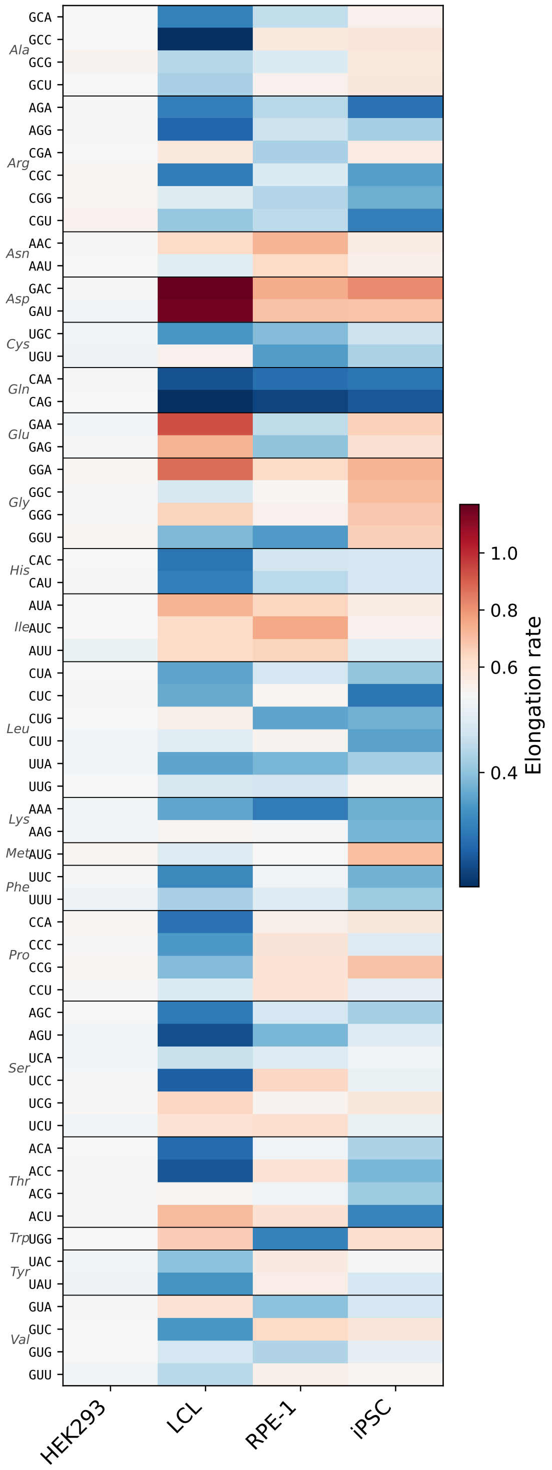


**Figure S20: Codon wait times across four cell lines.** Heatmap of estimated elongation-rate parameters for 61 sense codons in HEK293, LCL, RPE-1, and iPSC cells. Rows are grouped by encoded amino acid (italic labels, left margin). Color scale is centred at 0.5 (diverging red–blue); values above 0.5 indicate slower-than-average decoding. HEK293 rates cluster narrowly around 0.5. LCL and RPE-1 show the widest range, with GAC and GAU codons exhibiting the highest elongation rates in LCL.

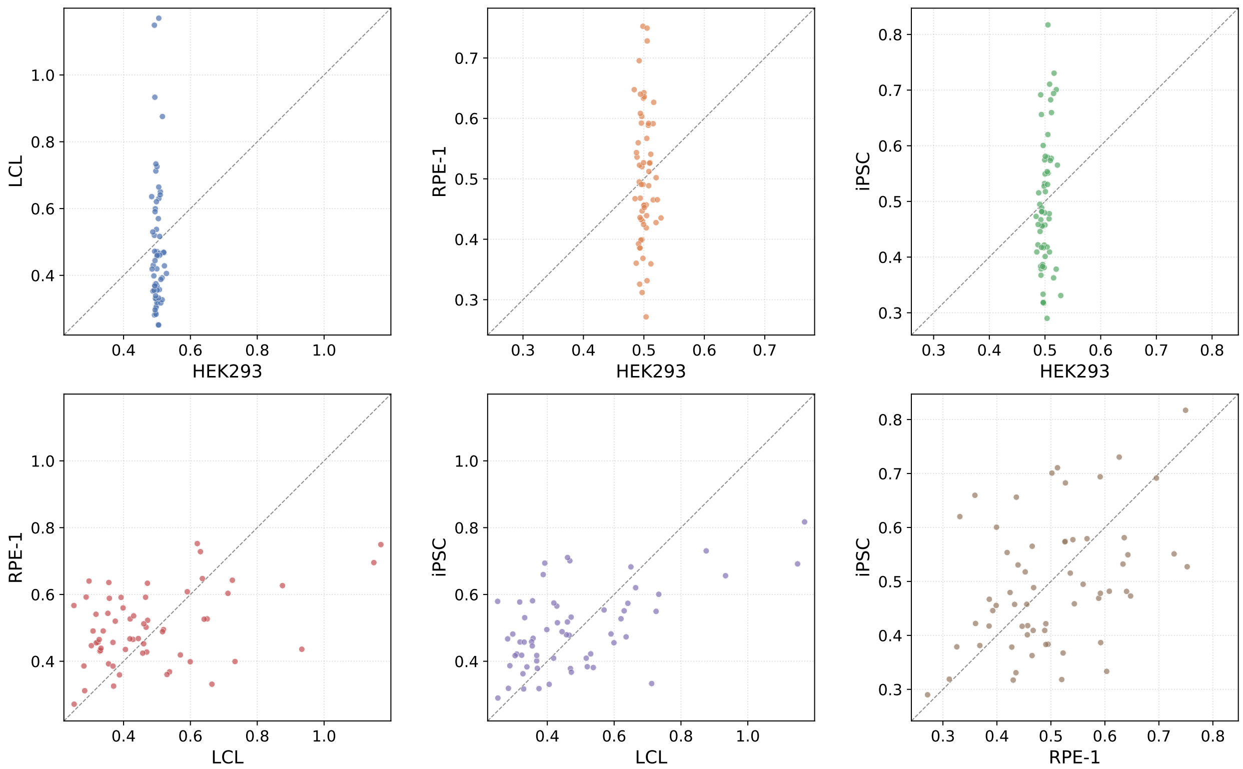


**Figure S21: Pairwise comparison of codon elongation rates.** Scatter plots of the 61 codon-rate parameters for all six cell-line pairs. Each point represents one sense codon; the dashed line indicates identity ($x = y$). Spearman rank-correlation coefficients ($\rho$) are annotated in each panel. Pairs involving HEK293 show compressed horizontal spread, reflecting its near-uniform rate profile. The three pairs (LCL–RPE-1, LCL–iPSC, RPE-1–iPSC) exhibit positive correlations, indicating that relative codon decoding speeds are broadly conserved across cell types despite differences in absolute magnitude.

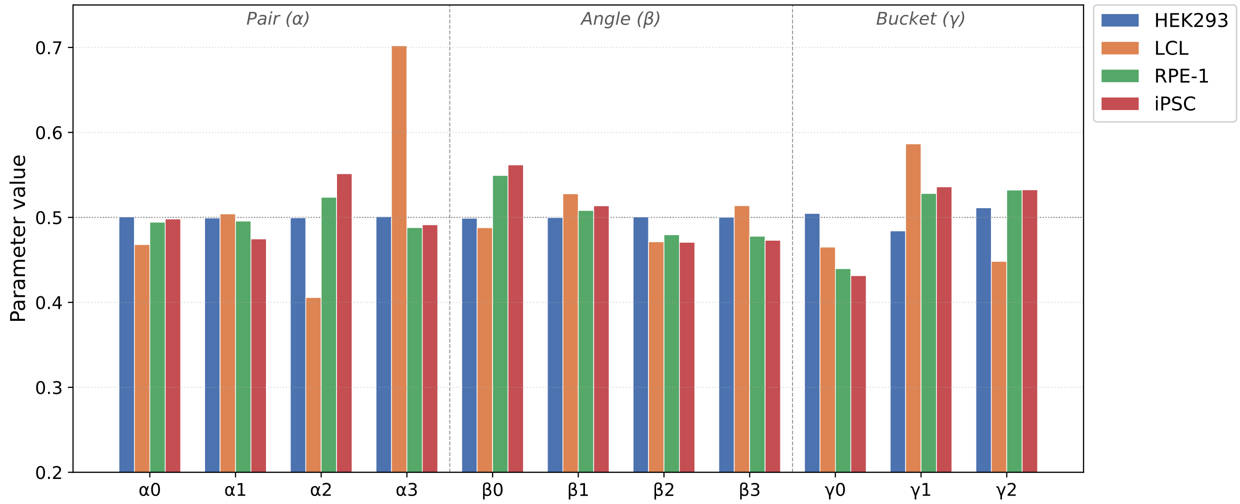


**Figure S22: Structural parameters across cell lines.** Grouped bar chart of the 11 non-codon model parameters: four base-pairing weights ($\alpha_0$–$\alpha_3$), four bond-angle weights ($\beta_0$–$\beta_3$), and three codon-bucket scaling factors ($\gamma_0$–$\gamma_2$). The dotted horizontal line marks the reference value of 0.5. HEK293 parameters are nearly flat at 0.5. LCL shows the most pronounced departures, particularly for $\alpha_3$ and $\gamma_1$, suggesting cell-type-specific differences in mRNA secondary-structure sensitivity and positional context effects on elongation.

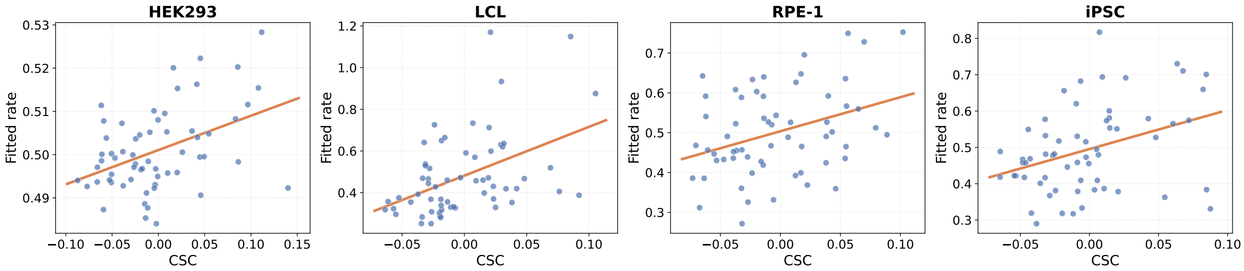


**Figure S23:** Fitted sTASEP codon wait times versus Codon Stabilization Coefficient (CSC) for each cell line. Each point represents one of the 61 sense codons. The orange line shows a linear fit. Positive correlations indicate that codons associated with higher ribosome density in the data receive longer fitted wait times.

#### 1.21 Cross-cell-line transfer

To test whether the polisher learns generalizable translational features, we evaluate each cell-line-specific `seq2ribo` model on the test sets of all four cell lines. Table S8 reports the full results.

The polisher retains substantial predictive power when transferred across cell lines. The iPSC model achieves a Tx-level $r$ of 0.576 on HEK293 and 0.524 on LCL, while the LCL model reaches 0.703 on iPSC and 0.540 on HEK293. These off-diagonal correlations indicate that a large fraction of the ribosome profile structure learned by each polisher generalizes beyond the training cell line, reflecting shared translational features such as codon-level elongation kinetics and mRNA structural barriers. Among the transferred models, the iPSC polisher shows the most consistent cross-cell performance.

We note that MAE metrics in the cross-cell setting should be interpreted with care. The HEK293 model frequently achieves the lowest pertranscriptMAE on non-HEK293 datasets, but this reflects conservative low-magnitude predictions rather than accurate profile reconstruction, as evidenced by its near-zero correlations on those datasets. This mirrors the pattern observed for the Translatomer baseline in the Results section.

While the matched models do achieve the highest correlations for their respective cell lines, the strong off-diagonal performance demonstrates that the polisher learns a largely transferable representation of ribosome dynamics.

**Table S8:** Cross-cell-line evaluation. Each column denotes the cell line on which the polisher was trained; each row block denotes the cell line on which the model is evaluated. Diagonal entries (same-cell) are underlined. Bold indicates the best model for each data cell line and metric.

| Eval. cell line | Metric | iPSC model | HEK293 model | LCL model | RPE-1 model |
|---|---|---|---|---|---|
| iPSC | pertranscriptMAE | 9.75 | **1.15** | 138.37 | 21.23 |
| iPSC | elemwiseMAE | 4.02 | **3.00** | 18.27 | 4.30 |
| iPSC | Shape $r$ | **0.162** | 0.021 | 0.029 | 0.041 |
| iPSC | Tx-level $r$ | **0.920** | 0.295 | 0.703 | 0.339 |
| iPSC | Elemwise $r$ | **0.598** | 0.162 | 0.205 | 0.145 |
| iPSC | codonMAE | 41,422 | 36,511 | 130,017 | **10,878** |
| iPSC | pairMAE | 662,755 | 584,203 | 2,080,311 | **171,261** |
| iPSC | angleMAE | 662,755 | 584,203 | 2,080,311 | **171,261** |
| iPSC | bucketMAE | 883,673 | 778,937 | 2,773,749 | **228,348** |
| HEK293 | pertranscriptMAE | 112.03 | **1.31** | 528.80 | 216.89 |
| HEK293 | elemwiseMAE | 3.50 | **0.14** | 17.27 | 2.98 |
| HEK293 | Shape $r$ | 0.031 | **0.054** | 0.010 | 0.022 |
| HEK293 | Tx-level $r$ | 0.576 | **0.657** | 0.540 | 0.223 |
| HEK293 | Elemwise $r$ | 0.197 | **0.396** | 0.082 | 0.123 |
| HEK293 | codonMAE | 21,826 | **3,759** | 123,147 | 22,442 |
| HEK293 | pairMAE | 349,230 | **60,146** | 1,970,386 | 359,105 |
| HEK293 | angleMAE | 349,230 | **60,146** | 1,970,386 | 359,105 |
| HEK293 | bucketMAE | 465,640 | **80,194** | 2,627,181 | 478,807 |
| LCL | pertranscriptMAE | 78.54 | **1.47** | 83.00 | 95.40 |
| LCL | elemwiseMAE | 17.33 | **15.63** | 21.72 | 16.46 |
| LCL | Shape $r$ | 0.020 | 0.005 | **0.186** | 0.009 |
| LCL | Tx-level $r$ | 0.524 | 0.272 | **0.901** | 0.246 |
| LCL | Elemwise $r$ | 0.131 | 0.047 | **0.612** | 0.087 |
| LCL | codonMAE | **132,809** | 214,500 | 155,512 | 155,682 |
| LCL | pairMAE | **2,124,929** | 3,432,009 | 2,487,817 | 2,490,898 |
| LCL | angleMAE | **2,124,929** | 3,432,009 | 2,487,817 | 2,490,898 |
| LCL | bucketMAE | **2,833,238** | 4,576,012 | 3,317,090 | 3,321,198 |
| RPE-1 | pertranscriptMAE | 106.56 | **1.65** | 334.27 | 17.14 |
| RPE-1 | elemwiseMAE | 4.85 | **2.79** | 18.11 | 2.92 |
| RPE-1 | Shape $r$ | 0.037 | 0.008 | 0.008 | **0.139** |
| RPE-1 | Tx-level $r$ | 0.428 | 0.097 | 0.165 | **0.771** |
| RPE-1 | Elemwise $r$ | 0.190 | 0.097 | 0.059 | **0.610** |
| RPE-1 | codonMAE | **9,353** | 29,011 | 104,126 | 35,652 |
| RPE-1 | pairMAE | **149,552** | 464,195 | 1,666,033 | 570,437 |
| RPE-1 | angleMAE | **149,552** | 464,195 | 1,666,033 | 570,437 |
| RPE-1 | bucketMAE | **199,402** | 618,927 | 2,221,377 | 760,582 |

##### 1.22 Bootstrap Confidence Intervals

To assess the statistical reliability of `seq2ribo`'s performance, we computed bootstrap 95% confidence intervals (CIs) for all evaluation metrics by resampling test-set transcripts with replacement (1,000 iterations). For each cell line, we resampled the test transcripts, recomputing all metrics on each resample. Predictions were averaged across $K = 32$ independent sTASEP simulation runs per transcript prior to metric computation.

Table S9 reports bootstrap CIs for all evaluation metrics. Correlation metrics are computed on unscaled (raw count) predictions, while MAE metrics are computed on scaled predictions normalized to match observed total counts per transcript, thereby isolating profile-shape accuracy.

For codon-level shape, `seq2ribo` achieves Pearson $r = 0.054$–$0.186$ across cell lines, with CIs that exclude zero in every case (e.g. HEK293: 0.054 $[0.049, 0.060]$; LCL: 0.186 $[0.178, 0.195]$), confirming statistically significant positional signal. At the transcript level, the model achieves Pearson $r = 0.657$–$0.920$, and element-wise Pearson correlations range from 0.396 to 0.612, with CIs well above zero even for HEK293 ($[0.314, 0.492]$), the most sparse cell line.

Scaled per-transcript MAE is tightly bounded across bootstrap resamples; for example, the HEK293 estimate is 1.909 $[1.901, 1.916]$ and the iPSC estimate is 1.386 $[1.373, 1.399]$. The binned MAEs exhibit wider CIs, particularly for pair and angle MAE in LCL and RPE-1, reflecting the sensitivity of these aggregate statistics to individual high-count transcripts.

**Table S9:** `seq2ribo` test set metrics with bootstrap 95% confidence intervals (1,000 resamples). Correlation metrics are computed on unscaled (raw count) predictions; MAE metrics are computed on scaled predictions normalized to match observed total counts per transcript. Values are reported as point estimate [95% CI].

| Cell line | Metric | `seq2ribo` |
|---|---|---|
| iPSC | pertranscriptMAE | 1.386 [1.373, 1.399] |
| iPSC | elemwiseMAE | 4.127 [3.891, 4.393] |
| iPSC | Shape *r* | 0.162 [0.155, 0.169] |
| iPSC | Tx-level *r* | 0.920 [0.883, 0.946] |
| iPSC | Elemwise *r* | 0.598 [0.559, 0.644] |
| iPSC | codonMAE | 12,028 [11,416, 13,995] |
| iPSC | pairMAE | 44,153 [30,562, 59,715] |
| iPSC | angleMAE | 85,032 [68,102, 101,054] |
| iPSC | bucketMAE | 287,231 [227,716, 353,525] |
| HEK293 | pertranscriptMAE | 1.909 [1.901, 1.916] |
| HEK293 | elemwiseMAE | 0.247 [0.232, 0.265] |
| HEK293 | Shape *r* | 0.054 [0.049, 0.060] |
| HEK293 | Tx-level *r* | 0.657 [0.578, 0.741] |
| HEK293 | Elemwise *r* | 0.396 [0.314, 0.492] |
| HEK293 | codonMAE | 3,627 [3,381, 3,933] |
| HEK293 | pairMAE | 15,164 [12,493, 18,118] |
| HEK293 | angleMAE | 23,899 [21,112, 26,790] |
| HEK293 | bucketMAE | 66,934 [62,404, 71,809] |
| LCL | pertranscriptMAE | 1.471 [1.459, 1.482] |
| LCL | elemwiseMAE | 22.41 [20.34, 24.71] |
| LCL | Shape *r* | 0.186 [0.178, 0.195] |
| LCL | Tx-level *r* | 0.901 [0.773, 0.959] |
| LCL | Elemwise *r* | 0.612 [0.442, 0.717] |
| LCL | codonMAE | 38,902 [37,303, 67,690] |
| LCL | pairMAE | 187,622 [70,073, 411,657] |
| LCL | angleMAE | 84,351 [36,425, 301,782] |
| LCL | bucketMAE | 920,437 [528,781, 1,355,892] |
| RPE-1 | pertranscriptMAE | 1.503 [1.489, 1.517] |
| RPE-1 | elemwiseMAE | 3.359 [2.906, 4.015] |
| RPE-1 | Shape *r* | 0.139 [0.131, 0.146] |
| RPE-1 | Tx-level *r* | 0.771 [0.480, 0.980] |
| RPE-1 | Elemwise *r* | 0.610 [0.482, 0.724] |
| RPE-1 | codonMAE | 12,404 [7,348, 24,553] |
| RPE-1 | pairMAE | 24,184 [11,071, 70,882] |
| RPE-1 | angleMAE | 15,336 [10,098, 48,627] |
| RPE-1 | bucketMAE | 356,474 [224,699, 576,959] |

##### 1.23 Variance Across Simulation Runs

Each transcript is associated with $K = 32$ independent sTASEP simulation runs. To quantify how much individual simulation stochasticity affects downstream metrics, we evaluate (i) the variance of each metric when computed on a single run, and (ii) how that variance decreases as predictions are averaged over increasing numbers of runs.

###### 1.23.1 Individual run variance

Table S10 reports the mean and standard deviation of all evaluation metrics computed independently for each of the 32 simulation runs. Correlation metrics are computed on unscaled predictions; MAE metrics are computed on scaled predictions normalized to match observed total counts per transcript.

For the model, all correlation standard deviations are at least two orders of magnitude smaller than the corresponding means (e.g. Shape $r$: std $< 3 \times 10^{-4}$ across all cell lines; Tx-level $r$: std $< 5 \times 10^{-4}$), indicating that any single simulation run already yields a highly stable metric estimate. The sTASEP simulation baseline exhibits similarly low absolute variability in its correlation values.

The scaled pertranscriptMAE shows negligible run-to-run variability for both model and baseline (std $< 5 \times 10^{-4}$). The binned MAE metrics (particularly pairMAE and angleMAE) show somewhat larger absolute variability, which is expected because these aggregate statistics are sensitive to the precise codon-level count distribution within each stochastic simulation.

###### 1.23.2 Convergence with number of averaged runs

Table S11 shows how the standard deviation of selected metrics decreases as predictions are averaged over $K = 1, 2, 4, 8, 16$, and 32 runs (50 random subsets per size). For the model, the standard deviation of Shape $r$ decreases by roughly $\sqrt{K}$, consistent with independent sampling, and is already below $10^{-4}$ at $K = 4$ in all cell lines. Tx-level $\rho$ converges more slowly but stabilizes by $K =$ 8–16. At $K = 32$ (all runs averaged), the standard deviation is effectively zero, confirming full convergence. These results justify our use of $K = 32$ averaged runs for all reported metrics, while also demonstrating that even substantially fewer runs would yield reliable estimates.

**Table S10:** Variance across individual simulation runs ($K = 32$). Each metric is computed independently per run; reported values are mean ± std across runs. Correlation metrics are computed on unscaled predictions; MAE metrics are computed on scaled predictions.

| Cell line | Metric | sTASEP | `seq2ribo` |
|---|---|---|---|
| iPSC | Tx-level $r$ | 0.209 ± 0.001 | 0.920 ± 4e-4 |
| | Shape $r$ | 0.019 ± 0.001 | 0.161 ± 3e-4 |
| | Elemwise $r$ | 0.072 ± 0.001 | 0.597 ± 0.001 |
| | Shape $\rho$ | 0.021 ± 0.001 | 0.161 ± 2e-4 |
| | Tx-level $\rho$ | 0.445 ± 0.002 | 0.731 ± 0.001 |
| | Elemwise $\rho$ | 0.213 ± 0.001 | 0.478 ± 5e-4 |
| | pertranscriptMAE | 1.308 ± 3e-4 | 1.386 ± 4e-4 |
| | elemwiseMAE | 4.566 ± 0.003 | 4.135 ± 0.002 |
| | codonMAE | 3,642 ± 108 | 12,035 ± 116 |
| | pairMAE | 13,237 ± 1,365 | 44,153 ± 1,740 |
| | angleMAE | 10,443 ± 1,578 | 85,032 ± 1,798 |
| | bucketMAE | 27,502 ± 6,340 | 287,231 ± 2,761 |

| Cell line | Metric | sTASEP | `seq2ribo` |
|---|---|---|---|
| HEK293 | Tx-level $r$ | 0.012 ± 0.003 | 0.657 ± 2e-4 |
| | Shape $r$ | 0.002 ± 0.001 | 0.054 ± 4e-5 |
| | Elemwise $r$ | 0.000 ± 0.002 | 0.396 ± 2e-4 |
| | Shape $\rho$ | 0.002 ± 0.001 | 0.029 ± 1e-4 |
| | Tx-level $\rho$ | 0.100 ± 0.004 | 0.292 ± 0.001 |
| | Elemwise $\rho$ | 0.000 ± 0.002 | 0.177 ± 7e-5 |
| | pertranscriptMAE | 1.807 ± 3e-4 | 1.909 ± 5e-5 |
| | elemwiseMAE | 0.221 ± 2e-4 | 0.247 ± 1e-5 |
| | codonMAE | 183 ± 4 | 3,627 ± 3 |
| | pairMAE | 375 ± 101 | 15,164 ± 32 |
| | angleMAE | 290 ± 57 | 23,899 ± 51 |
| | bucketMAE | 6,545 ± 227 | 66,934 ± 45 |

| Cell line | Metric | sTASEP | `seq2ribo` |
|---|---|---|---|
| LCL | Tx-level $r$ | 0.098 ± 0.003 | 0.901 ± 3e-4 |
| | Shape $r$ | 0.019 ± 0.001 | 0.186 ± 1e-4 |
| | Elemwise $r$ | 0.006 ± 0.001 | 0.612 ± 5e-4 |
| | Shape $\rho$ | 0.023 ± 0.001 | 0.159 ± 6e-5 |
| | Tx-level $\rho$ | 0.171 ± 0.002 | 0.767 ± 2e-4 |
| | Elemwise $\rho$ | 0.018 ± 0.002 | 0.503 ± 2e-4 |
| | pertranscriptMAE | 1.519 ± 3e-4 | 1.471 ± 1e-4 |
| | elemwiseMAE | 28.91 ± 0.03 | 22.42 ± 0.004 |
| | codonMAE | 36,304 ± 1,177 | 38,912 ± 154 |
| | pairMAE | 251,541 ± 17,900 | 187,622 ± 1,867 |
| | angleMAE | 164,994 ± 16,315 | 84,351 ± 2,525 |
| | bucketMAE | 330,954 ± 42,686 | 920,437 ± 5,442 |

| Cell line | Metric | sTASEP | `seq2ribo` |
|---|---|---|---|
| RPE-1 | Tx-level $r$ | 0.127 ± 0.007 | 0.771 ± 1e-4 |
| | Shape $r$ | 0.013 ± 0.001 | 0.138 ± 8e-5 |
| | Elemwise $r$ | 0.005 ± 0.003 | 0.610 ± 4e-4 |
| | Shape $\rho$ | 0.016 ± 0.001 | 0.125 ± 8e-5 |
| | Tx-level $\rho$ | 0.236 ± 0.003 | 0.638 ± 5e-4 |
| | Elemwise $\rho$ | 0.010 ± 0.004 | 0.436 ± 3e-4 |
| | pertranscriptMAE | 1.370 ± 4e-4 | 1.503 ± 2e-4 |
| | elemwiseMAE | 3.508 ± 0.005 | 3.362 ± 0.001 |
| | codonMAE | 2,481 ± 96 | 12,405 ± 78 |
| | pairMAE | 7,054 ± 2,039 | 24,267 ± 1,579 |
| | angleMAE | 4,572 ± 2,001 | 15,508 ± 1,309 |
| | bucketMAE | 29,463 ± 10,554 | 356,474 ± 2,384 |

**Table S11:** Convergence of metric standard deviation with number of averaged simulation runs ($K$). Values are the standard deviation of each metric across 50 random subsets of size $K$. Only `seq2ribo` results are shown.

| **Cell line** | **Metric** | $K$=1 | $K$=2 | $K$=4 | $K$=8 | $K$=16 | $K$=32 |
|---|---|---|---|---|---|---|---|
| HEK293 | Shape $r$ | 3.6e-5 | 3.1e-5 | 1.7e-5 | 1.2e-5 | 7.3e-6 | 0 |
| | Tx-level $\rho$ | 1.3e-3 | 9.6e-4 | 8.7e-4 | 8.9e-4 | 6.6e-4 | 0 |
| | pertranscriptMAE | 5.3e-5 | 3.1e-5 | 2.3e-5 | 1.5e-5 | 9.4e-6 | 0 |
| iPSC | Shape $r$ | 2.5e-4 | 1.8e-4 | 1.1e-4 | 8.1e-5 | 5.1e-5 | 0 |
| | Tx-level $\rho$ | 1.4e-3 | 9.0e-4 | 5.8e-4 | 3.5e-4 | 2.1e-4 | 0 |
| | pertranscriptMAE | 3.7e-4 | 2.7e-4 | 1.9e-4 | 1.4e-4 | 8.1e-5 | 0 |
| LCL | Shape $r$ | 8.6e-5 | 6.9e-5 | 4.2e-5 | 3.0e-5 | 1.9e-5 | 0 |
| | Tx-level $\rho$ | 2.6e-4 | 1.9e-4 | 1.2e-4 | 7.3e-5 | 4.7e-5 | 0 |
| | pertranscriptMAE | 9.3e-5 | 7.6e-5 | 4.4e-5 | 3.4e-5 | 2.1e-5 | 0 |
| RPE-1 | Shape $r$ | 6.8e-5 | 5.7e-5 | 3.5e-5 | 2.1e-5 | 1.6e-5 | 0 |
| | Tx-level $\rho$ | 4.4e-4 | 3.4e-4 | 1.9e-4 | 1.2e-4 | 9.2e-5 | 0 |
| | pertranscriptMAE | 1.6e-4 | 1.0e-4 | 8.3e-5 | 4.9e-5 | 3.0e-5 | 0 |

#### 1.24 Hardware and Runtime

##### 1.24.1 Hardware and Software Environment

All computational experiments were conducted on a high-performance computing node running Ubuntu 22.04.5 LTS. The CPU environment consisted of dual Intel(R) Xeon(R) Gold 6430 processors, providing a total of 64 physical cores (128 logical cores) and approximately 500 GB of system RAM. For deep learning tasks, we use a GPU environment containing NVIDIA RTX A6000 (48 GB VRAM) and NVIDIA A100-SXM4 (40 GB VRAM). The `seq2ribo` framework was implemented in Python using the PyTorch library. All dependencies and library versions are documented in the GitHub repository.

##### 1.24.2 Theoretical Complexity

The sTASEP module uses discrete-event Monte Carlo simulations. The computational complexity scales linearly, $O(T \times R)$, where $T$ is the number of simulation steps required to reach steady state and $R$ is the number of ribosomes. This event-driven approach avoids the computational overhead of solving large systems of differential equations and allows for efficient parallelization across CPU cores.

A primary advantage of selecting the Mamba backbone over standard Transformer architectures is its linear scaling with respect to sequence length. While the self-attention mechanism in Transformers scales quadratically $O(N^2)$ with input sequence length $N$, the Structured State Space Model (SSM) backbone of Mamba scales linearly $O(N)$.

##### 1.24.3 Practical Runtime Estimates and Implementation Details

We benchmarked the runtime of the `seq2ribo` pipeline across our training and inference workflows.

The fitting of kinetic parameters (wait times for codons and structural features) for the complete training set was parallelized across all 128 logical CPU cores and required approximately one week to reach convergence. Training the Mamba-based polisher was performed on a single GPU (using either an RTX A6000 or A100). Training a separate polisher model for each cell line took approximately 3 days. To maximize hardware efficiency, we used batch sizes ranging from 192 to 256, depending on the cell line, ensuring that at least 80% of GPU memory was used during training. Fine-tuning the task-specific heads for translation efficiency (TE) and protein expression was significantly faster, requiring a batch size of 192 and completing in approximately 30 minutes.

For inference on a single mRNA sequence, the sTASEP simulation typically completes in seconds to a few minutes, depending on transcript length and the number of trajectories. The polisher inference and downstream prediction heads process sequences in seconds.

### 2. Supplementary Tables

#### 2.1 Unscaled losses

**Table S12:** Test set metrics (raw) across cell lines. Lower is better for MAE metrics, while higher is better for Pearson correlation ($r$). Bold indicates the best performance within each cell line and metric.

| Cell line | Metric | TASEP | sTASEP | Translatomer (seq.) | seq2ribo |
|---|---|---|---|---|---|
| iPSC | pertranscriptMAE | 28.70 | 57.45 | **5.42** | 9.75 |
| iPSC | elemwiseMAE | 8.11 | 12.47 | **1.66** | 4.02 |
| iPSC | Shape $r$ | −0.015 | 0.039 | 0.001 | **0.162** |
| iPSC | Tx-level $r$ | 0.200 | 0.210 | 0.051 | **0.920** |
| iPSC | Elemwise $r$ | 0.056 | 0.078 | −0.002 | **0.598** |
| iPSC | codonMAE | 81591.4 | 296740.5 | 44969.0 | **41422.1** |
| iPSC | pairMAE | 1273106.8 | 4747805.5 | 2667391.3 | **662754.9** |
| iPSC | angleMAE | 1273106.8 | 4747805.5 | 2667391.3 | **662754.9** |
| iPSC | bucketMAE | 1697475.7 | 6330407.4 | 3556521.8 | **883673.2** |
| HEK293 | pertranscriptMAE | 131.62 | 305.52 | **1.10** | 1.31 |
| HEK293 | elemwiseMAE | 1.95 | 4.54 | **0.05** | 0.14 |
| HEK293 | Shape $r$ | 0.002 | 0.007 | 0.000 | **0.054** |
| HEK293 | Tx-level $r$ | 0.006 | 0.013 | 0.023 | **0.657** |
| HEK293 | Elemwise $r$ | −0.004 | −0.001 | 0.000 | **0.396** |
| HEK293 | codonMAE | 57855.5 | 143093.6 | **1631.6** | 3759.2 |
| HEK293 | pairMAE | 925688.3 | 2289497.3 | 75711.7 | **60145.8** |
| HEK293 | angleMAE | 925688.3 | 2289497.3 | 75711.7 | **60145.8** |
| HEK293 | bucketMAE | 1234251.0 | 3052663.1 | 100948.9 | **80194.4** |
| LCL | pertranscriptMAE | 148.64 | 156.90 | **69.21** | 83.00 |
| LCL | elemwiseMAE | 24.21 | 23.81 | **7.76** | 21.72 |
| LCL | Shape $r$ | −0.012 | 0.046 | 0.000 | **0.186** |
| LCL | Tx-level $r$ | 0.079 | 0.101 | 0.055 | **0.901** |
| LCL | Elemwise $r$ | −0.006 | 0.014 | 0.000 | **0.612** |
| LCL | codonMAE | 742530.8 | 677243.8 | 208501.0 | **155512.2** |
| LCL | pairMAE | 11027928.5 | 10835930.4 | 12558537.6 | **2487817.4** |
| LCL | angleMAE | 11027928.5 | 10835930.4 | 12558537.6 | **2487817.4** |
| LCL | bucketMAE | 14703904.7 | 14447907.2 | 16744716.9 | **3317089.9** |
| RPE-1 | pertranscriptMAE | 83.54 | 184.78 | 34.32 | **17.14** |
| RPE-1 | elemwiseMAE | 4.29 | 5.95 | **1.33** | 2.92 |
| RPE-1 | Shape $r$ | −0.005 | 0.041 | 0.003 | **0.139** |
| RPE-1 | Tx-level $r$ | 0.137 | 0.130 | 0.035 | **0.771** |
| RPE-1 | Elemwise $r$ | 0.003 | 0.017 | 0.003 | **0.610** |
| RPE-1 | codonMAE | 45379.9 | 41746.9 | **11298.9** | 35652.3 |
| RPE-1 | pairMAE | 655463.8 | 667931.4 | 1376701.0 | **570436.8** |
| RPE-1 | angleMAE | 655463.8 | 667931.4 | 1376701.0 | **570436.8** |
| RPE-1 | bucketMAE | 873951.7 | 890575.2 | 1835601.3 | **760582.4** |

**Table S13:** Summary of polisher feature ablation results (raw) on the iPSC test set across six error metrics and two correlation metrics. Lower is better for all MAE metrics, while higher is better for Pearson and Spearman. Bold indicates the best performance in each row.

| | **Seq.** | **sTASEP** | **Struc.** | **Seq+sTASEP** | **Seq+Struc.** | **sTASEP+Struc.** | `seq2ribo` |
|---|---|---|---|---|---|---|---|
| pertranscriptMAE | 18.77 | 36.53 | 36.76 | 16.59 | 17.24 | 26.58 | **9.75** |
| elemwiseMAE | 4.66 | 6.60 | 6.05 | 4.67 | 4.82 | 5.96 | **4.02** |
| Shape *r* | 0.1293 | 0.0575 | 0.0259 | 0.1199 | 0.1016 | 0.0512 | **0.1620** |
| Tx-level *r* | 0.8983 | 0.2158 | 0.2051 | 0.9133 | 0.9032 | 0.2583 | **0.9204** |
| Elemwise *r* | 0.5536 | 0.1020 | 0.0839 | 0.5626 | 0.5456 | 0.1280 | **0.5981** |
| codonMAE | 7,958 | 22,025 | 41,483 | **4,545** | 5,225 | 34,231 | 41,422 |
| pairMAE | 110,400 | 123,510 | 654,086 | 23,530 | **19,624** | 533,415 | 662,755 |
| angleMAE | 88,320 | 98,808 | 654,086 | **22,146** | 22,465 | 533,415 | 662,755 |
| bucketMAE | 147,200 | 612,756 | 872,115 | **58,205** | 67,274 | 711,220 | 883,673 |

#### 2.2 Fitted parameters

**Table S14:** Estimated model parameters across four cell lines.

| Parameter | HEK293 | LCL | RPE-1 | iPSC |
|---|---|---|---|---|
| AAA | 0.4930 | 0.3701 | 0.3257 | 0.3788 |
| AAC | 0.5055 | 0.6300 | 0.7282 | 0.5510 |
| AAG | 0.4923 | 0.5196 | 0.4949 | 0.3836 |
| AAU | 0.4992 | 0.4714 | 0.6335 | 0.5322 |
| ACA | 0.4986 | 0.3086 | 0.4906 | 0.4217 |
| ACC | 0.4959 | 0.2879 | 0.5922 | 0.3867 |
| ACG | 0.5080 | 0.5165 | 0.4886 | 0.4092 |
| ACU | 0.4965 | 0.7123 | 0.6033 | 0.3333 |
| AGA | 0.4978 | 0.3292 | 0.4302 | 0.3172 |
| AGC | 0.5038 | 0.3233 | 0.4569 | 0.4181 |
| AGG | 0.4971 | 0.3036 | 0.4468 | 0.4169 |
| AGU | 0.4926 | 0.2813 | 0.3859 | 0.4670 |
| AUA | 0.5001 | 0.7256 | 0.6426 | 0.5494 |
| AUC | 0.4983 | 0.6205 | 0.7524 | 0.5273 |
| AUG | 0.5203 | 0.4674 | 0.5019 | 0.7008 |
| AUU | 0.4840 | 0.6358 | 0.6476 | 0.4730 |
| CAA | 0.4971 | 0.2836 | 0.3118 | 0.3190 |
| CAC | 0.4999 | 0.3184 | 0.4553 | 0.4577 |
| CAG | 0.5036 | 0.2520 | 0.2715 | 0.2900 |
| CAU | 0.4954 | 0.3308 | 0.4328 | 0.4574 |
| CCA | 0.5114 | 0.3174 | 0.5407 | 0.5774 |
| CCC | 0.5072 | 0.3575 | 0.5885 | 0.4688 |
| CCG | 0.5153 | 0.3925 | 0.5913 | 0.6939 |
| CCU | 0.5078 | 0.4658 | 0.5917 | 0.4780 |
| CGA | 0.5040 | 0.5699 | 0.4189 | 0.5536 |
| CGC | 0.5154 | 0.3271 | 0.4651 | 0.3626 |
| CGG | 0.5201 | 0.4692 | 0.4276 | 0.3783 |
| CGU | 0.5283 | 0.4057 | 0.4353 | 0.3309 |
| CUA | 0.5002 | 0.3682 | 0.4564 | 0.4011 |
| CUC | 0.4968 | 0.3751 | 0.5201 | 0.3182 |
| CUG | 0.4984 | 0.5376 | 0.3684 | 0.3812 |
| CUU | 0.4928 | 0.4720 | 0.5226 | 0.3674 |
| GAA | 0.4936 | 0.9333 | 0.4360 | 0.6561 |

*Continued on next page*

| Parameter | HEK293 | LCL | RPE-1 | iPSC |
|---|---|---|---|---|
| GAC | 0.5053 | 1.1699 | 0.7494 | 0.8172 |
| GAG | 0.4967 | 0.7335 | 0.3993 | 0.6005 |
| GAU | 0.4922 | 1.1489 | 0.6954 | 0.6915 |
| GCA | 0.5046 | 0.3323 | 0.4391 | 0.5305 |
| GCC | 0.5048 | 0.2514 | 0.5669 | 0.5793 |
| GCG | 0.5223 | 0.4284 | 0.4654 | 0.5652 |
| GCU | 0.4994 | 0.4195 | 0.5267 | 0.5745 |
| GGA | 0.5163 | 0.8756 | 0.6265 | 0.7304 |
| GGC | 0.5083 | 0.4605 | 0.5122 | 0.7106 |
| GGG | 0.5101 | 0.6501 | 0.5269 | 0.6825 |
| GGU | 0.5116 | 0.3877 | 0.3592 | 0.6596 |
| GUA | 0.4950 | 0.5994 | 0.3986 | 0.4555 |
| GUC | 0.5006 | 0.3559 | 0.6358 | 0.5811 |
| GUG | 0.4995 | 0.4572 | 0.4244 | 0.4795 |
| GUU | 0.4886 | 0.4302 | 0.5359 | 0.5154 |
| UAC | 0.4906 | 0.3985 | 0.5597 | 0.4948 |
| UAU | 0.4877 | 0.3530 | 0.5434 | 0.4586 |
| UCA | 0.4940 | 0.4442 | 0.4680 | 0.4885 |
| UCC | 0.4942 | 0.2966 | 0.6401 | 0.4815 |
| UCG | 0.5095 | 0.6411 | 0.5259 | 0.5735 |
| UCU | 0.4941 | 0.5906 | 0.6082 | 0.4818 |
| UGC | 0.4911 | 0.3550 | 0.3924 | 0.4460 |
| UGG | 0.5052 | 0.6644 | 0.3314 | 0.6202 |
| UGU | 0.4873 | 0.5298 | 0.3604 | 0.4221 |
| UUA | 0.4937 | 0.3683 | 0.3855 | 0.4174 |
| UUC | 0.4957 | 0.3390 | 0.4907 | 0.3832 |
| UUG | 0.5007 | 0.4601 | 0.4525 | 0.5176 |
| UUU | 0.4853 | 0.4189 | 0.4671 | 0.4092 |
| $\alpha_0$ | 0.5007 | 0.4680 | 0.4944 | 0.4981 |
| $\alpha_1$ | 0.4993 | 0.5041 | 0.4956 | 0.4746 |
| $\alpha_2$ | 0.4996 | 0.4056 | 0.5237 | 0.5514 |
| $\alpha_3$ | 0.5009 | 0.7020 | 0.4880 | 0.4913 |
| $\beta_0$ | 0.4990 | 0.4879 | 0.5494 | 0.5618 |
| $\beta_1$ | 0.4998 | 0.5278 | 0.5081 | 0.5136 |
| $\beta_2$ | 0.5006 | 0.4713 | 0.4796 | 0.4707 |
| $\beta_3$ | 0.5003 | 0.5138 | 0.4778 | 0.4731 |

*Continued on next page*

| Parameter | HEK293 | LCL | RPE-1 | iPSC |
|---|---|---|---|---|
| $\gamma_0$ | 0.5046 | 0.4650 | 0.4397 | 0.4314 |
| $\gamma_1$ | 0.4841 | 0.5865 | 0.5282 | 0.5359 |
| $\gamma_2$ | 0.5113 | 0.4483 | 0.5322 | 0.5324 |

#### 2.3 Translation efficiency

**Table S15:** Translation efficiency prediction using raw sTASEP and polisher outputs on the test set. We report Pearson correlation $r$ between predicted total ribosome load and experimental TE.

| Cell line | sTASEP | Polisher |
|---|---|---|
| Mean | −0.239 | −0.451 |
| HEK293 | −0.360 | −0.360 |
| LCL | −0.506 | 0.250 |
| RPE-1 | −0.397 | 0.190 |

#### 2.4 Protein expression

**Table S16:** Protein expression prediction using raw sTASEP and polisher outputs on the test set. We report Pearson correlation $r$ between predicted total ribosome load and experimental protein expression.

| Cell line | sTASEP | Polisher |
|---|---|---|
| iPSC | −0.061 | 0.030 |
| HEK293 | −0.095 | 0.024 |
| LCL | −0.162 | 0.232 |
| RPE-1 | −0.247 | 0.123 |

### 3. Supplementary Figures

#### 3.1 Training Loss Curves for Simulation Methods Across Cell Lines

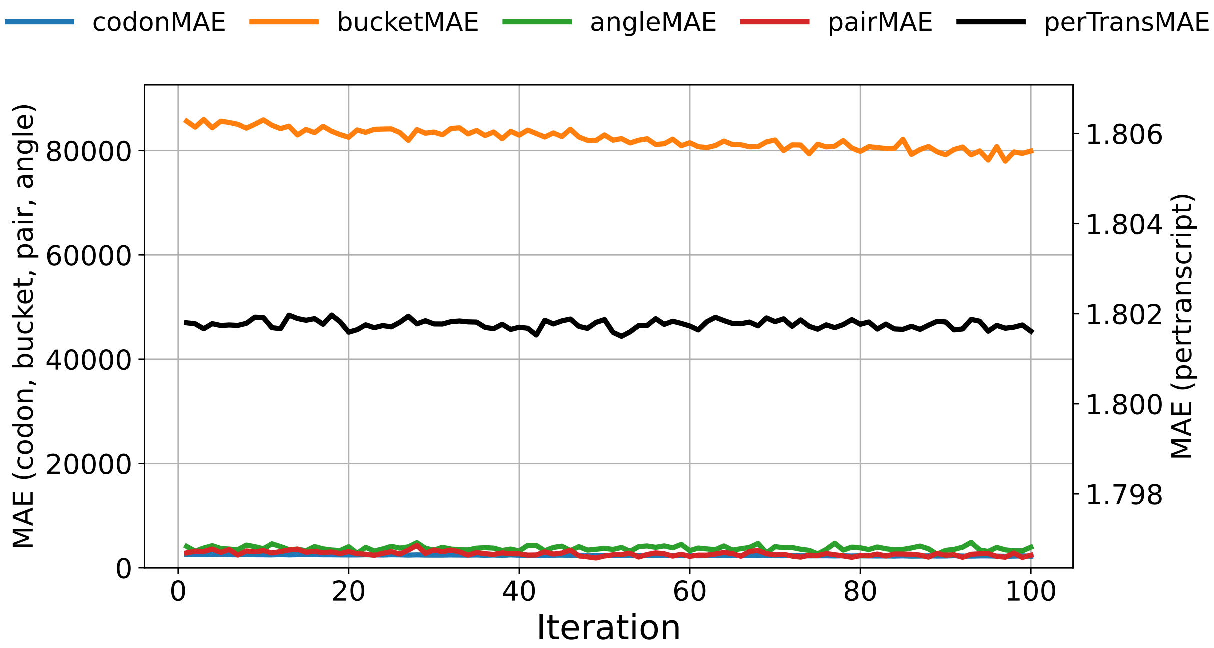


**Figure S24:** sTASEP training loss convergence for HEK293.

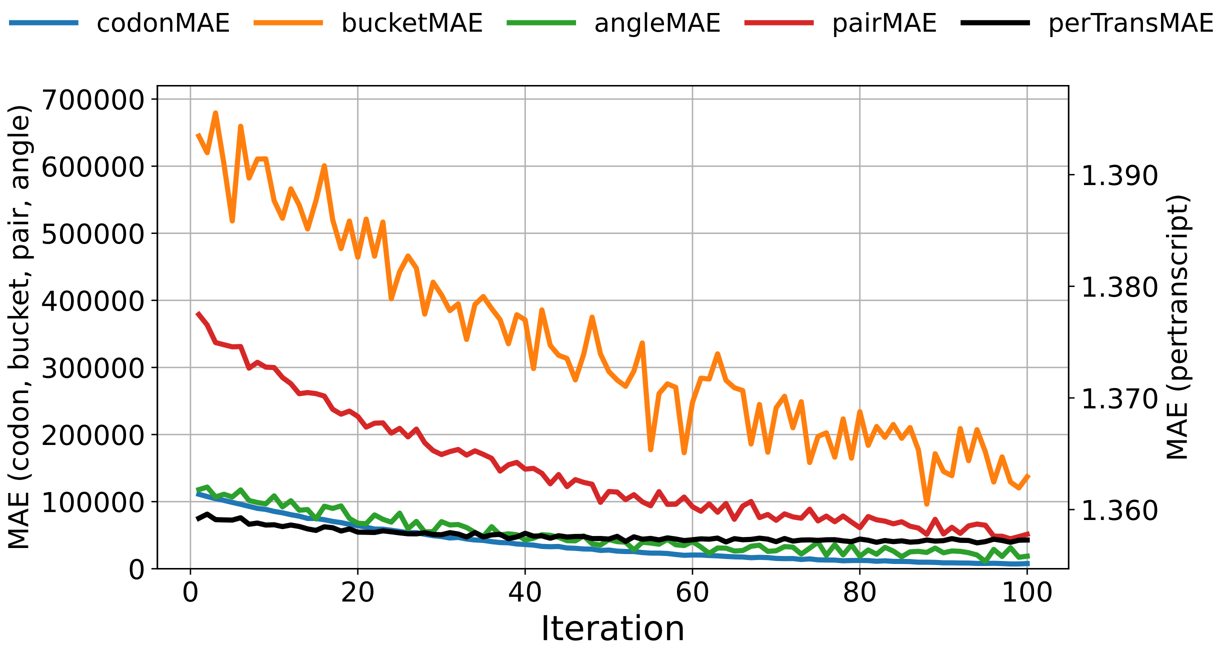


**Figure S25:** sTASEP training loss convergence for RPE-1.

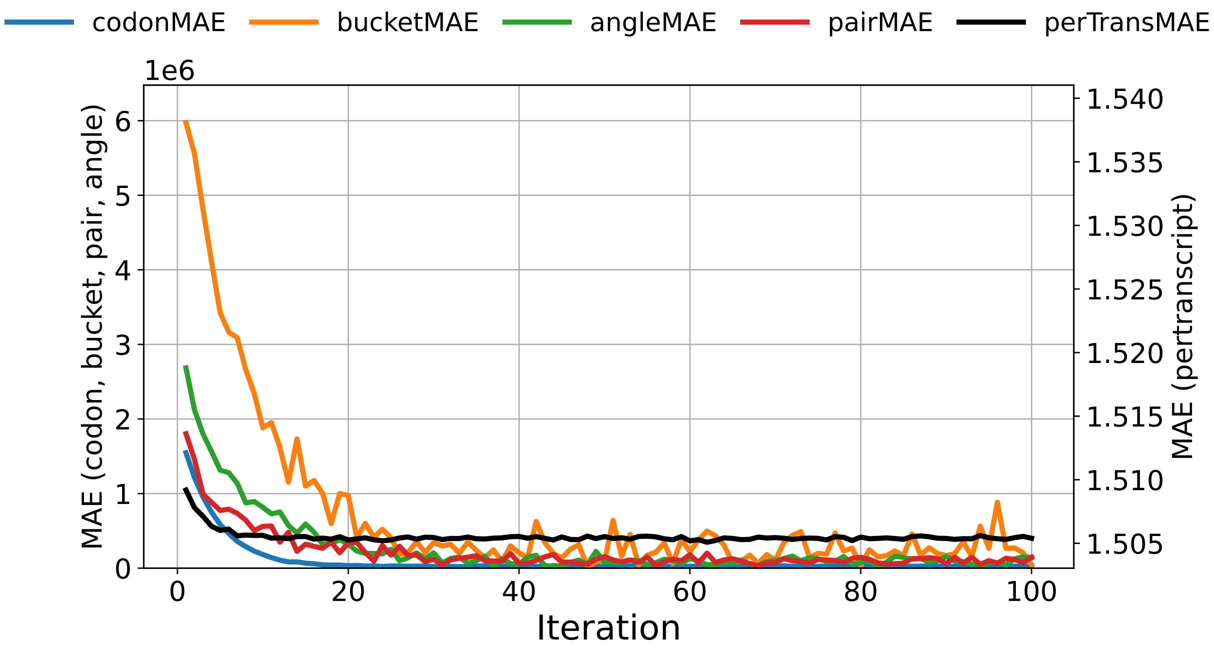


**Figure S26:** sTASEP training loss convergence for LCL.

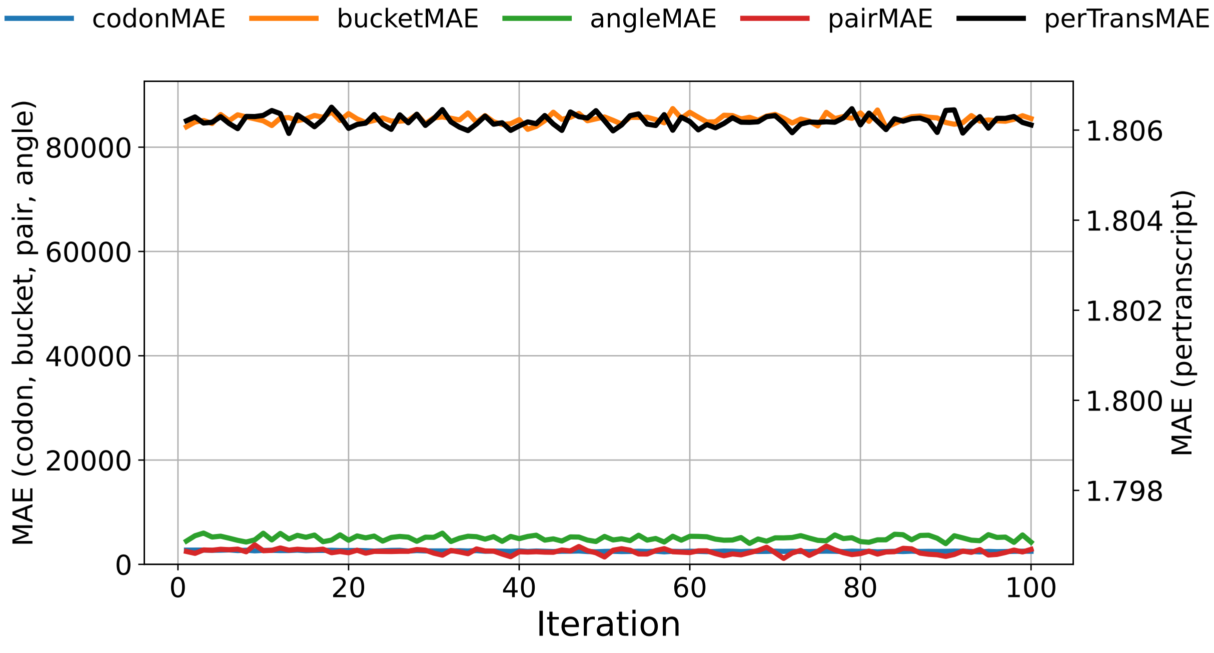


**Figure S27:** Classical TASEP training loss convergence for HEK293.

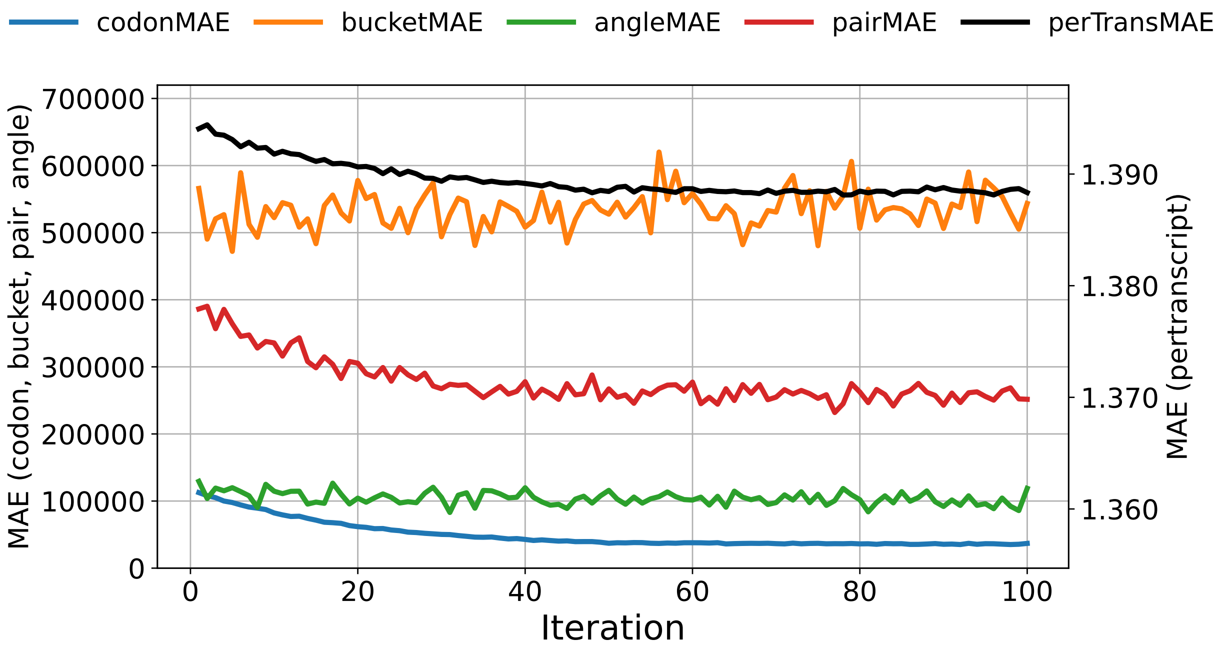


**Figure S28:** Classical TASEP training loss convergence for RPE-1.

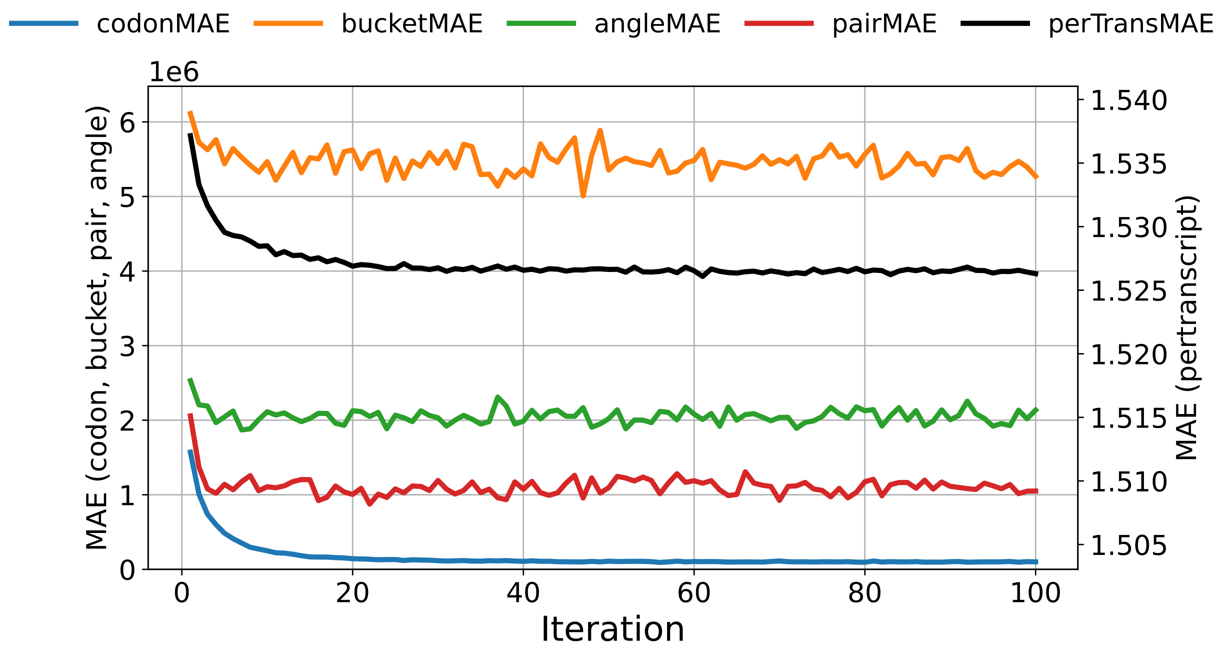


**Figure S29:** Classical TASEP training loss convergence for LCL.

##### 3.2 Validation Loss Curves for Simulation Methods

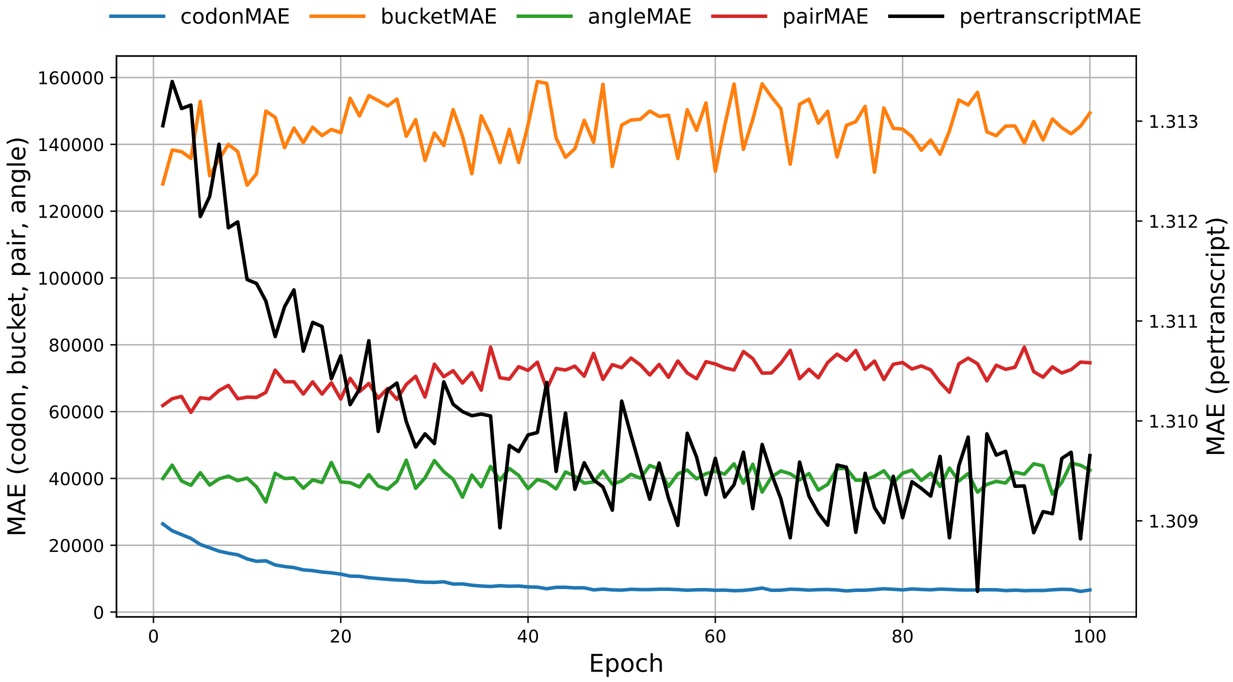


**Figure S30:** Loss convergence plot of validation set for classical TASEP model fit on iPSC.

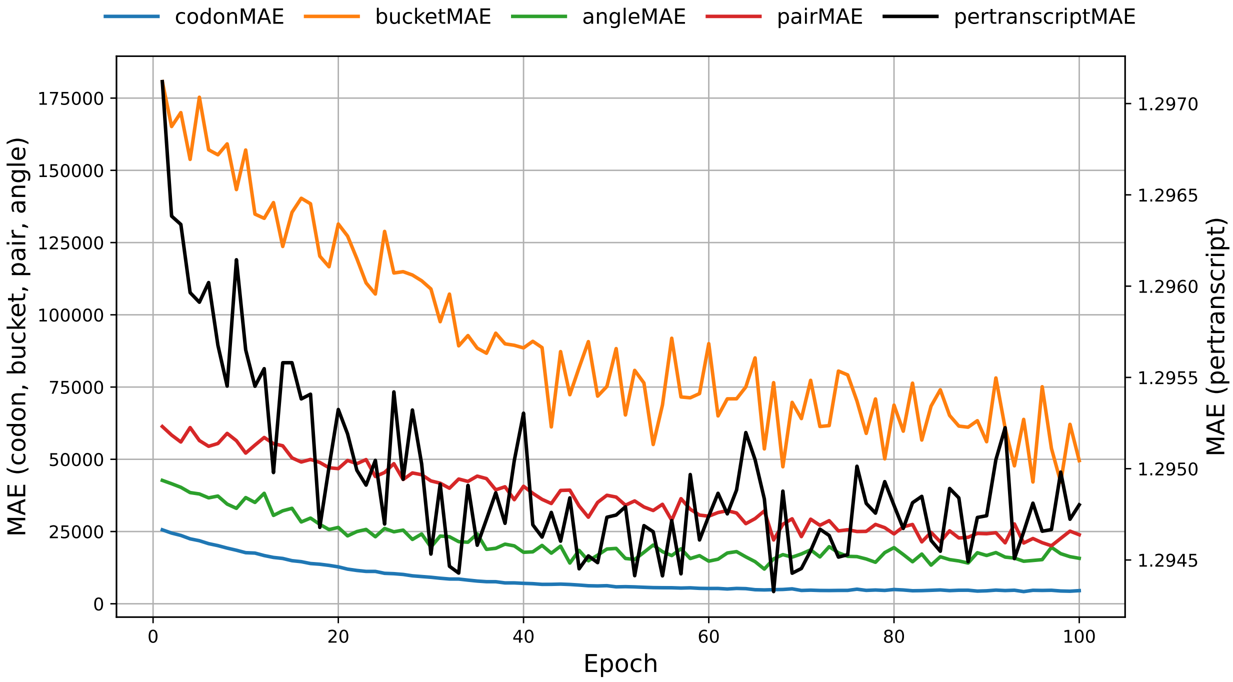


**Figure S31:** Loss convergence plot of validation set for sTASEP model fit on iPSC.

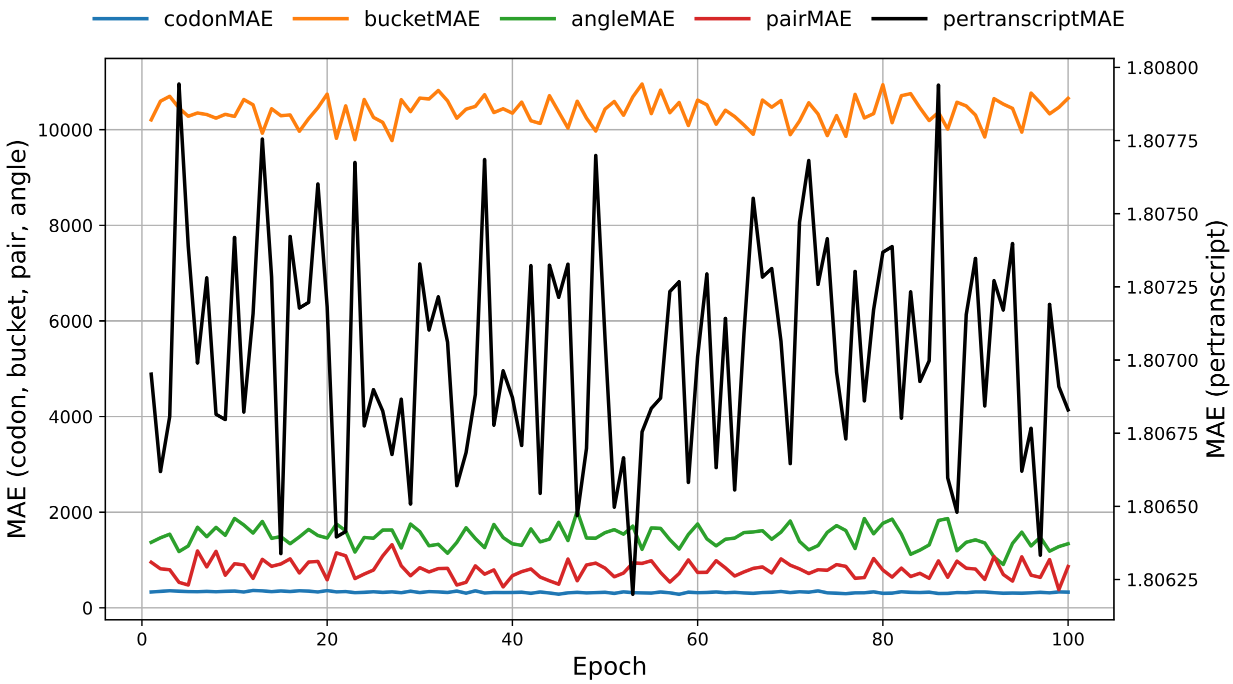


**Figure S32:** Loss convergence plot of validation set for classical TASEP model fit on HEK293.

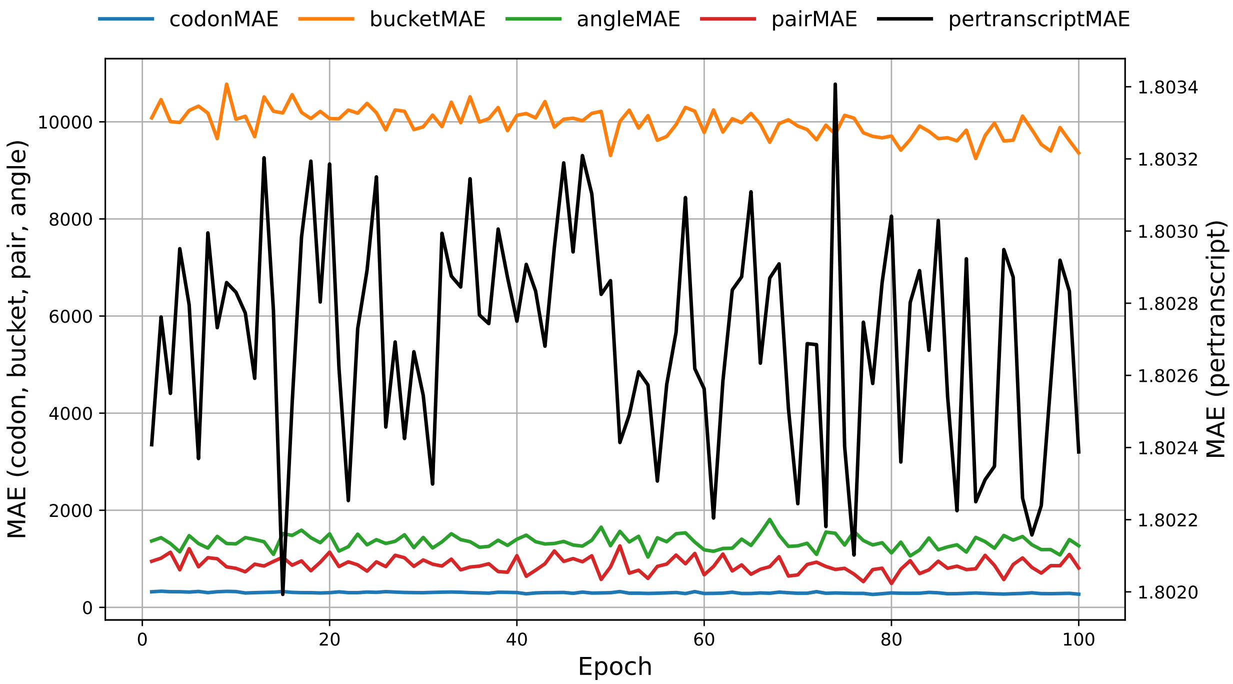


**Figure S33:** Loss convergence plot of validation set for sTASEP model fit on HEK293.

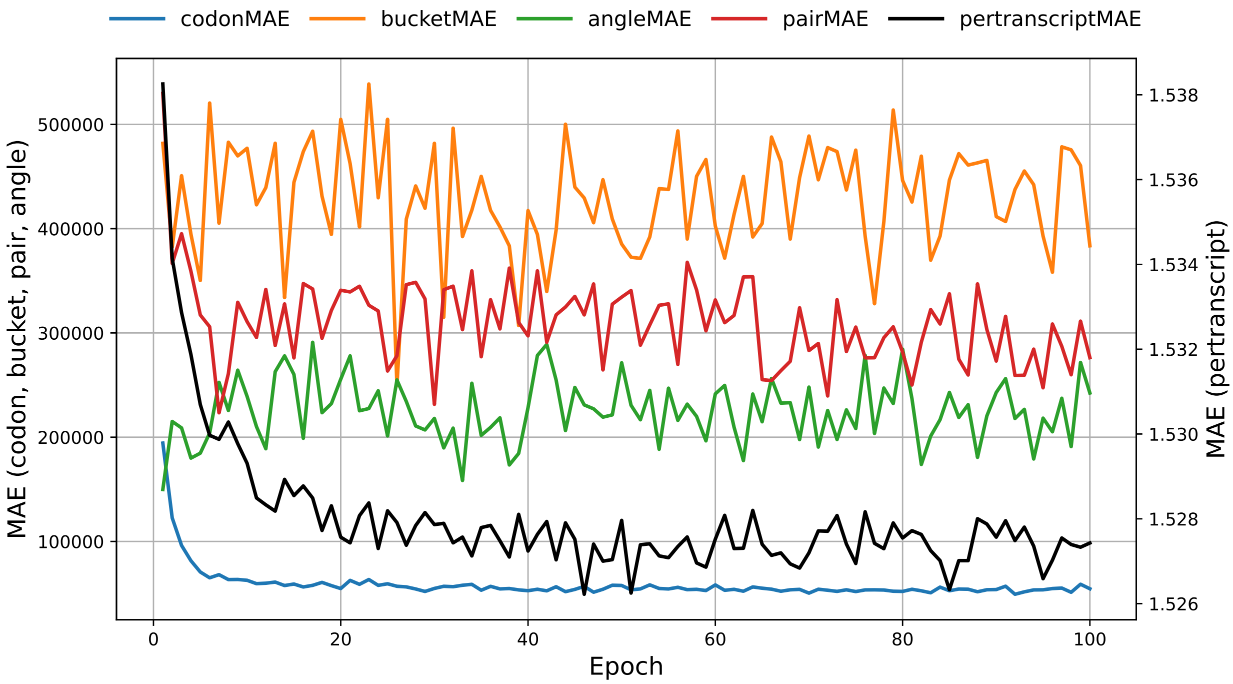


**Figure S34:** Loss convergence plot of validation set for classical TASEP model fit on LCL.

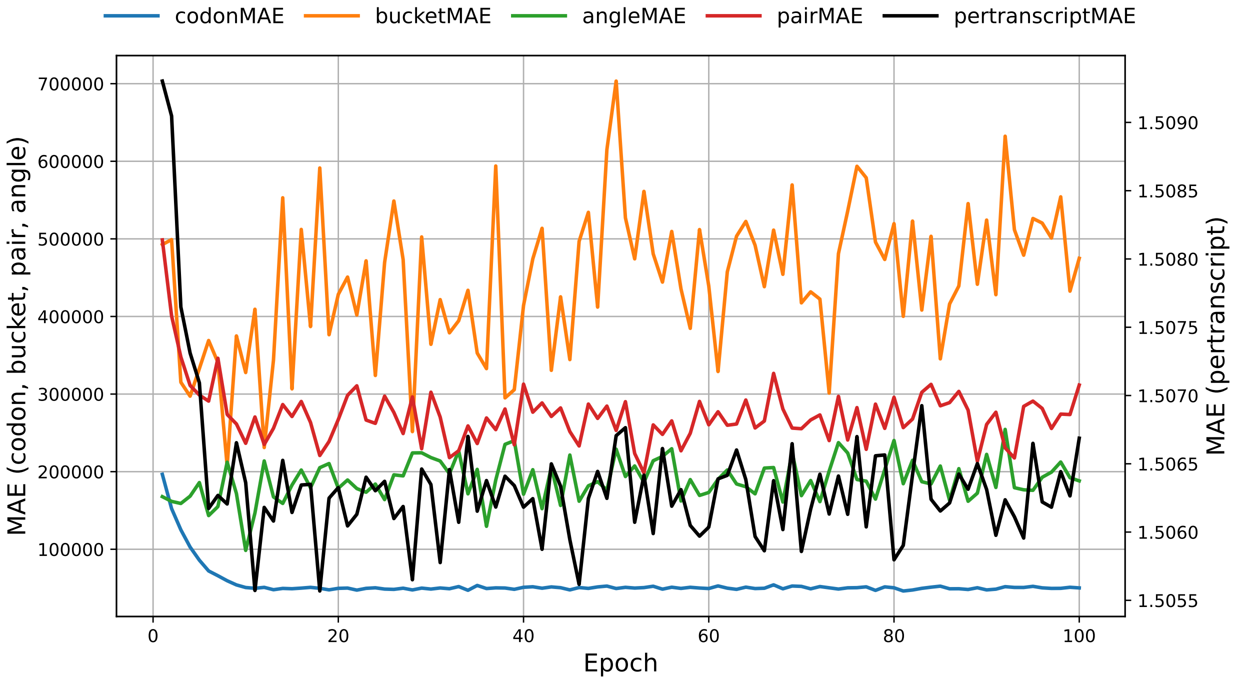


**Figure S35:** Loss convergence plot of validation set for sTASEP model fit on LCL.

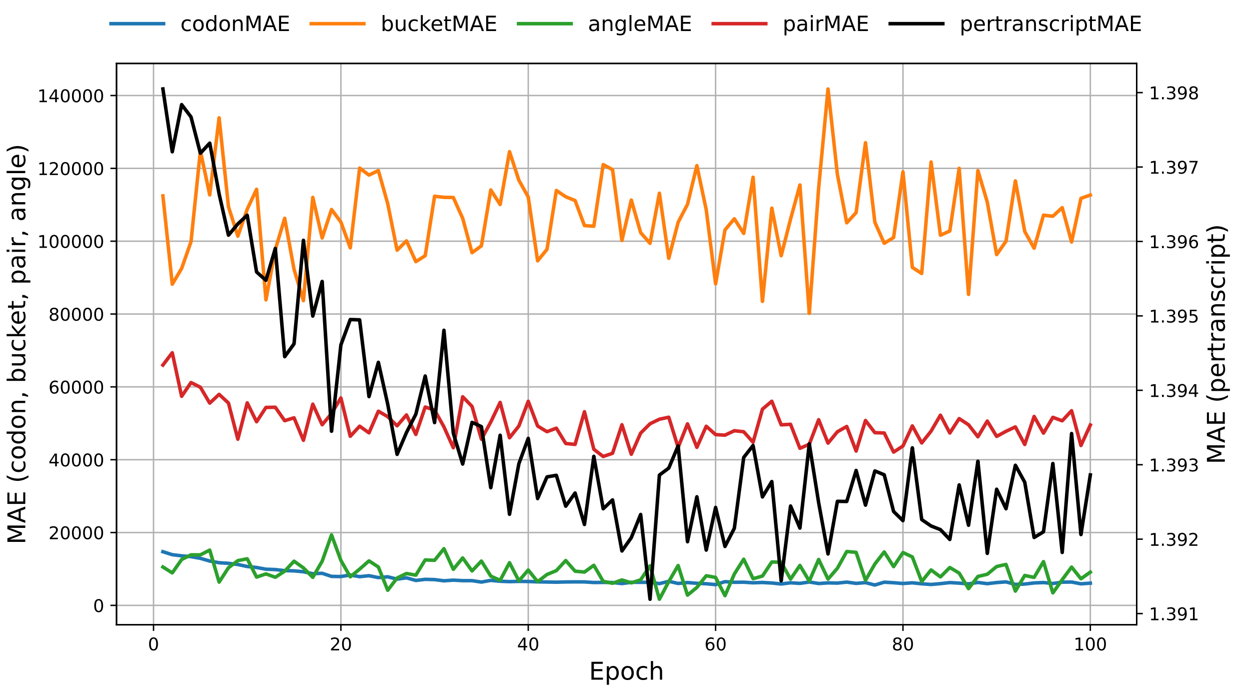


**Figure S36:** Loss convergence plot of validation set for classical TASEP model fit on RPE-1.

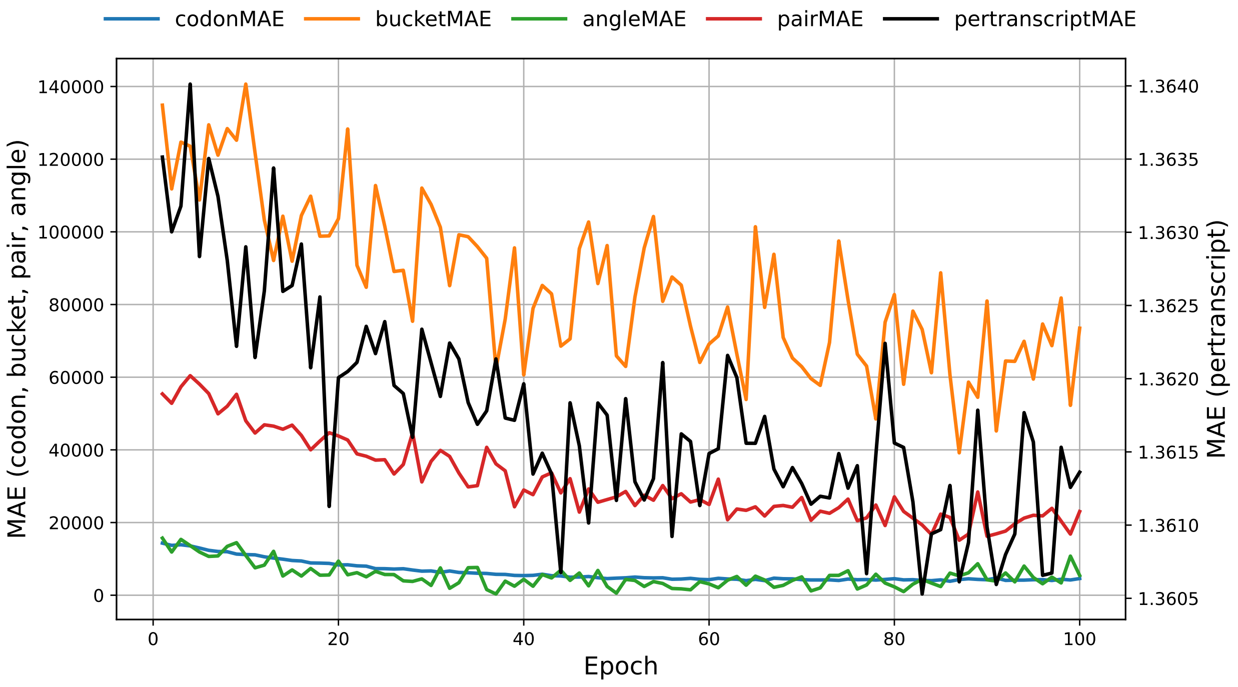


**Figure S37:** Loss convergence plot of validation set for sTASEP model fit on RPE-1.

##### 3.3 Training and Validation Loss Curves for Polisher Training

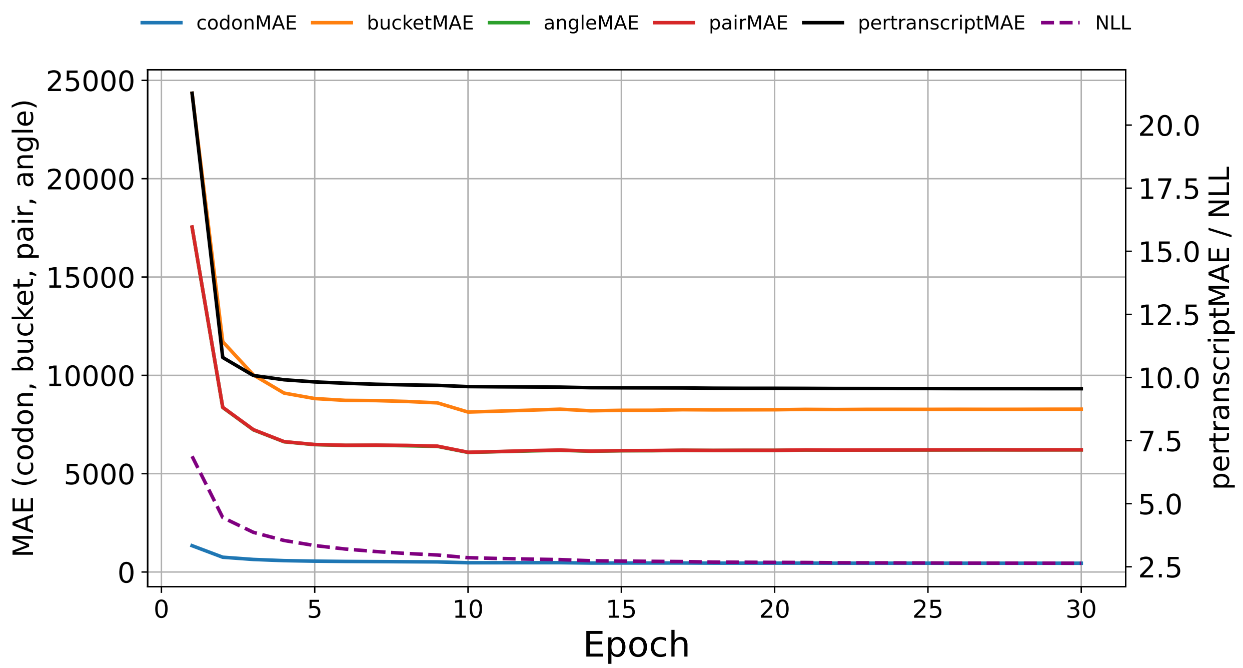


**Figure S38:** Training loss convergence during training of the `seq2ribo` polisher (iPSC).

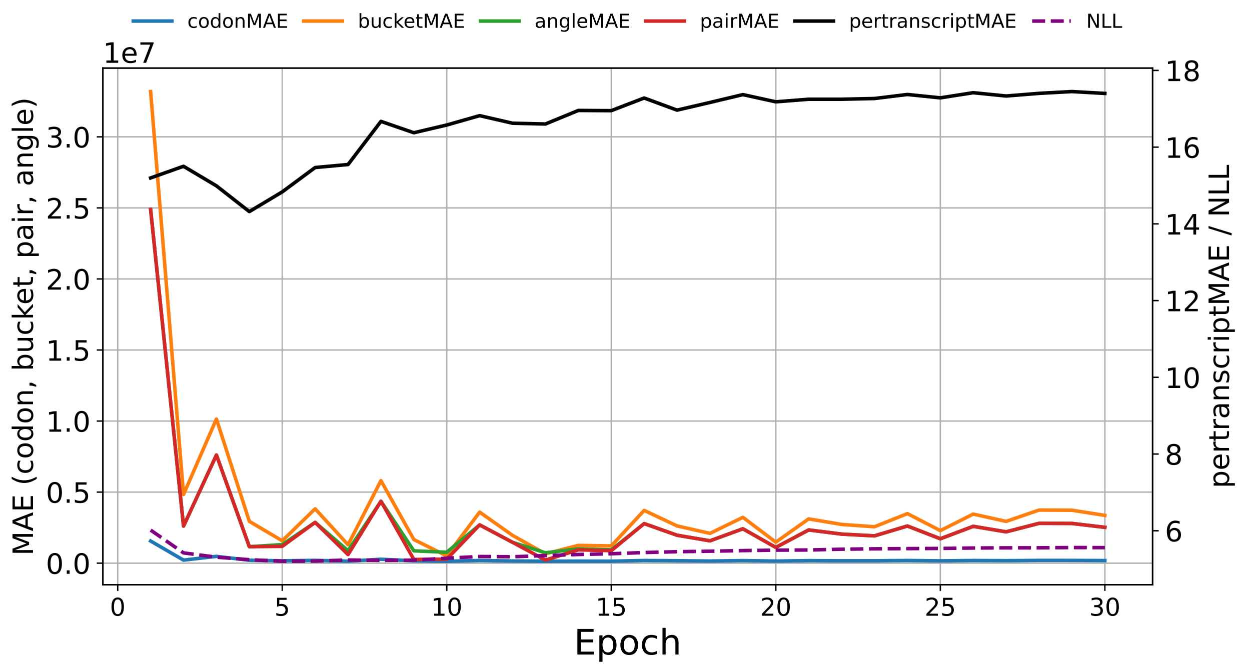


**Figure S39:** Validation loss convergence during training of the `seq2ribo` polisher (iPSC).

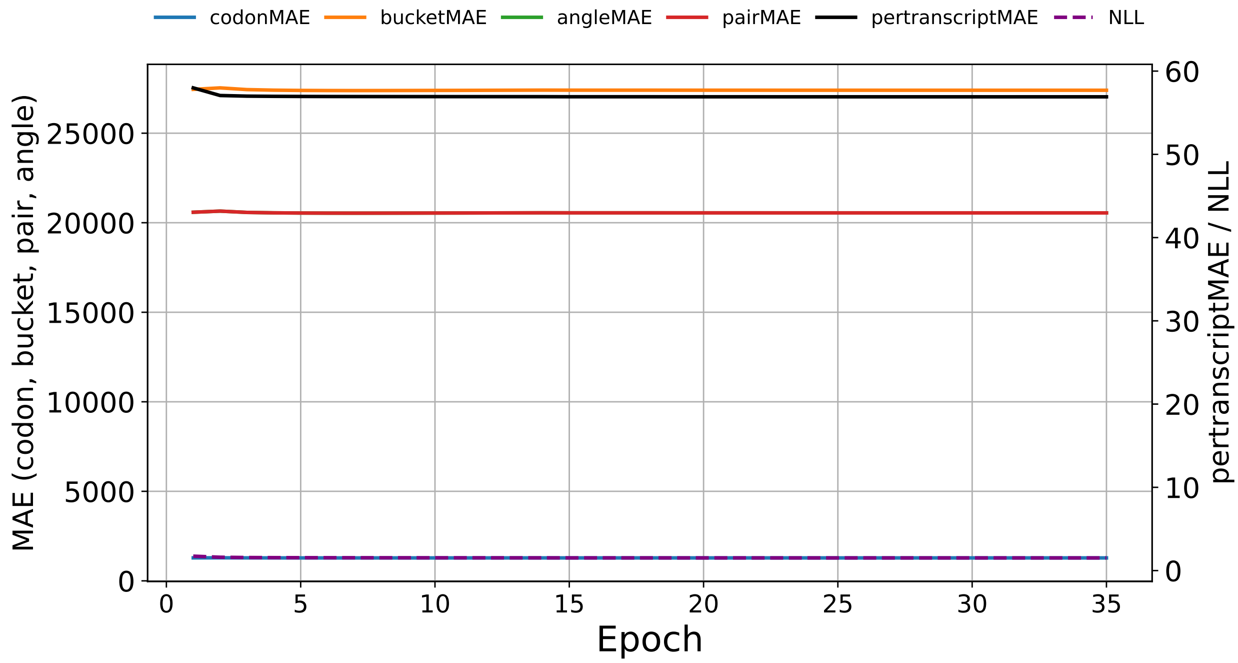


**Figure S40:** Training loss convergence during training of the `seq2ribo` polisher (HEK293).

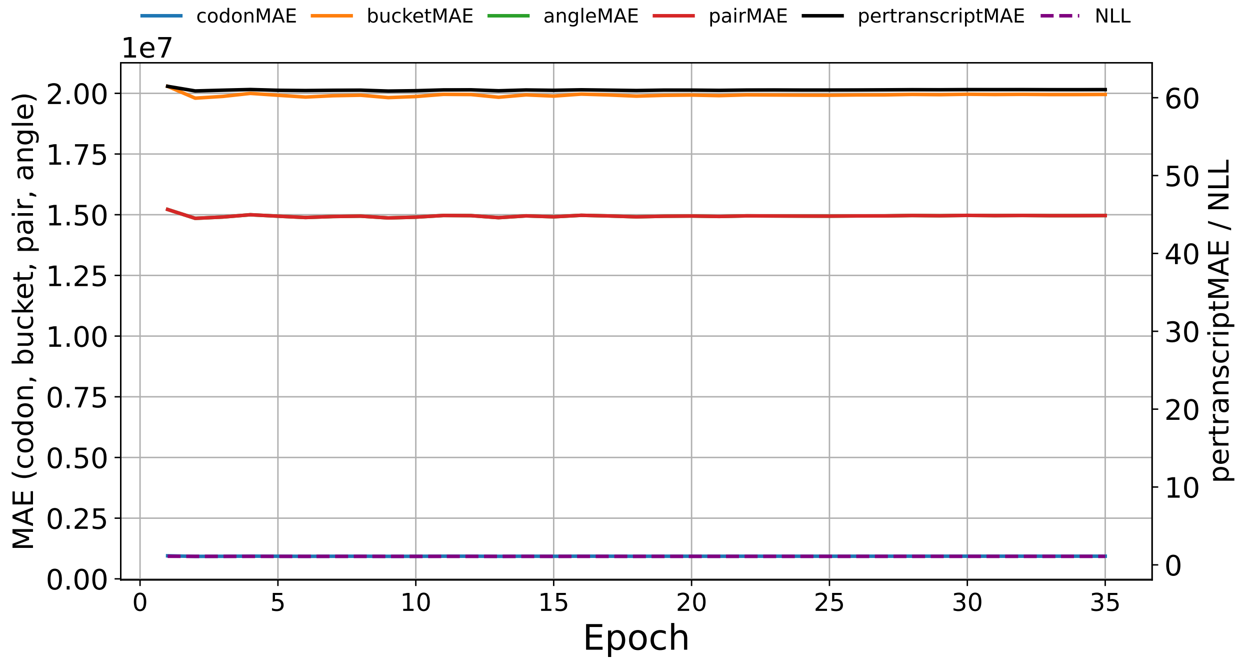


**Figure S41:** Validation loss convergence during training of the `seq2ribo` polisher (HEK293).

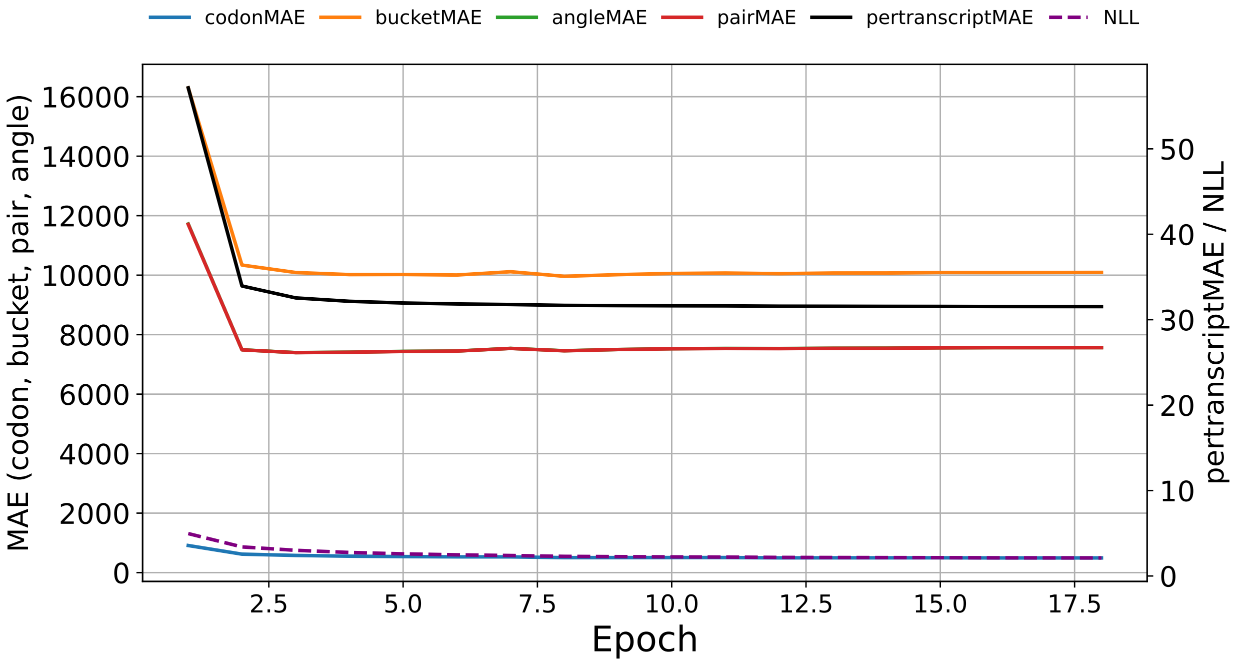


**Figure S42:** Training loss convergence during training of the `seq2ribo` polisher (RPE-1).

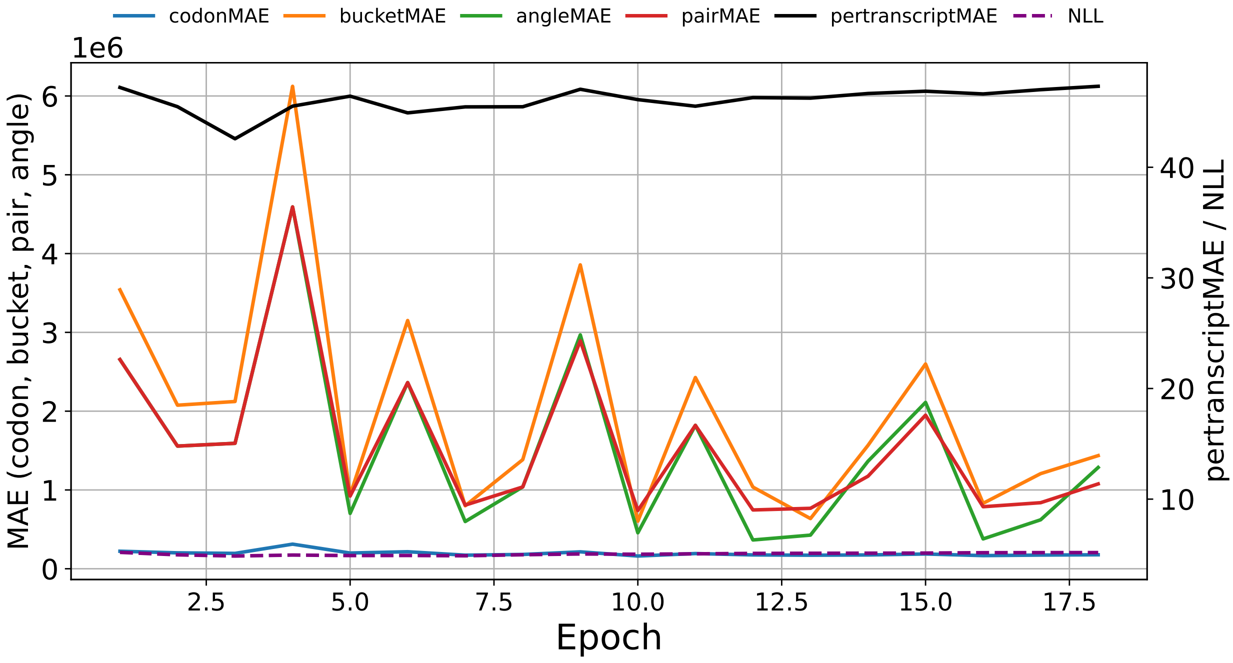


**Figure S43:** Validation loss convergence during training of the `seq2ribo` polisher (RPE-1).

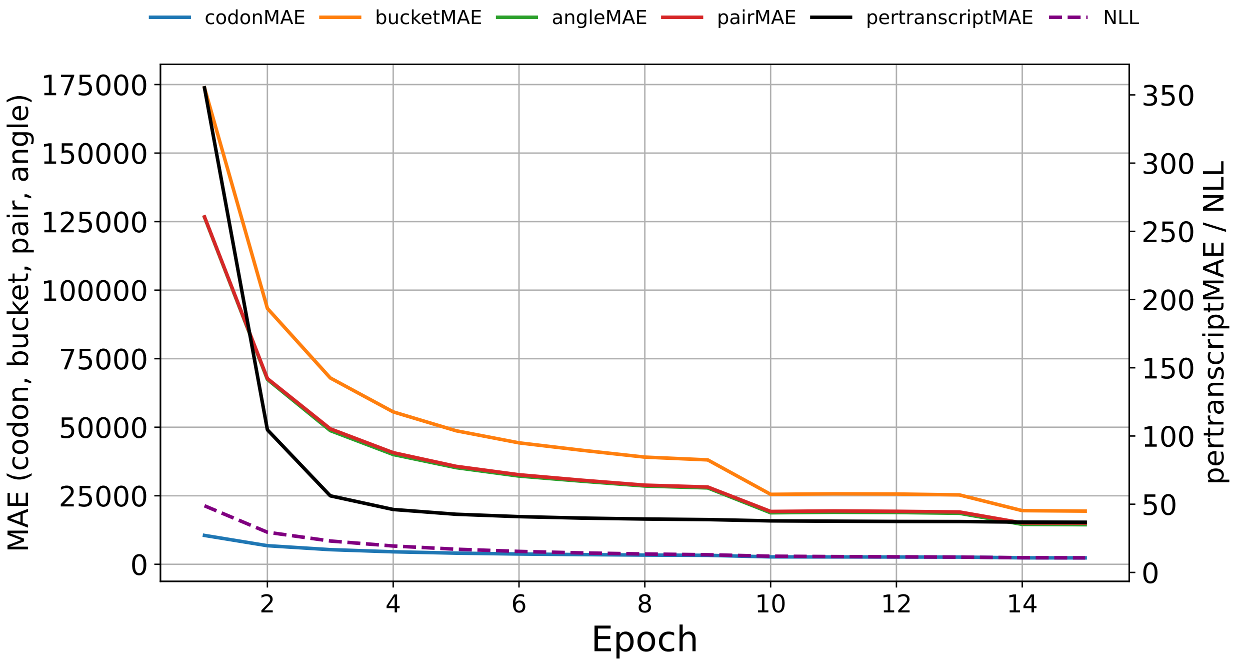


**Figure S44:** Training loss convergence during training of the `seq2ribo` polisher (LCL).

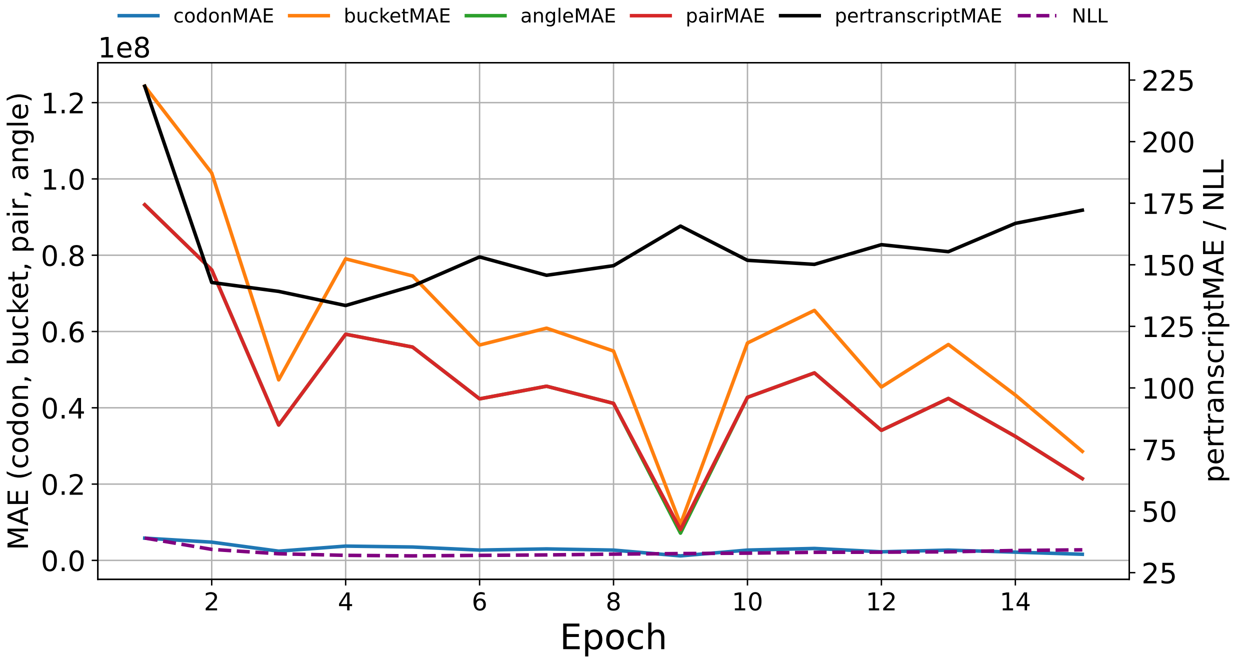


**Figure S45:** Validation loss convergence during training of the `seq2ribo` polisher (LCL).

#### 3.4 Translation Efficiency `seq2ribo` Scatter Plots

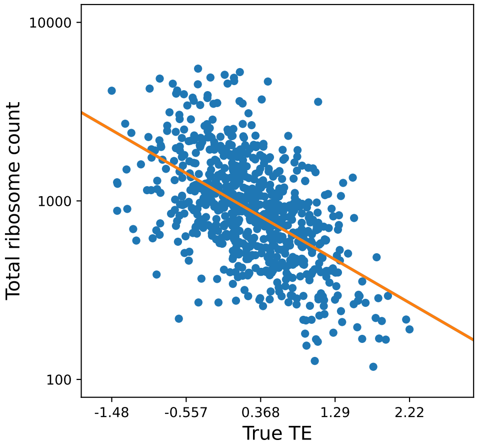


**Figure S46:** Correlation between total predicted ribosome load from the fixed `seq2ribo` polisher and experimental translation efficiency on the iPSC test set. The y-axis is displayed in log-transformed space for visualization.

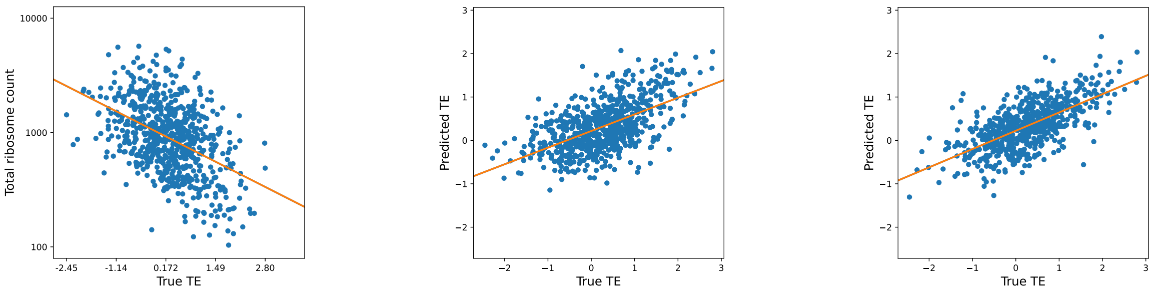


**(a)** Total ribosome load vs. experimental TE

**(b)** Finetuned `seq2ribo` (CDS only)

**(c)** Finetuned `seq2ribo` (CDS + UTR)

**Figure S47:** Translation efficiency prediction on the HEK293 test set. (a) Correlation between total predicted ribosome load from the fixed polisher and experimental TE (y-axis in log-transformed space for visualization). (b) Finetuned `seq2ribo` predictions using CDS-only input. (c) Finetuned `seq2ribo` predictions using CDS + UTR input.

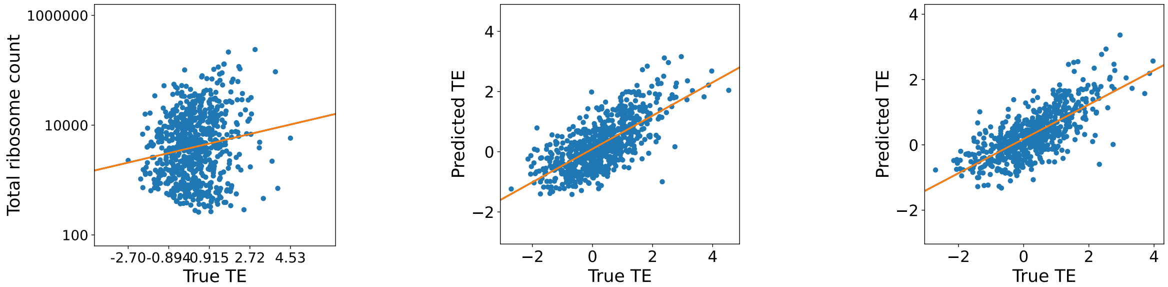


**(a)** Total ribosome load vs. experimental TE  **(b)** Finetuned `seq2ribo` (CDS only)  **(c)** Finetuned `seq2ribo` (CDS + UTR)

**Figure S48:** Translation efficiency prediction on the LCL test set. (a) Correlation between total predicted ribosome load from the fixed polisher and experimental TE (y-axis in log-transformed space for visualization). (b) Finetuned `seq2ribo` predictions using CDS-only input. (c) Finetuned `seq2ribo` predictions using CDS + UTR input. See Figure S49 for the corresponding RPE-1 results.

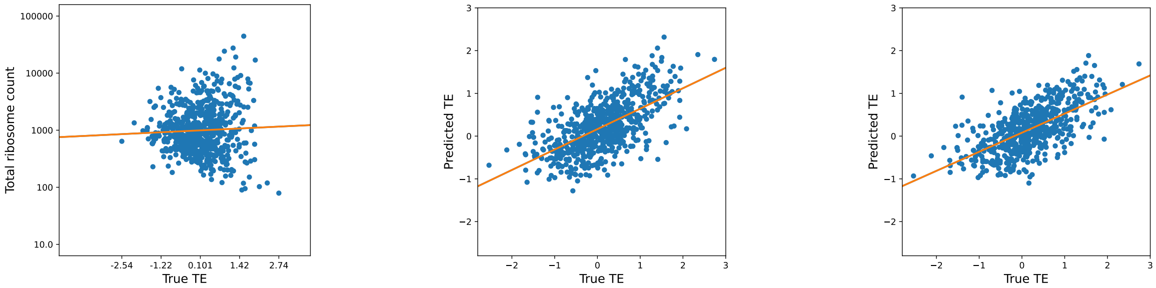


**(a)** Total ribosome load vs. experimental TE  **(b)** Finetuned `seq2ribo` (CDS only)  **(c)** Finetuned `seq2ribo` (CDS + UTR)

**Figure S49:** Translation efficiency prediction on the RPE-1 test set. (a) Correlation between total predicted ribosome load from the fixed polisher and experimental TE (y-axis in log-transformed space for visualization). (b) Finetuned `seq2ribo` predictions using CDS-only input. (c) Finetuned `seq2ribo` predictions using CDS + UTR input.

#### 3.5 Translation Efficiency Baseline Scatter Plots

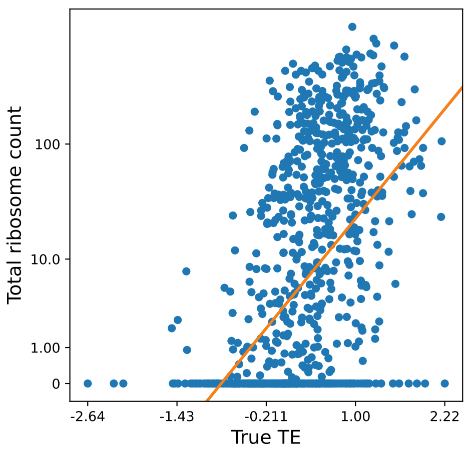


**Figure S50:** Translatomer (sequence-only) predictions vs. experimental translation efficiency on the iPSC test set.

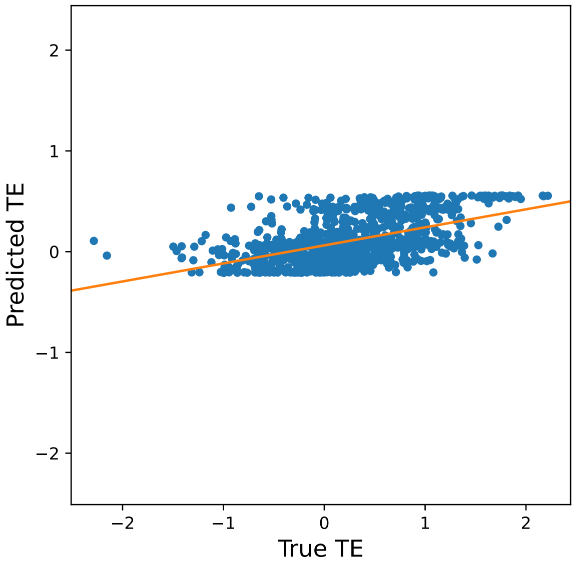


**(a)** Individual, CDS only

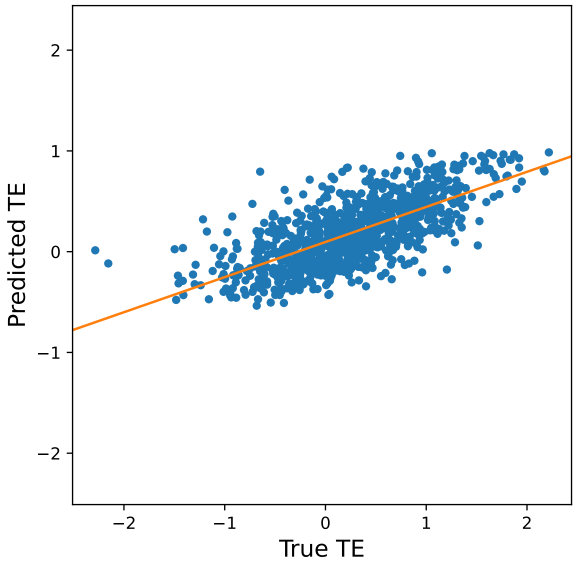


**(b)** Individual, CDS + UTR

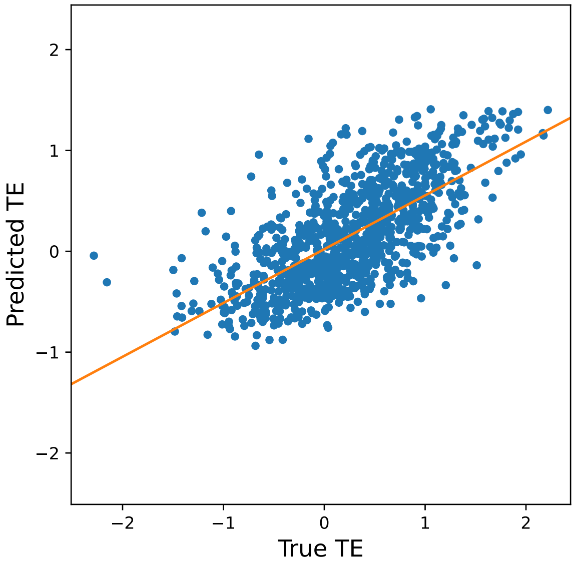


**(c)** Multitask, CDS only

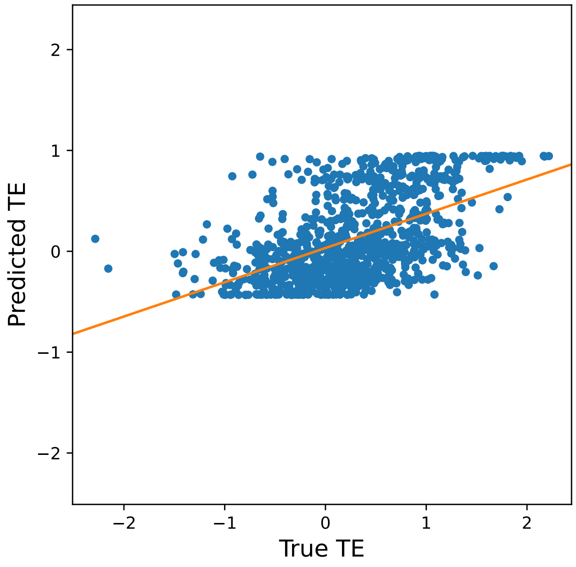


**(d)** Multitask, CDS + UTR

**Figure S51:** RiboNN predictions vs. experimental translation efficiency on the iPSC test set. Top row: independently trained models. Bottom row: multitask models. Left column: CDS-only input. Right column: CDS + UTR input.

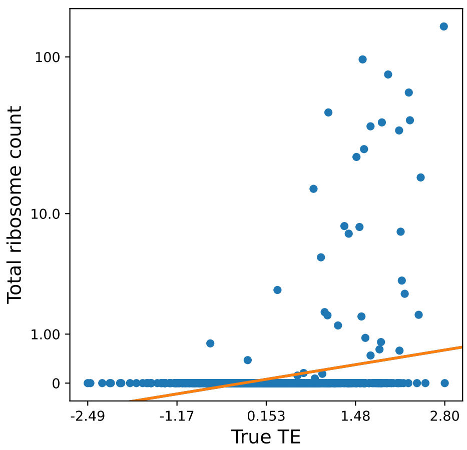


**Figure S52:** Translatomer (sequence-only) predictions vs. experimental translation efficiency on the HEK293 test set.

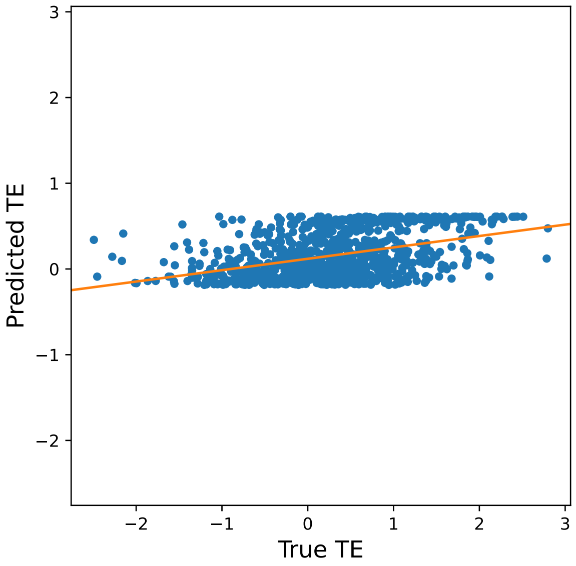


**(a)** Individual, CDS only

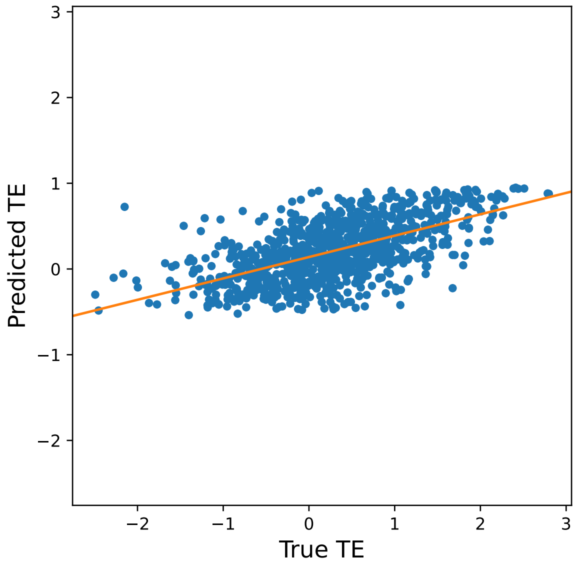


**(b)** Individual, CDS + UTR

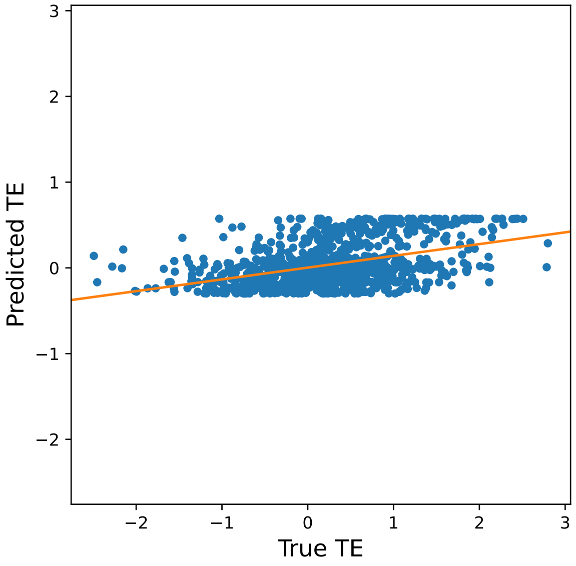


**(c)** Multitask, CDS only

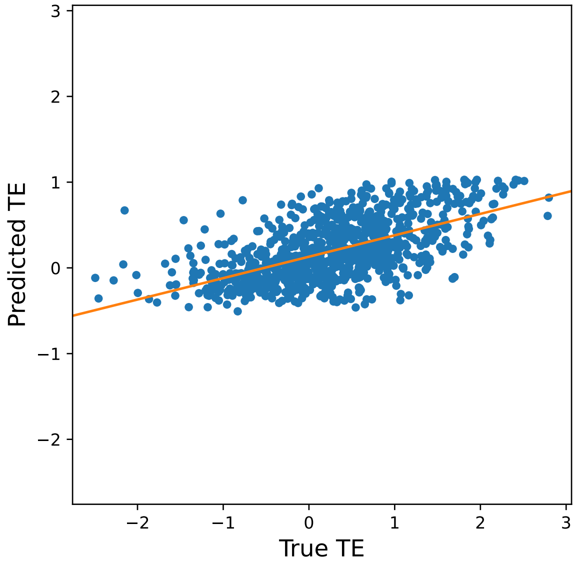


**(d)** Multitask, CDS + UTR

**Figure S53:** RiboNN predictions vs. experimental translation efficiency on the HEK293 test set. Top row: independently trained models. Bottom row: multitask models. Left column: CDS-only input. Right column: CDS + UTR input.

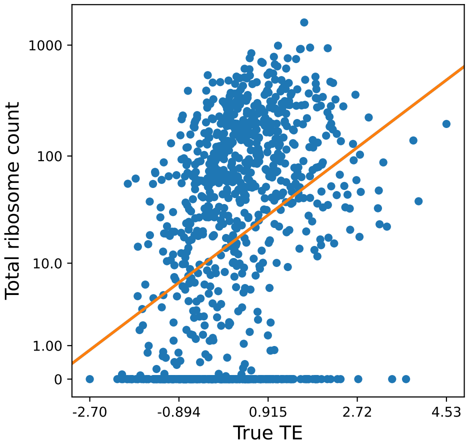


**Figure S54:** Translatomer (sequence-only) predictions vs. experimental translation efficiency on the LCL test set.

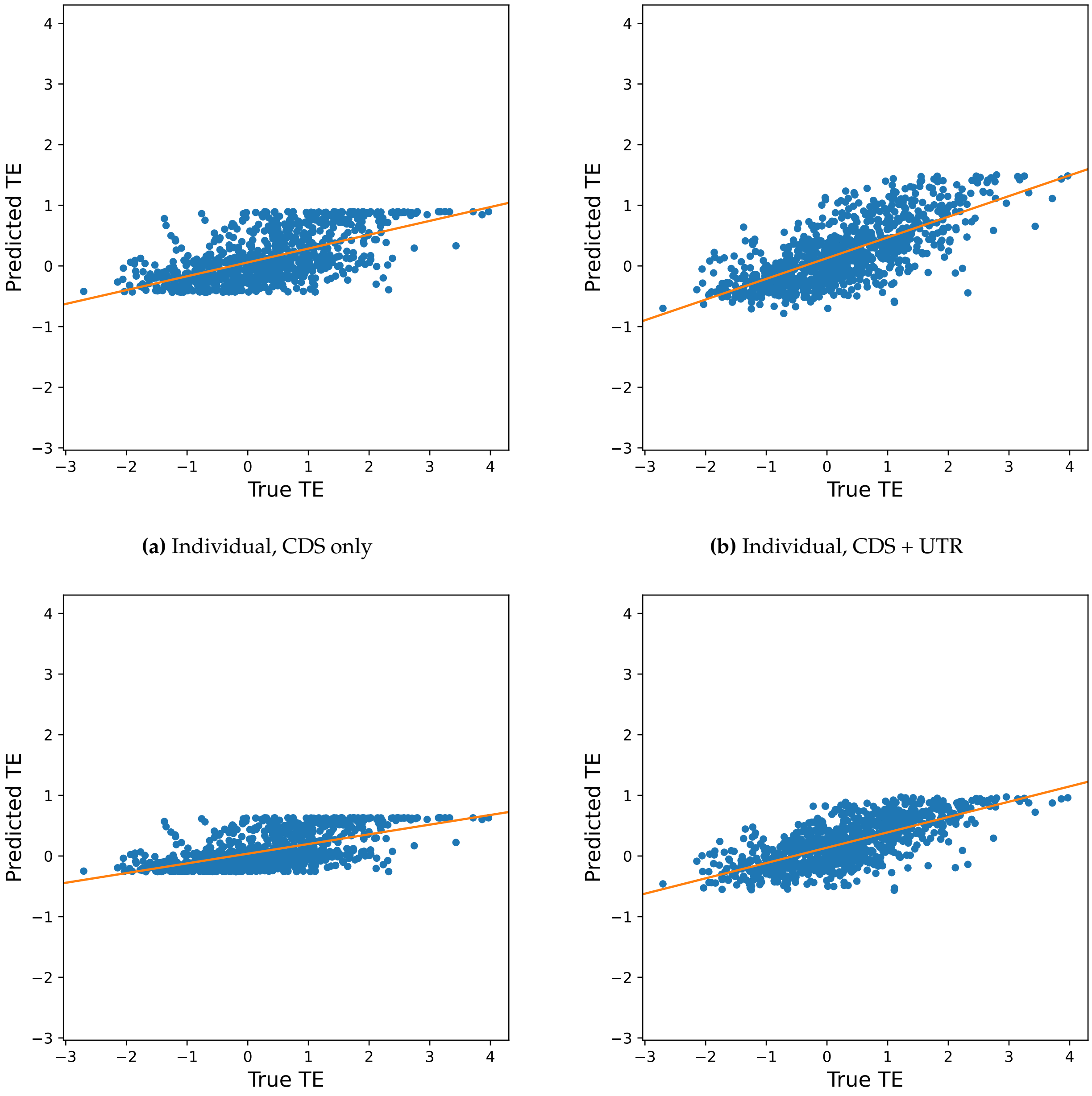


**(a)** Individual, CDS only

**(b)** Individual, CDS + UTR

**(c)** Multitask, CDS only

**(d)** Multitask, CDS + UTR

**Figure S55:** RiboNN predictions vs. experimental translation efficiency on the LCL test set. Top row: independently trained models. Bottom row: multitask models. Left column: CDS-only input. Right column: CDS + UTR input.

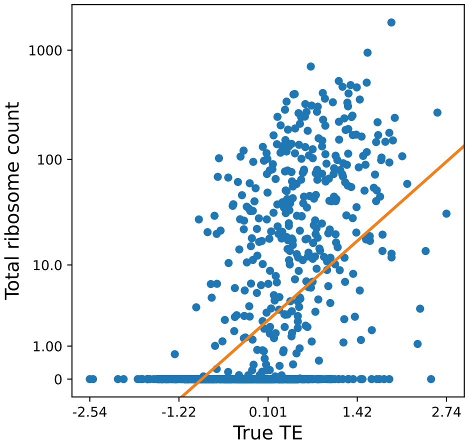


**Figure S56:** Translatomer (sequence-only) predictions vs. experimental translation efficiency on the RPE-1 test set.

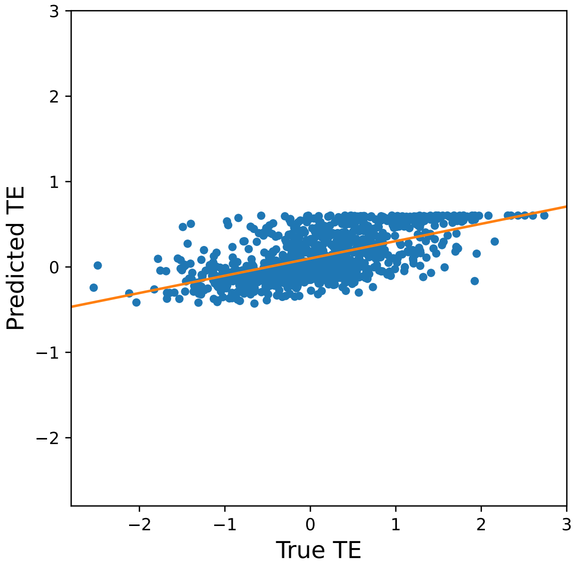


**(a)** Individual, CDS only

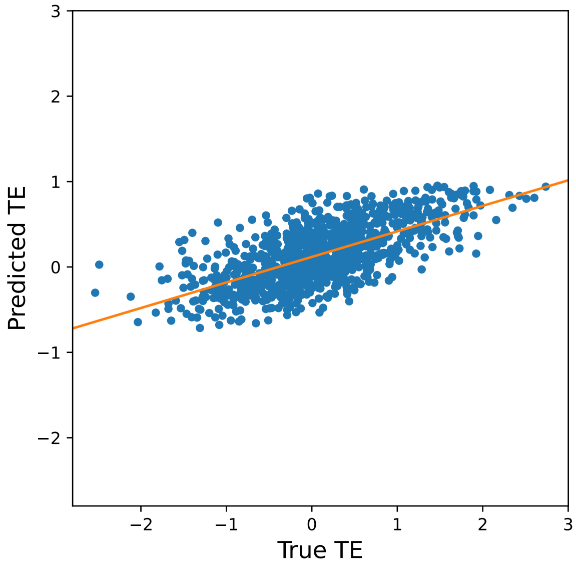


**(b)** Individual, CDS + UTR

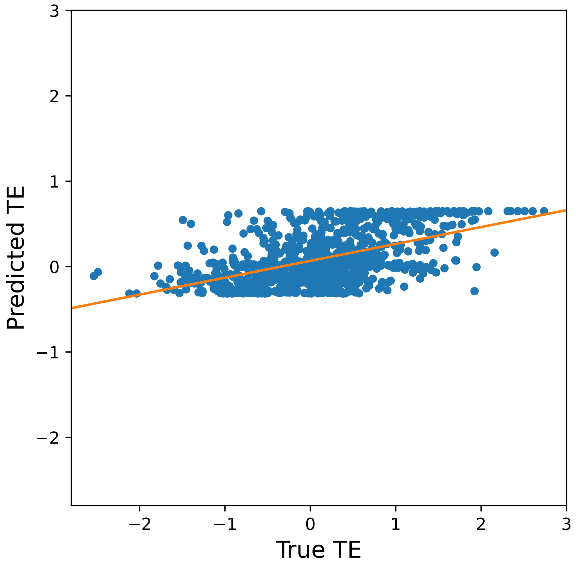


**(c)** Multitask, CDS only

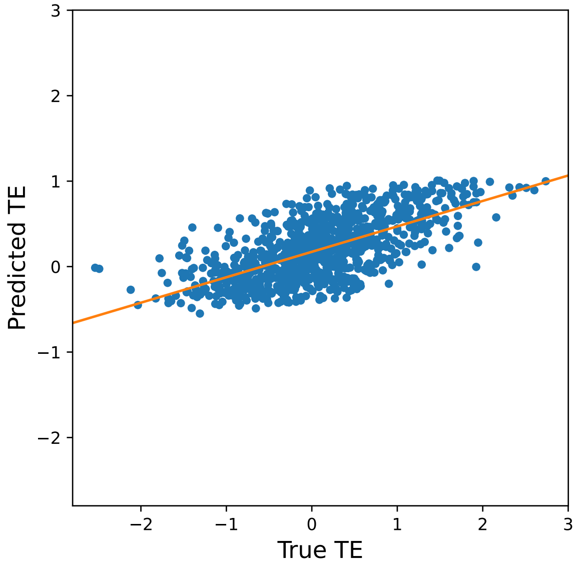


**(d)** Multitask, CDS + UTR

**Figure S57:** RiboNN predictions vs. experimental translation efficiency on the RPE-1 test set. Top row: independently trained models. Bottom row: multitask models. Left column: CDS-only input. Right column: CDS + UTR input.

#### 3.6 Protein Expression `seq2ribo` Scatter Plots

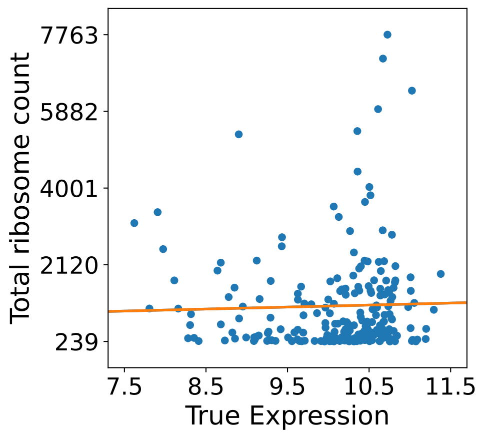


**(a)** iPSC polisher

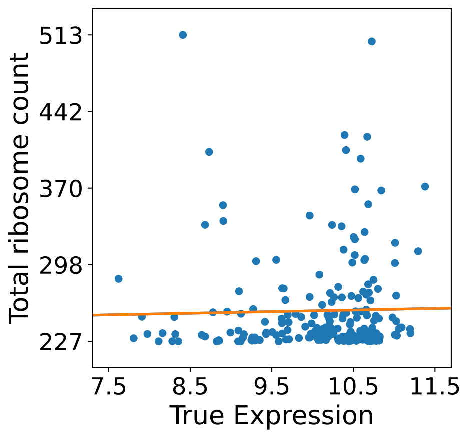


**(b)** HEK293 polisher

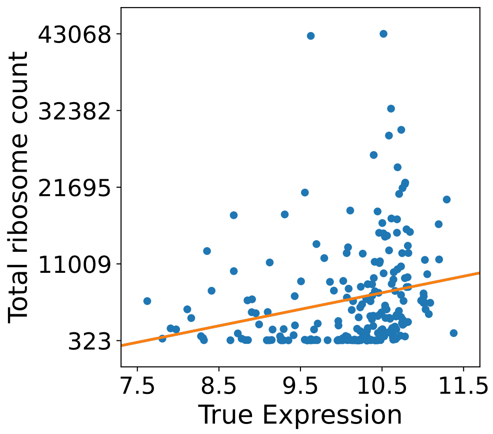


**(c)** LCL polisher

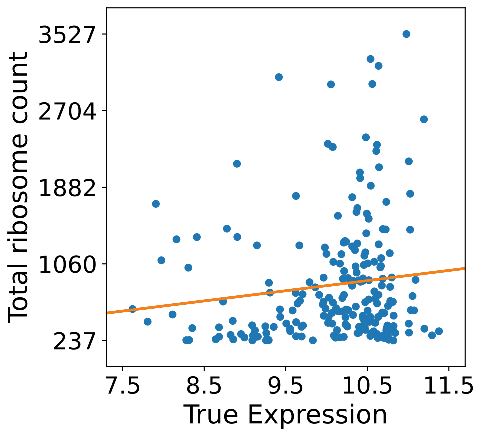


**(d)** RPE-1 polisher

**Figure S58:** Correlation between total predicted ribosome load from the fixed `seq2ribo` polisher and measured protein expression on the mRFP test set, for each cell-line-specific polisher. No expression-specific finetuning is applied; the total ribosome count per transcript is used as the predictor.

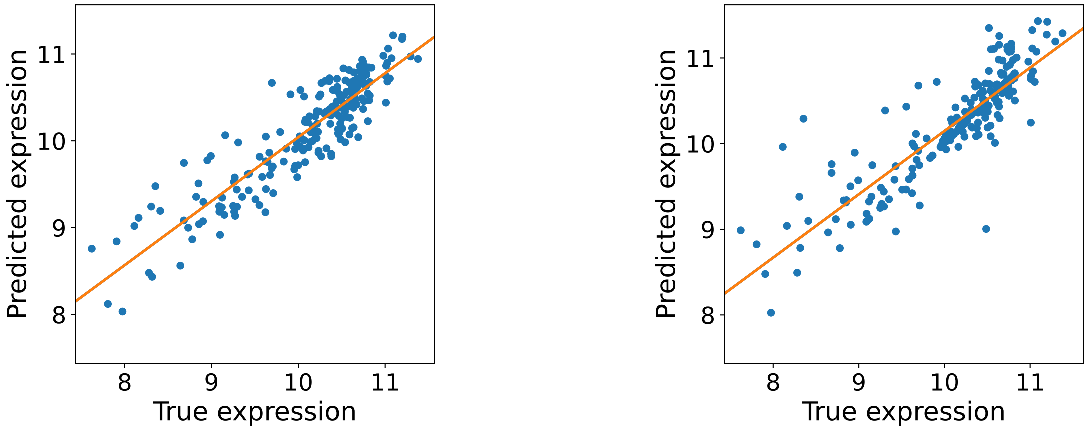


**(a)** HEK293 polisher (Pearson $r = 0.883$)

**(b)** RPE-1 polisher (Pearson $r = 0.830$)

**Figure S59:** Correlation of finetuned `seq2ribo` predictions with measured protein expression on the mRFP test set for HEK293 and RPE-1 polishers. Corresponding scatter plots for the iPSC and LCL polishers appear in the main text.

#### 3.7 Protein Expression Baseline Scatter Plots

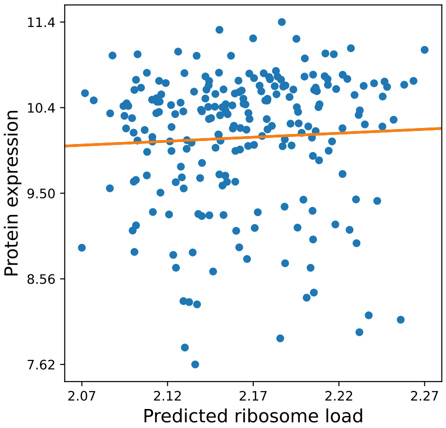


**(a)** iPSC

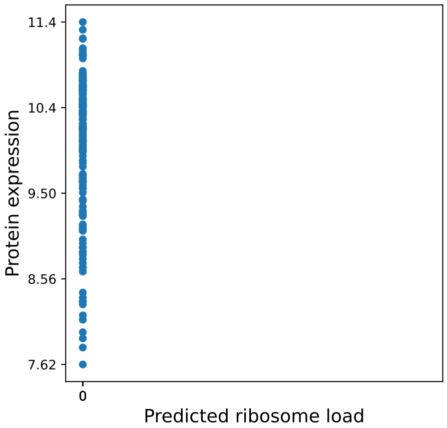


**(b)** HEK293

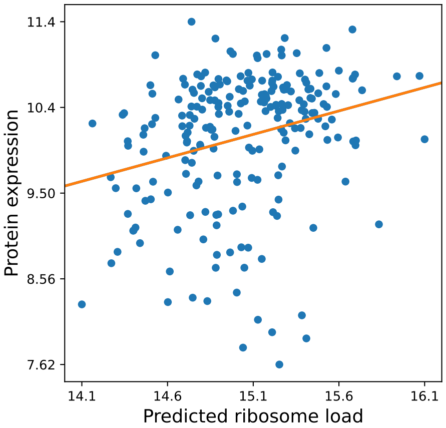


**(c)** LCL

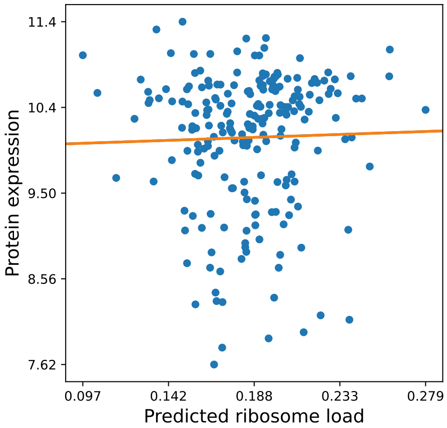


**(d)** RPE-1

**Figure S60:** Translatomer (sequence-only) predictions vs. measured protein expression on the mRFP test set, for each cell-line-specific model.
